# High yield, low magnesium flexizyme reactions in a water-ice eutectic phase

**DOI:** 10.1101/2023.12.03.569792

**Authors:** Joshua A. Davisson, Jose L. Alejo, Mace Blank, Evan M. Kalb, Angelin Prasad, Isaac J. Knudson, Alanna Schepartz, Aaron E. Engelhart, Katarzyna P. Adamala

## Abstract

Flexizymes enable the stoichiometric acylation of tRNAs with a variety of compounds, enabling the *in vitro* translation of peptides with both non-natural backbones and side chains. However, flexizyme reactions have several drawbacks, including single-turnover kinetics, high Mg(II) carryover inhibiting *in vitro* translation, and rapid product hydrolysis. Here we present flexizyme reactions utilizing an ice-eutectic phase, with high yields, 30X lower Mg(II), and long-term product stability. The eutectic flexizyme reactions increase the ease of use and flexibility of flexizyme aminoacylation, and increase the *in vitro* protein production.

## Introduction

Flexizymes have become ubiquitous tools for *in vitro* genetic code reprogramming, acylating an enormous range of chemically diverse molecules onto tRNA. Unlike aminoacyl tRNA synthetases, flexizymes can acylate a wide array of amino acids to tRNA (Katoh & Suga, 2022). Because of this, flexizymes have been used to radically reprogram the existing codon table to translate aramids (Ad et al., 2019), *N*-methyl amino acids (Kawakami et al., 2008), D-amino acids (Katoh, Tajima, et al., 2017), β-amino acids (Katoh & Suga, 2018), γ-amino acids (Ohshiro et al., 2011), pyridazinone oligomers (Lee et al., 2022), aromatic foldamers (Rogers et al., 2018), and aminobenzoic acid derivatives (Katoh & Suga, 2020). However, the established protocols for tRNA acylation using flexizyme present several genuine disadvantages: single-turnover kinetics, high Mg(II) carryover that lower the yields of in vitro translation, and low yields/slow reaction rates for many interesting substrates. In a flexizyme reaction, a 1:1 ratio of flexizyme and tRNA are essential because early studies of flexizyme kinetics demonstrated high levels of product inhibition (Murakami et al., 2003).

Furthermore, flexizyme reactions require Mg(II) concentrations as high as 600 mM, and the excess Mg(II) is challenging to remove and can dramatically inhibit subsequent in vitro translation reactions, especially when higher concentrations of acyl-tRNA are used. A recent paper demonstrating aaRS-free translation using tRNAs acylated with flexizyme resorted to HPLC purification of all acyl-tRNAs because the excess Mg(II) inhibited translation (Chen et al., 2021).

Taken together, these factors limit the utility of flexizymes for many applications. Since no widely used alternative currently exists to acylate tRNA with non-cognate amino acids (especially those with alternative backbones), we set out to engineer flexizyme reactions to circumvent those limiting factors.

Inspired by work using a water ice-eutectic phase for both enzymatic and chemical extension of nucleic acid chains (Attwater et al., 2010), (Monnard et al., 2003) we sought to apply the same concepts to flexizyme reaction conditions. We discovered we could use 30 fold lower magnesium concentrations than typical flexizyme reactions with similar or improved yields and by slowing hydrolysis we significantly extended the window of acceptable reaction times.

## Results and Discussion

Inspired by the water ice-eutectic phase used previously to facilitate an RNA polymerase ribozyme reaction (Attwater et al., 2010), we decided to test flexizyme reactions under eutectic conditions. Important for this previous work was the reduction of buffer components and salts in the reactions. Similarly, our eutectic phase reactions differ from standard flexizyme reactions by using reduced buffer, MgCl_2_, and DMSO concentrations in addition to using a lower reaction temperature. In our initial tests, flexizyme reactions were first frozen in liquid nitrogen, then incubated in an ethylene glycol-water bath to raise the temperature and initiate the reaction (**Figure 1a** and S73). We tested charging of activated amino acids L-Tyr-CME, L-Ala-DBE, and L-Leu-ABT with eFx, dFx, and aFx respectively to confirm eutectic phase reactions worked for all three flexizymes (**Figure 2a-d**). Furthermore, we screened varying amounts of magnesium to determine an ideal concentration (Figure S72). Charging efficiency was estimated by gel analysis of aminoacylated microhelix (**Fig. 2a-d**). In all cases, the observed yields under low magnesium eutectic conditions were similar to the corresponding reaction under the high magnesium standard flexizyme conditions. These eutectic phase reactions demonstrated that aminoacylation not only proceeds to high yields, but also requires significantly lower magnesium concentrations in an eutectic environment than in the standard reactions. Encouraged by those results, we proceeded to characterize the eutectic aminoacylation conditions with all three flexizymes.

**Figure 1.**
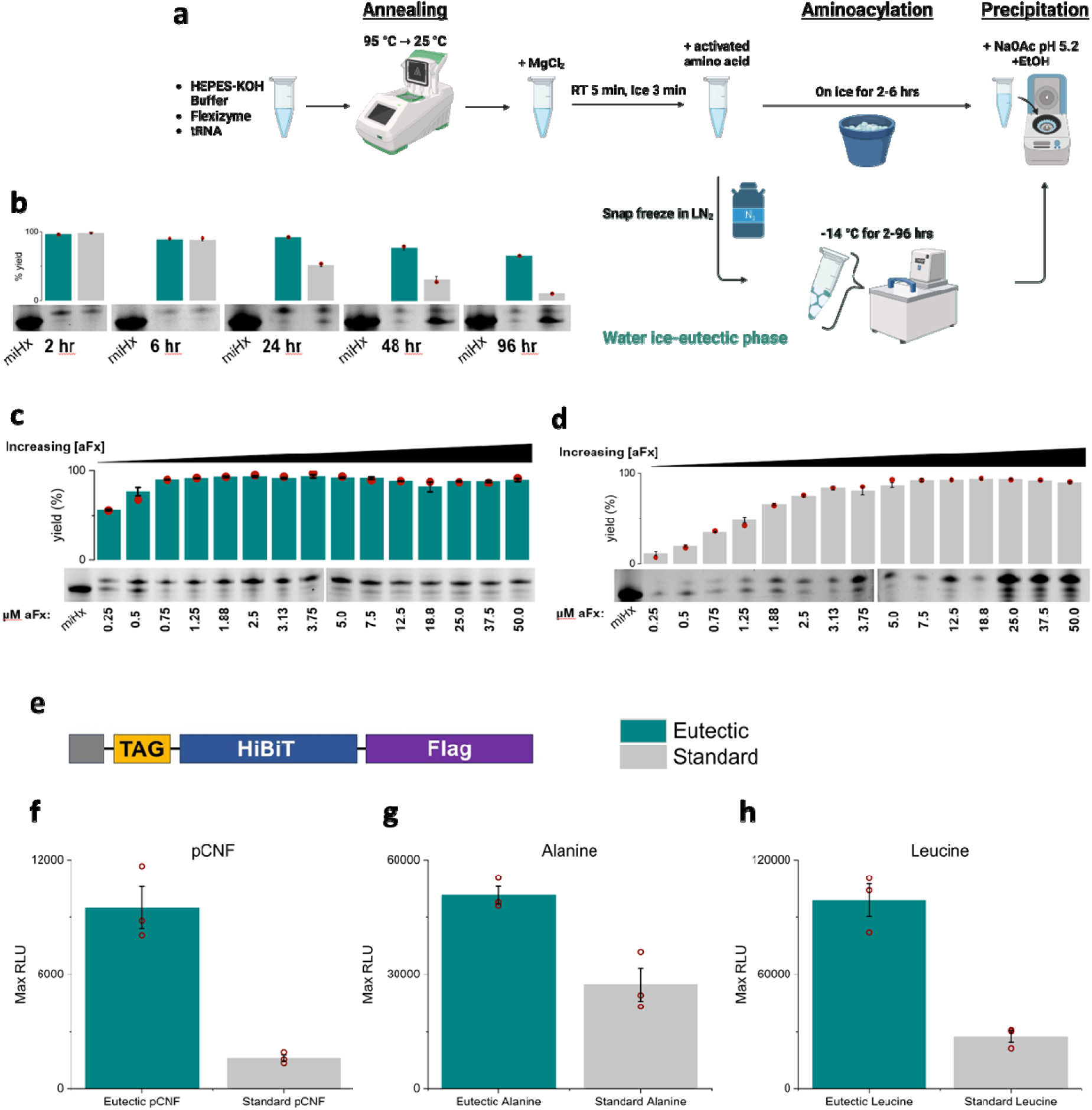
The workflow and advantages of the eutectic flexizyme reactions. **a)** A comparison of the workflow between standard and eutectic flexizyme reactions. Eutectic reactions use the same reagents as standard reactions, but with reduced concentrations of HEPES-KOH buffer, MgCl2, and DMSO. Eutectic flexizyme reactions are snap frozen after adding the activated amino acid, then the reaction is initiated by incubation at -14 °C in an ethylene glycol-water bath. The reaction is quenched with low pH and precipitated with ethanol like a standard flexizyme reaction. **b)** Representative gel image comparing a flexizyme reaction time course (aFx) using eutectic vs standard reaction conditions from 2-96 hours. Dark teal bars are eutectic conditions and gray bars are standard flexizyme conditions. Bars represent the average yield of n=3 replicates, and error bars indicate the standard error of the mean for this data set. Red circles show the yield from each lane of the gel image. Gel analysis in supplementary figures S16-20. **c)** Representative gel image of showing aminoacylation yield with increasing concentrations of aFx in eutectic reaction conditions. Gel analysis in supplementary figures S39-44. **d)** Representative gel image of showing aminoacylation yield with increasing concentrations of aFx in standard reaction conditions. Gel analysis in supplementary figures S45-52. **e)** A representative diagram of the translated HiBiT template with amber codon. **f-g)** tRNA aminoacylated using eutectic vs standard flexizyme reaction conditions used to decode a stop codon suppression in PURExpress, with **(b)** 4-cyano-L-phenylalanine (pCNF), **(c)** L-Alanine, or (**d)** L-Leucine. Error bars show the standard error of the mean for n=3 replicates. Dark teal: eutectic conditions, gray: standard conditions. Red circles represent individual data points.

**Figure 2.**
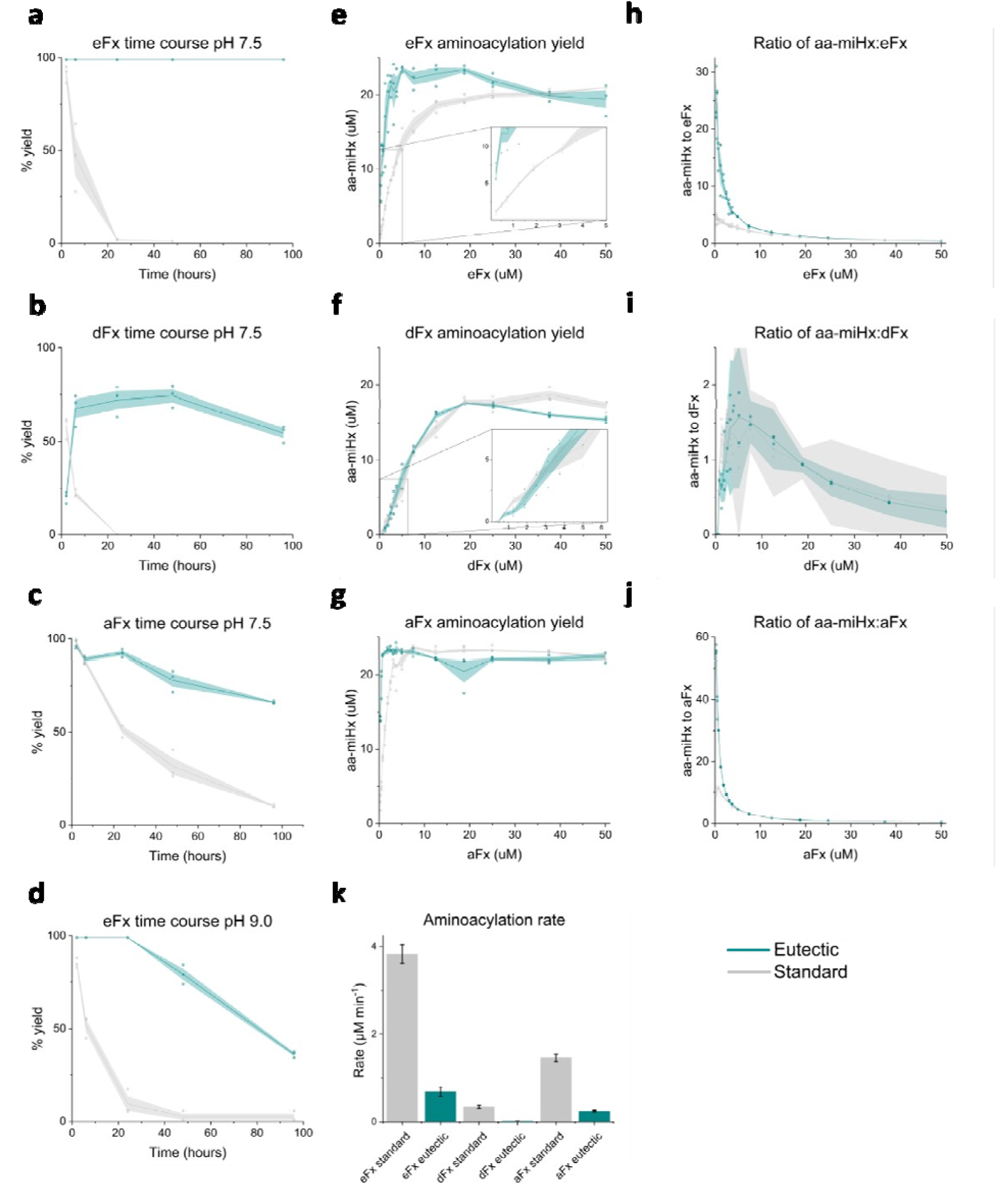
Characterization of flexizyme aminoacylation in a eutectic phase. **a-d)** Time course analysis of aminoacylation catalyzed by standard and eutectic flexizyme reactions. The Y axes on panels are μM aminoacylated miHx (aa-miHx) calculated using the peak intensity ratio between the aminoacyl and total intensity of both gel bands. **a)** Aminoacylation of L-tyrosine-CME using eFx from 2-96 hours at pH 7.5. **b)** Time course of L- alanine-DBE aminoacylation using dFx from 2-96 hours at pH 7.5. **c)** L-leucine-ABT aminoacylation from 2-96 hours at pH 7.5. **d)** L-tyrosine-CME aminoacylation from 2-96 hours at pH 9. Gel analysis in supplementary figures S1-20 **e-g)** Yield of aminoacylation of miHx at varying concentration of flexizyme. miHx concentration is always 25 μM. **e)** L-Tyr-CME aminoacylation using eFx. **f)** L-Ala-DBE aminoacylation using dFx. **g)** L-Leu-DBT aminoacylation using aFx. The inset plots on **e** and **f** enlarge the plot at low concentrations of flexizyme. Gel analysis in supplementary figures S21-52, S77-80 **h-j)** Comparison of the ratio of aminoacyl miHx to flexizyme with varying flexizyme concentrations. Ratio of aa-miHx to flexizyme was calculated by dividing the concentration of aa-miHx by for each concentration of flexizyme **h)** L-Tyr-CME aminoacylation with eFx **i)** L-Ala- DBE aminoacylation with dFx **j)** L-Leu-ABT aminoacylation with aFx. **k)** Comparison of aminoacylation rates of standard and eutectic reaction conditions for eFx (L- Tyr-CME), dFx (L-Ala-DBE), and aFx (L-Leu-ABT). The errors in the rates are the parameter standard errors derived from the fits (supplementary figure S87, curves in red). Gel analysis in supplementary figure S88. For all panels: Dark teal and gray lines represent the average yield from n=3 replicates for eutectic and standard reactions respectively. Circles show individual reaction yields from these replicates. The shaded areas represent the standard error of the mean of each data set.

### Increased translation yields

Magnesium carryover from flexizyme reactions has been shown to inhibit *in vitro* translation and is especially problematic when using multiple flexizyme aminoacyl tRNAs, such as in aaRS-free translation (Chen et al., 2021). We hypothesized that lower magnesium concentrations in eutectic reactions could reduce magnesium carryover into *in vitro* translation. Because eutectic flexizyme reactions use only 20 mM MgCl2, while standard flexizyme reactions use up to 600 mM MgCl_2_ decreased magnesium carryover should result in increased translation yield compared to standard flexizyme conditions. To test this, we aminoacylated 4-cyano-L-phenylalanine (pCNF) onto a stop codon suppressor tRNA, GluE2, using either eutectic or standard reaction conditions (Katoh et al., 2017). We used the HiBiT peptide as a model for this translation assay, linking the production of the peptide to an increase in luminescence (Dixon et al., 2016). We prepared a DNA template with an Amber codon before the HiBiT coding region (**Figure 1g**). We compared translation yields in reactions using aminoacyl tRNA prepared via standard or eutectic flexizyme conditions. The translation yield, measured as luciferase luminescence, in reactions using aminoacyl tRNA from eutectic flexizyme reactions was greater than the tRNA from the standard flexizyme reaction (**Figure 1f-g**).

Those experiments demonstrate the major advantages of the eutectic protocol over the standard flexizyme protocol: the aminoacylation reactions are the most commonly a means to an end, with the *in vitro* translation being the final goal. The use of eutectic conditions, without the need for HPLC purification, allows for significantly increased yield in translation with flexizyme aminoacylated tRNA.

### Flexibility in reaction time

Because previous literature has shown increased RNA stability and decreased rates of hydrolysis at low temperatures (Monnard et al., 2003), (Attwater et al., 2010), the stability of the aminoacyl bond in water ice-eutectic conditions was tested from 2 to 96 hours (**Figure 2a-d**).

We demonstrate that the aminoacylation product in the eutectic phase is remarkably stable compared to standard flexizyme reactions, even at higher pH (**Figure 2b**). Using eFx and L-Tyr-CME, nearly all the microhelix (miHx) was aminoacylated within two hours under eutectic conditions. This yield did not decrease even after 96 hours of incubation (**Figure 2a**). In the same reaction under standard flexizyme conditions, there was little aminoacyl product remaining after 24 hours (**Fig. 2a**). This stability of aminoacylation product highlights the increased flexibility of reaction timing in the eutectic phase.

Most eutectic phase reactions are stable and high yielding between 2 and 96 hours. We analyzed the time course of aminoacylation of the microhelix with Tyr-CME, Alanine-DBE, and Leucine-ABT using their respective flexizymes. In all cases, aminoacylation in the eutectic conditions remained stable far longer than a standard flexizyme reaction (**Figure 2a-d**). Slow hydrolysis means flexizyme reactions that would have different optimal incubation times using a standard reaction can be set up simultaneously using the eutectic conditions, and left to incubate within a wide margin of time before the reactions need to be quenched. Different flexizyme reactions can be initiated and processed together at a flexible end time, without the need to monitor each individual reaction. New flexizyme amino acids usually need to be tested to determine ideal reaction time, but with eutectic reactions this step could be omitted from the workflow. This is especially useful when scaling up the number of flexizyme reactions for radically reprogrammed genetic codes (Passioura et al., 2018) and aaRS-free translation (Chen et al., 2021). Increased product stability at higher pH is vital with difficult to amino acylate monomers like aminobenzoic acid derivatives, which can take 144-192 hr (6-8 days) at pH 9-10.5 to reach substantial yields (Katoh & Suga, 2020).

Surprised by the high yields from reactions in the eutectic phase, we hypothesized increased yields might result from an increase in the aminoacyl miHx:flexizyme ratio, compared to standard reaction conditions. To test this, we titrated the concentration of flexizyme from 0.25 μM to 50 μM, while keeping miHx concentrations constant at 25 μM. Relative to standard reaction conditions, we observed increases in aminoacylation yield, especially at low concentrations of flexizyme. We found a stark increase in the ratio of aminoacyl miHx to flexizyme for the eutectic reactions compared to standard reaction conditions (**Figure 2h-j**). This trend was present for eFx and aFx, but interestingly no differences were observed using dFx.

For eFx and aFx at 0.25 μM, we observed ∼6X increase in aa-miHx:flexizyme ratio over standard reactions. We also varied the concentration of miHx from 0.25-50 μM, while keeping flexizyme constant at 25 μM. In contrast to varying the flexizyme, the ratio between aminoacyl and non-aminoacyl miHx follows a linear relationship: when flexizyme is in excess, miHx approaches maximum acylation yield for an amino acid (Figures S84-86). Observed rates of aminoacyl bond formation for eFx, dFx, and aFx were slower in the eutectic phase than in a standard reaction, yet for both standard and eutectic reactions, all amino acids tested (except eutectic L-Ala-DBE) reached maximal yield in a standard two hour reaction time (**Figure 2k**). These experiments demonstrate that using an eutectic flexizyme protocol the reaction time becomes significantly less critical. It is possible to set up the reactions and focus elsewhere, returning to purify the aminoacylated tRNA at a convenient time.

### Conclusions and outlook

The urgent need to shift from petrochemicals to sustainable bioeconomy results in the increased importance of developing new enzymes, a field that has been relying heavily on expanding the chemical repertoire of peptides with unnatural amino acids (Katoh & Suga, 2022). This means it is likely we will see increased reliance on flexizyme in metabolic engineering and industrial applications. The eutectic protocol presented here simplifies the flexizyme aminoacylation, elevating this from a specialized research tool to a simple, widely usable technology.

The eutectic protocol demonstrated here does not rely on expensive equipment, utilizing water baths and freezers commonly present in any research environment suitable for performing *in vitro* translation. We demonstrated generally higher tRNA aminoacylation yields than using the traditional protocol. Lower magnesium carryover enables using those tRNA directly in translation, without the need to HPLC purify the fragile, charged tRNA. The extended, and very flexible, reaction times allow for easy multiplexing, using substrates that have different optimal reaction times in the standard protocol. It also allows for fitting flexizyme aminoacylation into many different workflows, without the need to plan around the sensitive timing of the flexizyme protocols. Taken together, this lowers the barrier of entry for using multiple unnatural amino acids, especially those with noncanonical peptide backbones.

## Materials and methods

### Synthesis of activated amino acids

L-phenylalanine-CME and 4-cyano-L-phenylalanine-CME were synthesized as reported previously (Robertson et al., 1991), (Murakami et al., 2006). L-alanine-DBE was synthesized as reported previously (Murakami et al., 2006). For cyanomethyl esters (CME), 0.83 mmol *N*- butoxycarbonyl (*N*-boc)-L-tyrosine-OH or *N-*boc-4-cyano-L-phenylalanine-OH was dissolved in 1 mL of dimethyl formamide (DMF), 0.3 mL of chloroacetonitrile, and 0.26 mL of diisopropylethylamine (DIPEA) and stirred for 16 hours at room temperature. The reaction was diluted with ethyl acetate (EtOAc) and washed twice with 1 M HCl, twice with saturated NaHCO_3_, and five times with brine. The organic layer was dried over MgSO_4_ and concentrated by rotary evaporation.

For dinitrobenzyl esters (DBE), 0.75 mmol of *N*-boc-L-alanine-OH was dissolved in 1 mL of DMF and 0.6 mL of triethylamine (TEA) was added to initiate the reaction. The mixture was stirred at room temperature for 16 hours. The reaction was diluted with EtOAc and washed twice with 1 M HCl, twice with saturated NaHCO_3_, and five times with brine. The organic layer was dried over MgSO_4_ and concentrated by rotary evaporation.

L-Leu-ABT was synthesized as previously reported (Niwa et al., 2009).

To deprotect, the compounds were mixed with 2 mL of 4 M HCl in dioxane and stirred for 30 minutes. The solution was concentrated by rotary evaporation and the compound precipitated and washed with cold diethyl ether three times. Residual solvent was removed under vacuum overnight. Compounds were verified using ^1^H nuclear magnetic resonance (NMR) dissolved in DMSO-d6 (Figures S74-76).

### Preparation of flexizymes

DNA templates for eFx, dFx, and microhelix (miHx) were prepared via overlap extension and amplification of oligonucleotides purchased from Integrated DNA Technologies (IDT) using polymerase chain reaction (PCR) as reported previously (Goto et al., 2011). These templates were purified using a phenol chloroform extraction and ethanol precipitation before being resuspended in water and run on a 3% agarose gel to confirm the correct PCR product length. Flexizymes were transcribed using a T7 transcription reaction as previously described (Deich et al., 2023). Briefly, the reaction consisted of 1X template, 1X Homemade NEB Buffer, 8 mM GTP, 4 mM A/C/UTP, 0.005X phosphatase 25 ng/μL, 1 μM T7 RNAP, RNAse inhibitor 0.4 U/μL. The transcripts were digested with Turbo DNAse (New England Biolabs) at 37 °C for 20 minutes before being gel purified from a denaturing 10% polyacrylamide gel. The purified flexizyme was desalted using a Monarch RNA purification kit using 2X volume ethanol (New England Biolabs), concentration determined using a Nanodrop ND-1000, diluted to 250 μM, and stored in aliquots at -80 °C. aFx RNA was ordered from IDT and gel purified after receiving.

### Standard flexizyme conditions

Reaction conditions: 50 mM HEPES-KOH pH 7.5, 25 μM miHx or tRNA, 25 μM flexizyme (unless otherwise specified), 600 mM MgCl_2_, 5 mM activated amino acid, 20% DMSO. 1 μL of 500 mM HEPES-KOH pH 7.5, 1 μL of 250 μM miHx, 1 μL of 250 μM eFx, and 3 μL of water were mixed and annealed at 95 °C for 2 minutes and then slowly cooled to 25 °C over 7 minutes. Next, 2 μL of 3 M MgCl_2_ was added and the mixture was incubated on the benchtop for 5 minutes and then on ice for 3 minutes. To initiate the reaction, 2 μL of 25 mM activated amino acid in 100% DMSO was added and mixed thoroughly. The reaction was incubated on ice for two hours (unless otherwise specified). To quench, the reaction mixture was diluted to 80 mM NaOAc pH 5.2 in 70% ethanol then centrifuged at 15,000 x g for 15 minutes at room temperature as reported previously (Goto et al., 2011). The supernatant was removed.

### Water ice-eutectic flexizyme reactions

Reaction conditions: 5 mM HEPES-KOH pH 7.5, 25 μM miHx or tRNA, and 25 μM flexizyme (unless otherwise specified). This mixture was annealed at 95 °C for 2 minutes and then slowly cooled to 25 °C over 7 minutes. Next, 2 μL of 100 mM MgCl_2_ was added (final concentration of 20 mM MgCl_2_), incubated on the benchtop for 5 minutes and then cooled on ice for 3 minutes. Then, 2 μL of activated amino acid in 12.5% DMSO was added and mixed well (final concentration 5 mM and 2.5% DMSO). The reaction was snap frozen in liquid nitrogen and incubated in an ethylene glycol water bath at -14 °C for 18 hours (unless otherwise specified).

To quench, the reaction mixture was diluted to 80 mM NaOAc pH 5.2, 70% ethanol and centrifuged at 15,000 x g at room temperature for 15 minutes as reported previously (Goto et al., 2011). The supernatant was removed.

### Acid PAGE gel shift assay

The miHx-flexizyme pellet was resuspended in 4 μL of 10 mM NaOAc pH 5.2 and 12 μL of acid PAGE loading buffer. The reactions were run at 125 V for 2.5 hours on a denaturing (6 M urea) 20% (w/v) polyacrylamide gel in 50 mM NaOAc pH 5.2 running buffer (using the Biorad Mini-PROTEAN tetra gel, 1 mm gel thickness). The gels were stained with Sybr Gold (Invitrogen) and visualized using an Omega Lum gel imager (Aplegen). Microhelix bands were quantified using ImageJ (Schneider et al., 2012).

### *In vitro* translation reactions

With *in vitro* translation reactions, the tRNA-flexizyme pellet was washed using 100 mM NaOAc in 70% ethanol, vortexing to break up the pellet and centrifuged at 15,000 x g for 5 minutes.

This step was repeated. The pellet was washed once more with just 70% ethanol without vortexing and centrifuged at 15,000 x g for 3 minutes. The tube was dried on the benchtop covered with a kimwipe for 5 minutes. The pellet was suspended in 1.5 μL of 1 mM NaOAc pH 5.2 and added immediately to the *in vitro* translation reaction.

### HiBiT Luminescence Assay

Translation yield from the PURExpress reactions was quantified using Nano-Glo HiBiT Extracellular Detection System (Promega). To make 1X NanoGlo solution, 100 μL of water, 1 μL of LgBiT, and 2 μL of HiBiT extracellular substrate were mixed. A part of the PURExpress reaction was diluted 1:1000 with water before being mixed 1:1 with the 1X NanoGlo solution for a total volume of 20 μL solution. This was immediately put into a plate reader (Molecular Devices) set at 37 °C and luminescence was quantified over time. An average of the maximum five values was recorded as the maximum luminescence value obtained.

## Supplementary information

**Figure S1.**
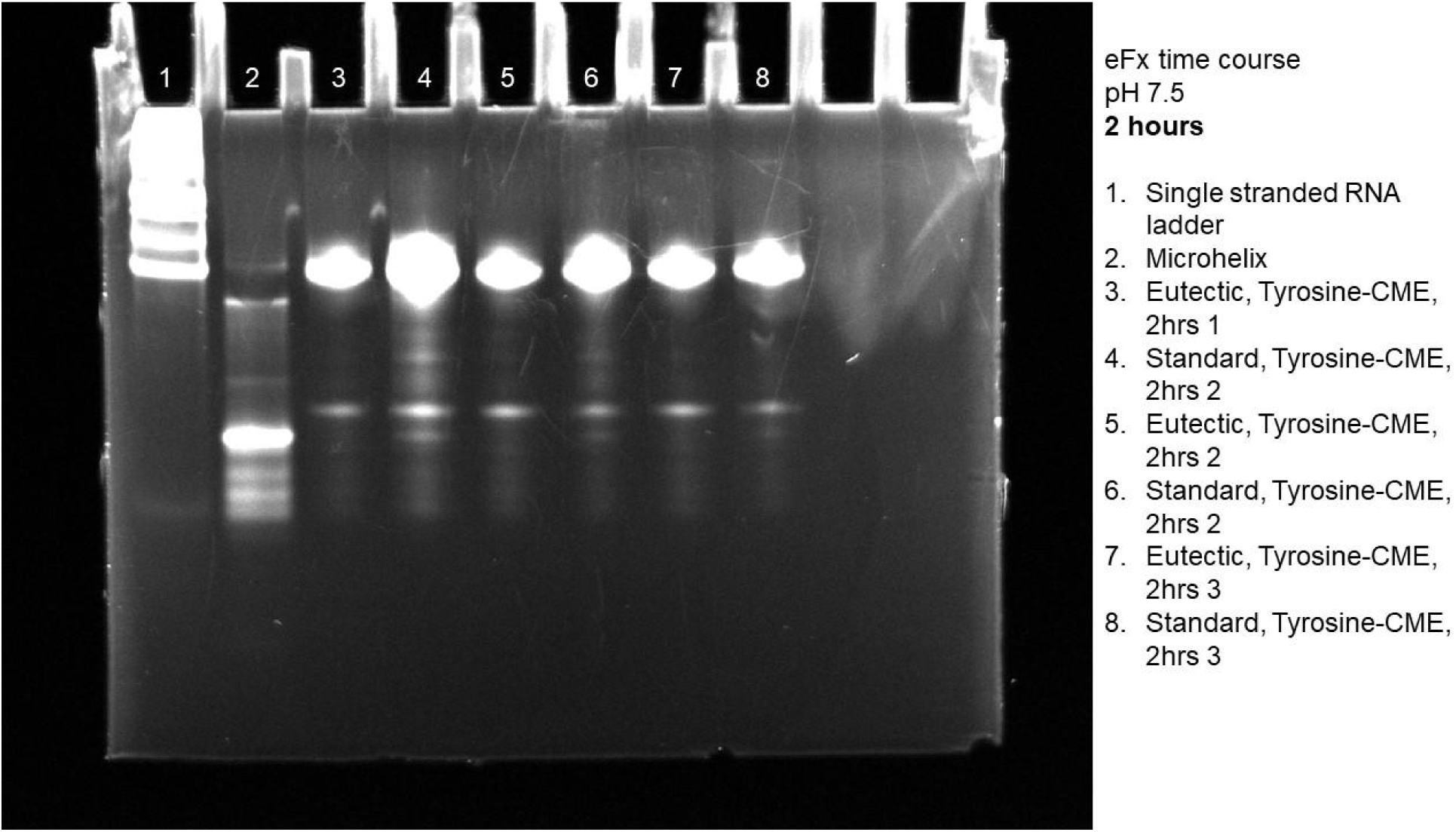
Gel of eutectic and standard reactions for eFx at pH 7.5 using L-Tyr-CME at 2 hours.

**Figure S2.**
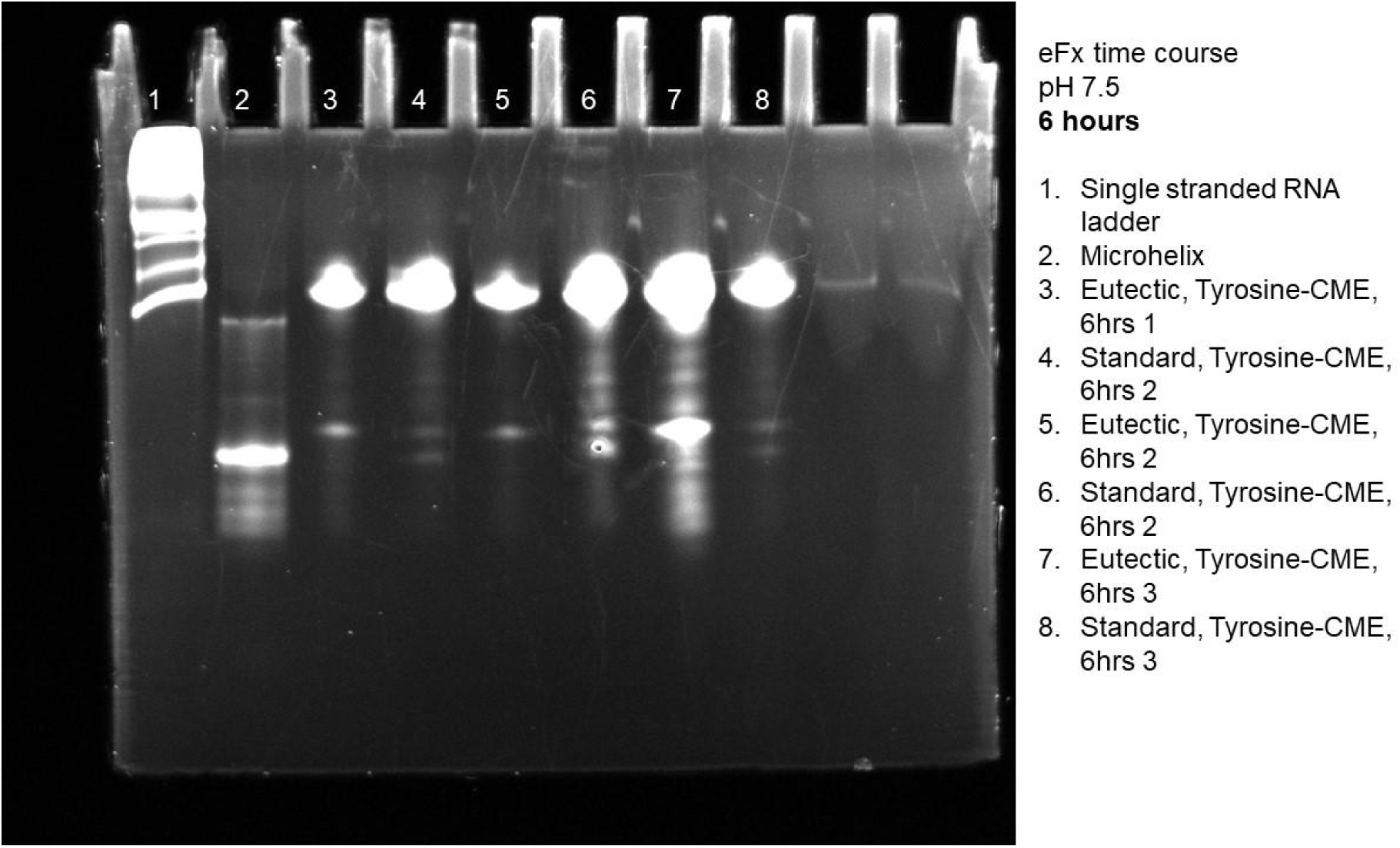
Gel of eutectic and standard reactions for eFx at pH 7.5 using L-Tyr-CME at 6 hours.

**Figure S3.**
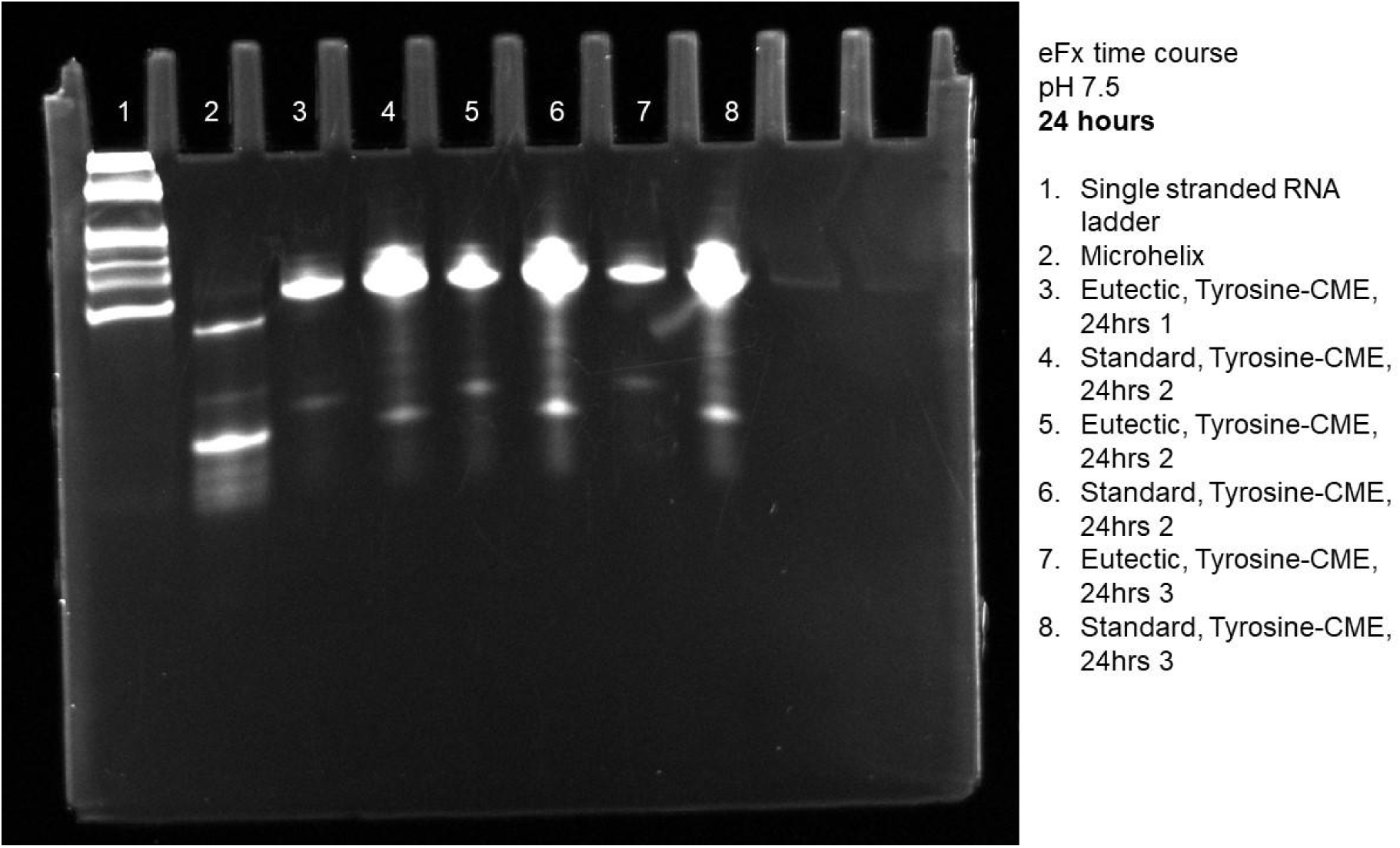
Gel of eutectic and standard reactions for eFx at pH 7.5 using L-Tyr-CME at 24 hours.

**Figure S4.**
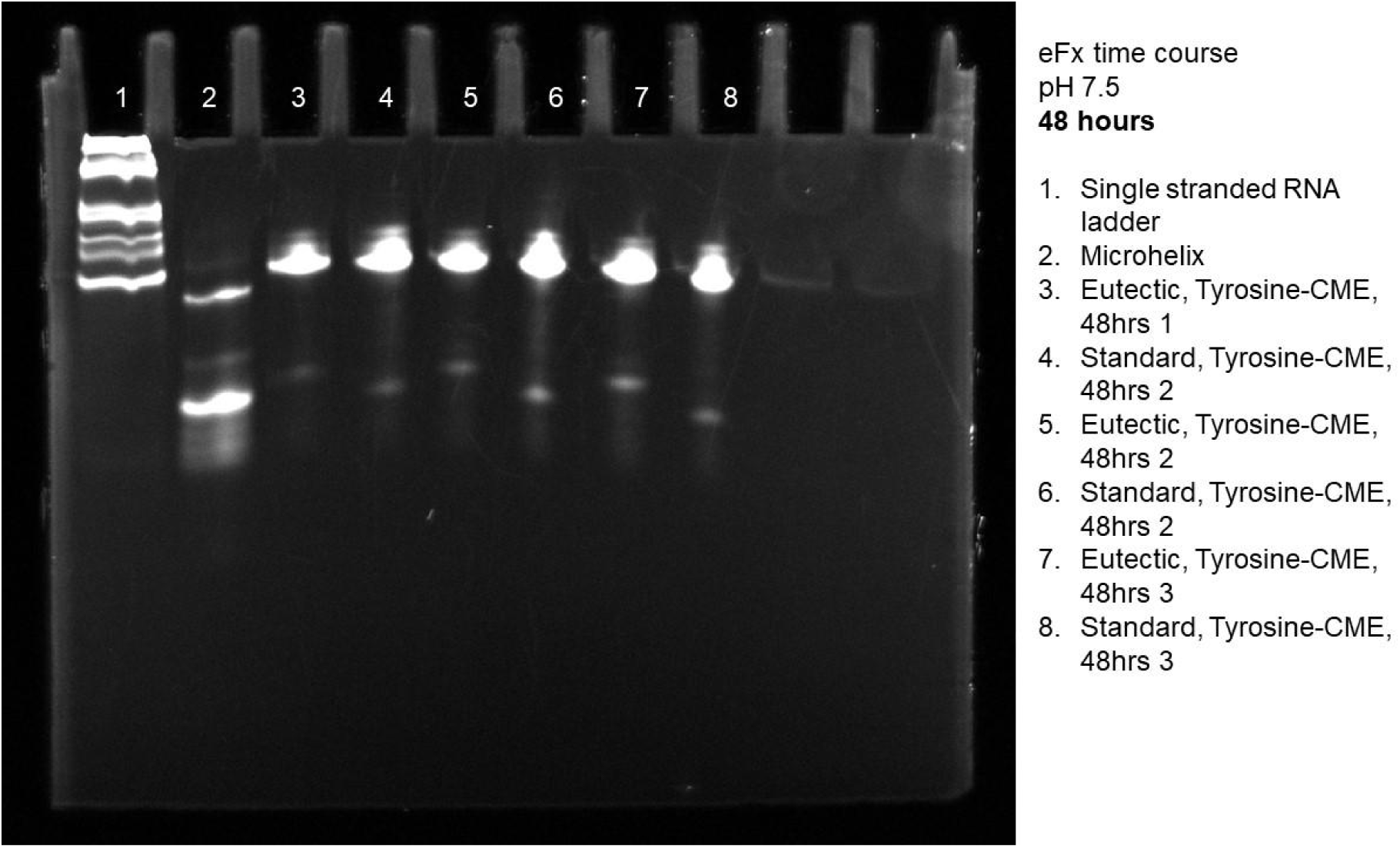
Gel of eutectic and standard reactions for eFx at pH 7.5 using L-Tyr-CME at 48 hours.

**Figure S5.**
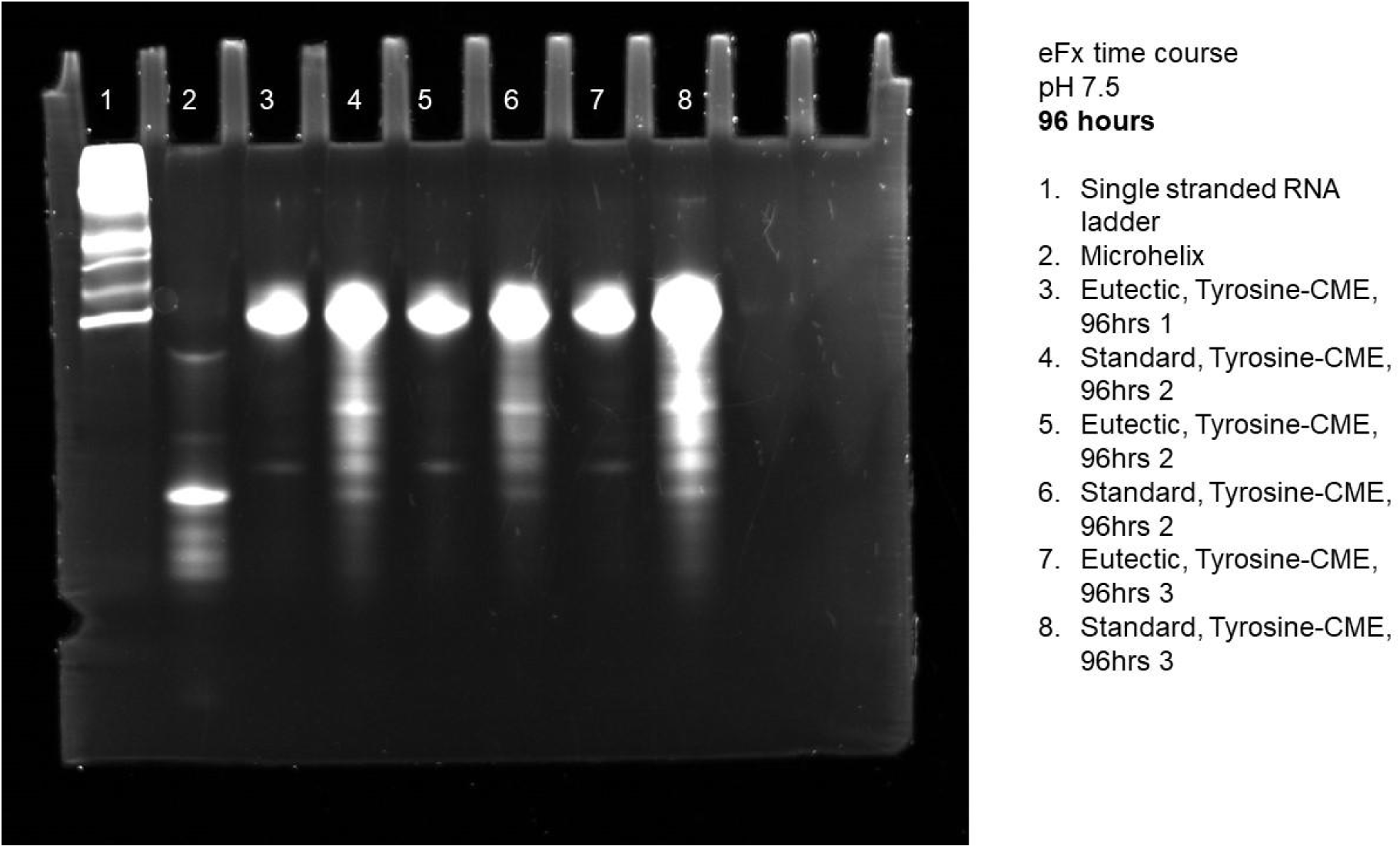
Gel of eutectic and standard reactions for eFx at pH 7.5 using L-Tyr-CME at 96 hours.

**Figure S6.**
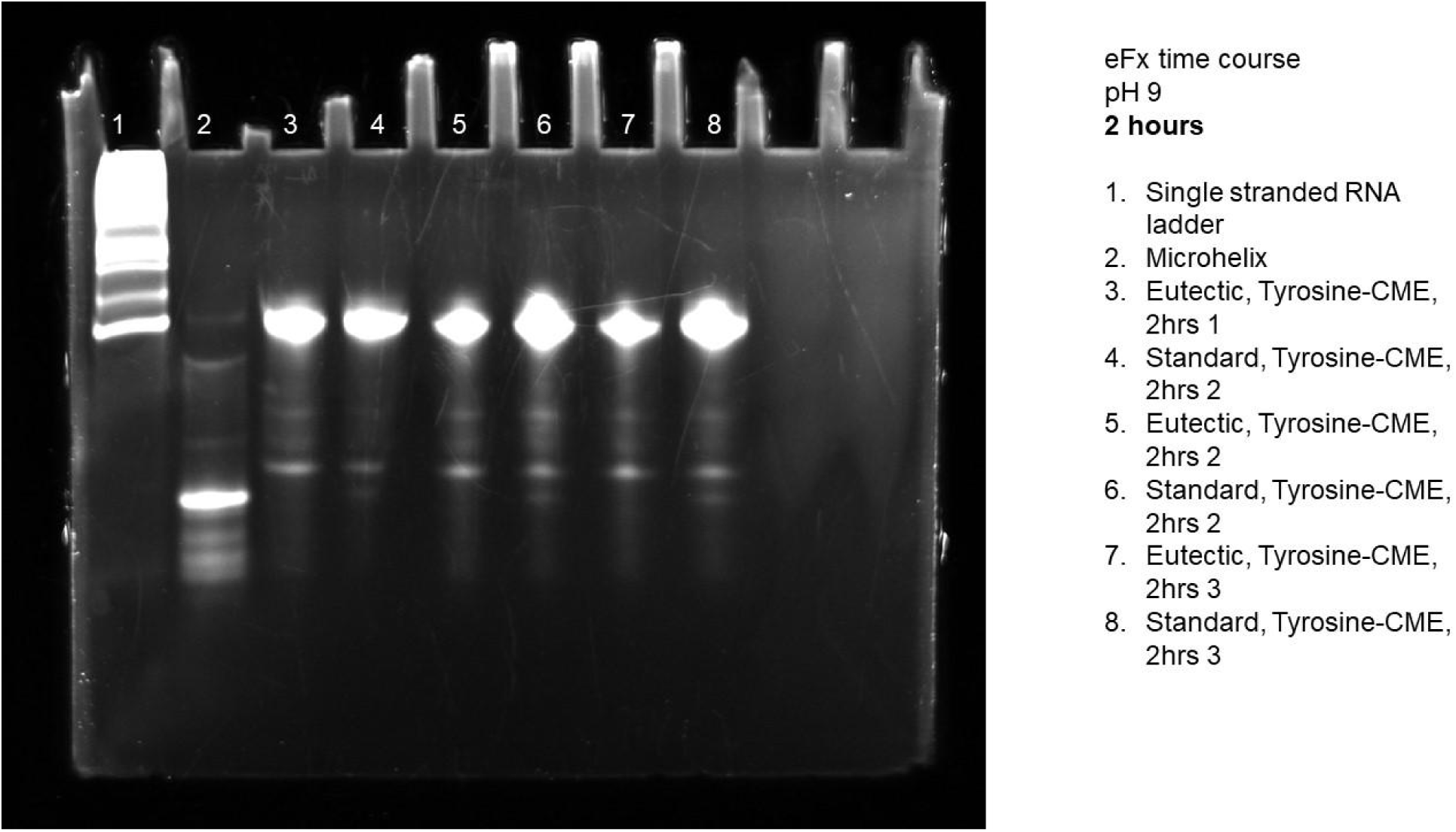
Gel of eutectic and standard reactions for eFx at pH 9 using L-Tyr-CME at 2 hours.

**Figure S7.**
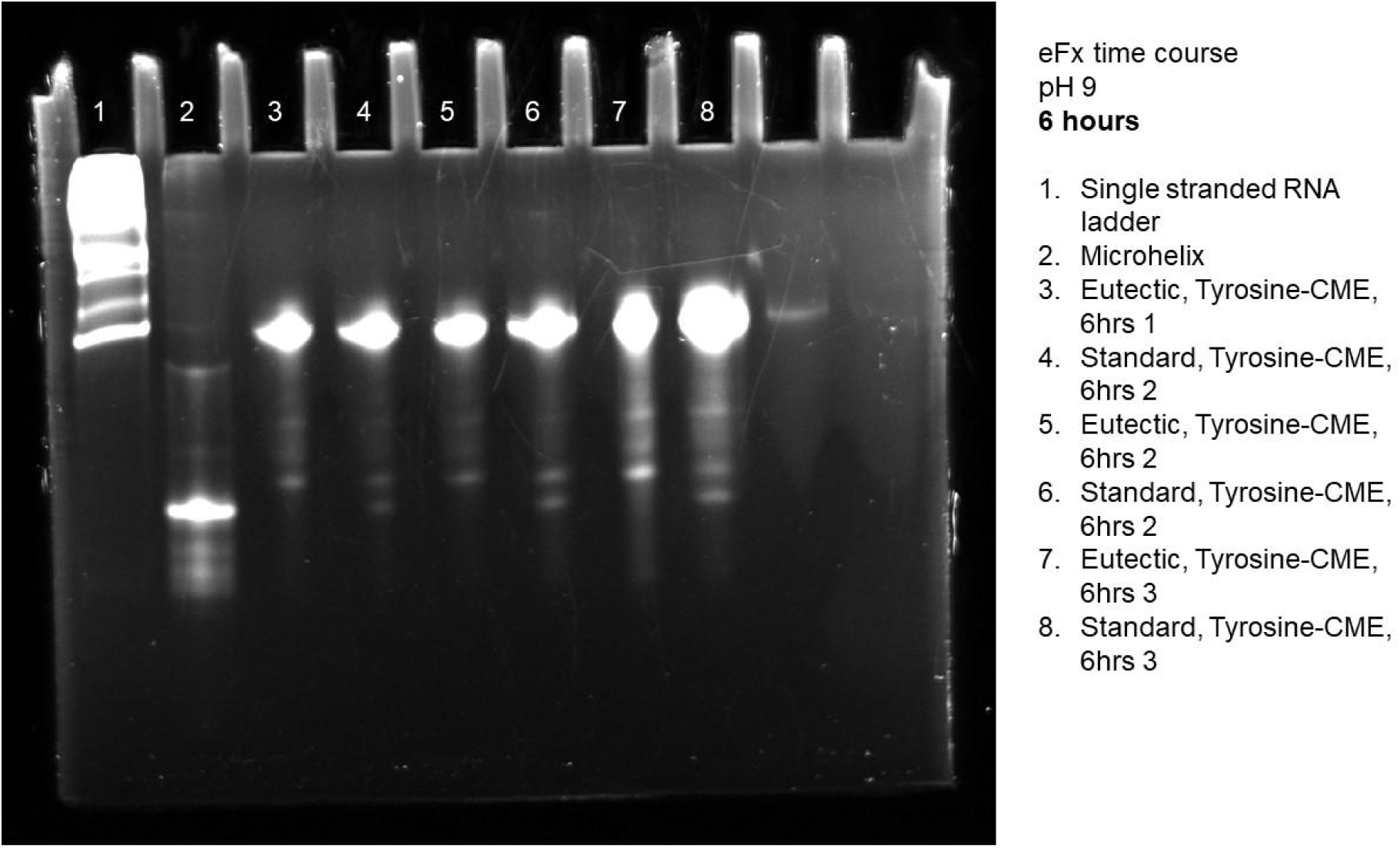
Gel of eutectic and standard reactions for eFx at pH 9 using L-Tyr-CME at 6 hours.

**Figure S8.**
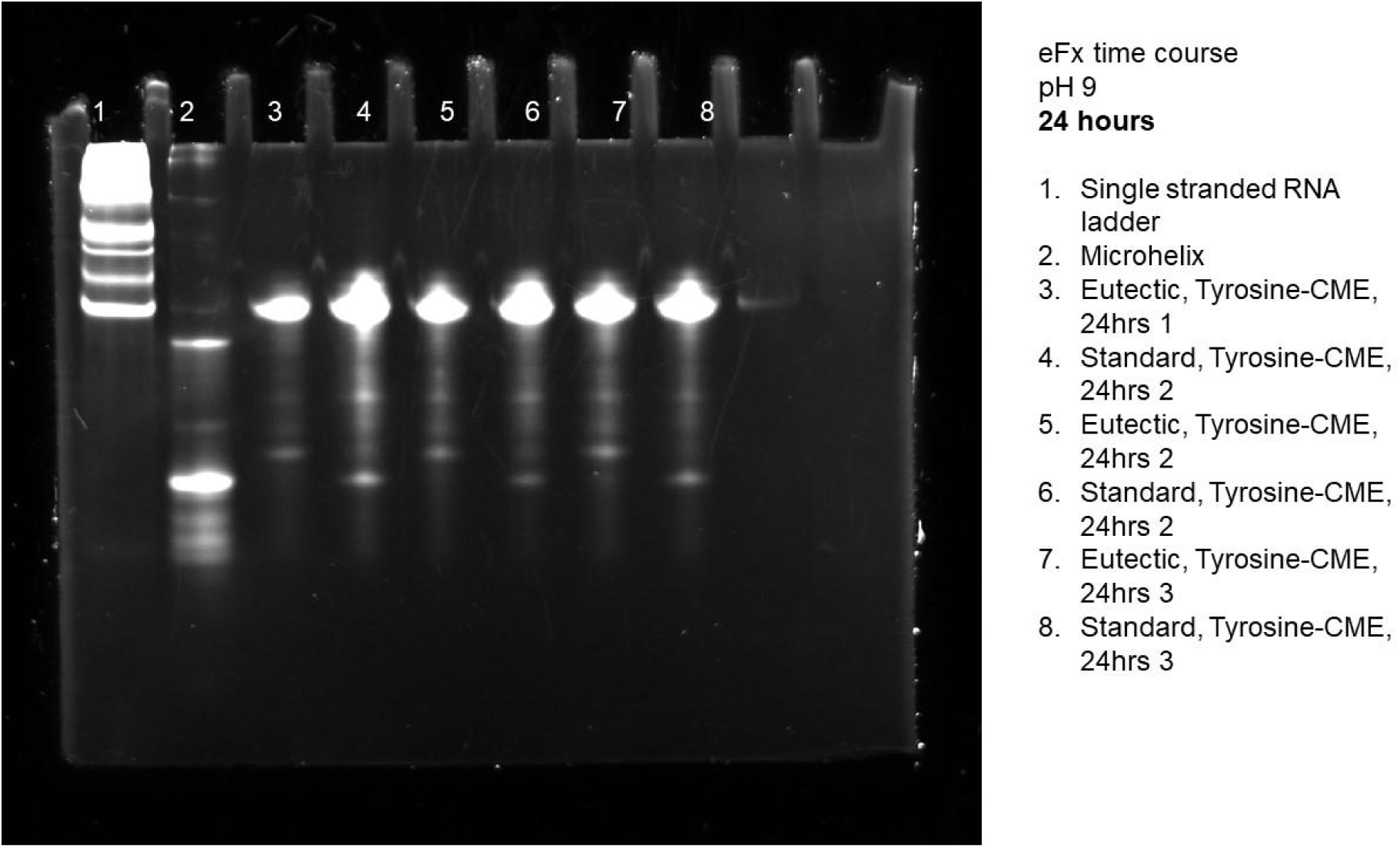
Gel of eutectic and standard reactions for eFx at pH 9 using L-Tyr-CME at 24 hours.

**Figure S9.**
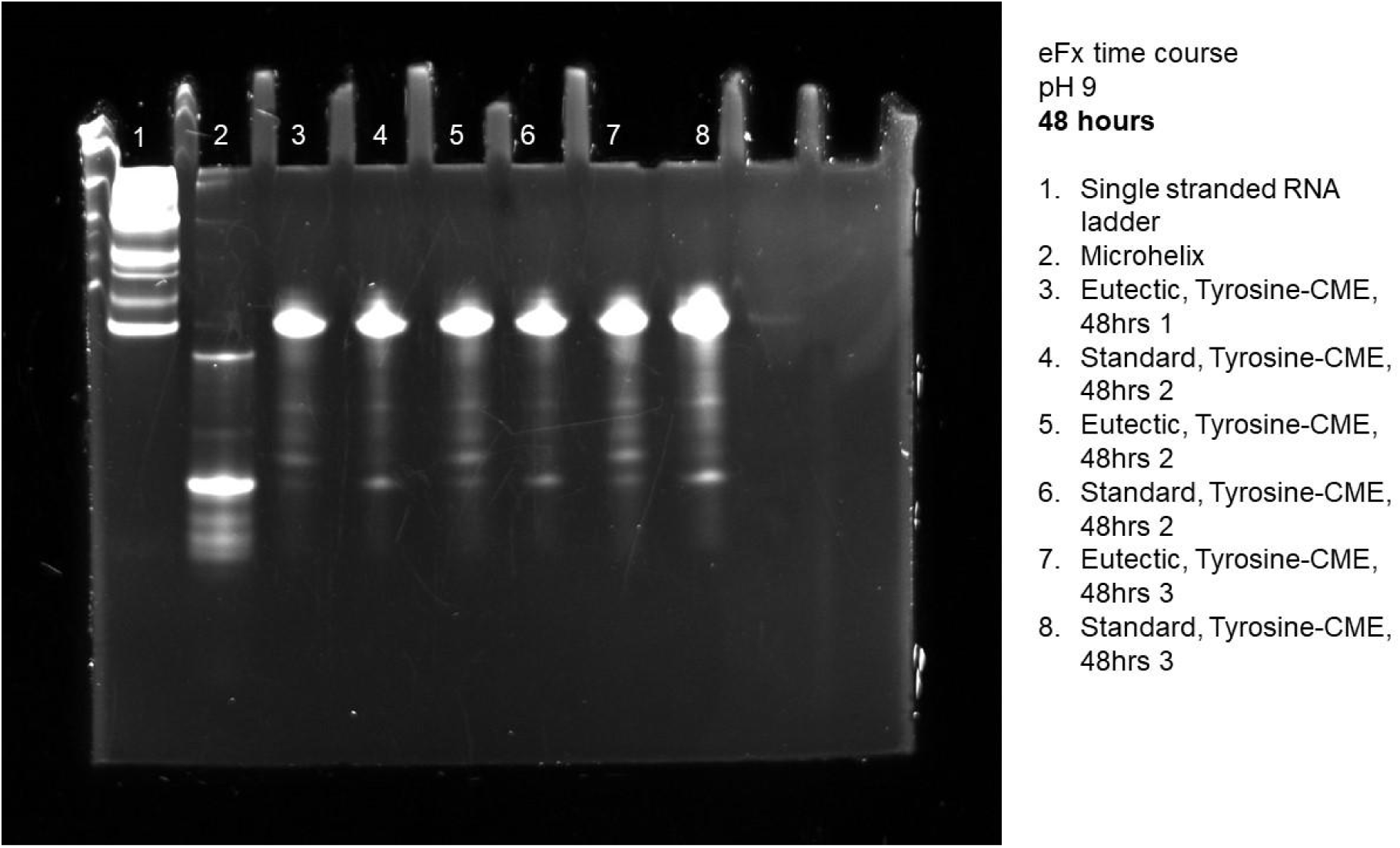
Gel of eutectic and standard reactions for eFx at pH 9 using L-Tyr-CME at 48 hours.

**Figure S10.**
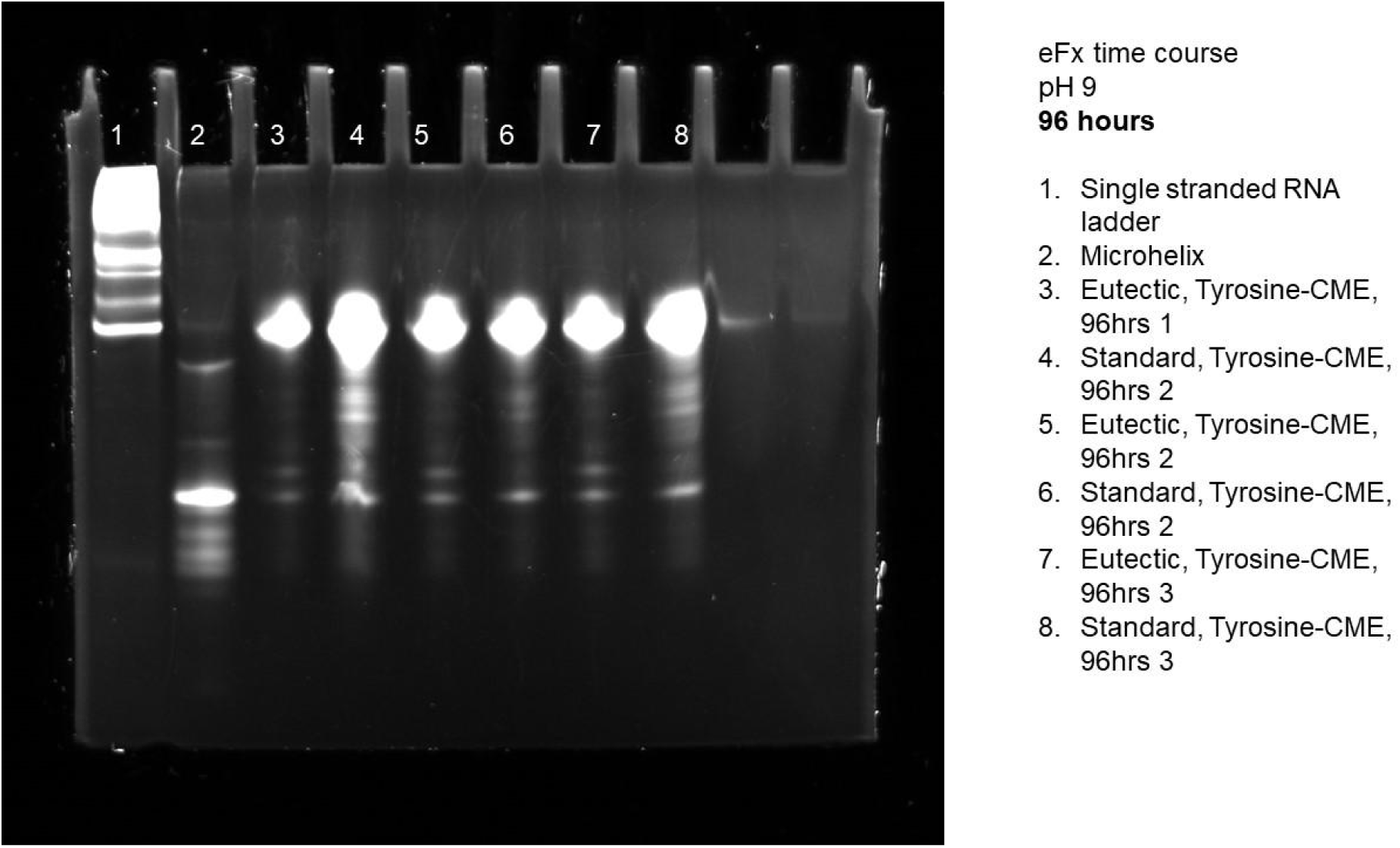
Gel of eutectic and standard reactions for eFx at pH 9 using L-Tyr-CME at 96 hours.

**Figure S11.**
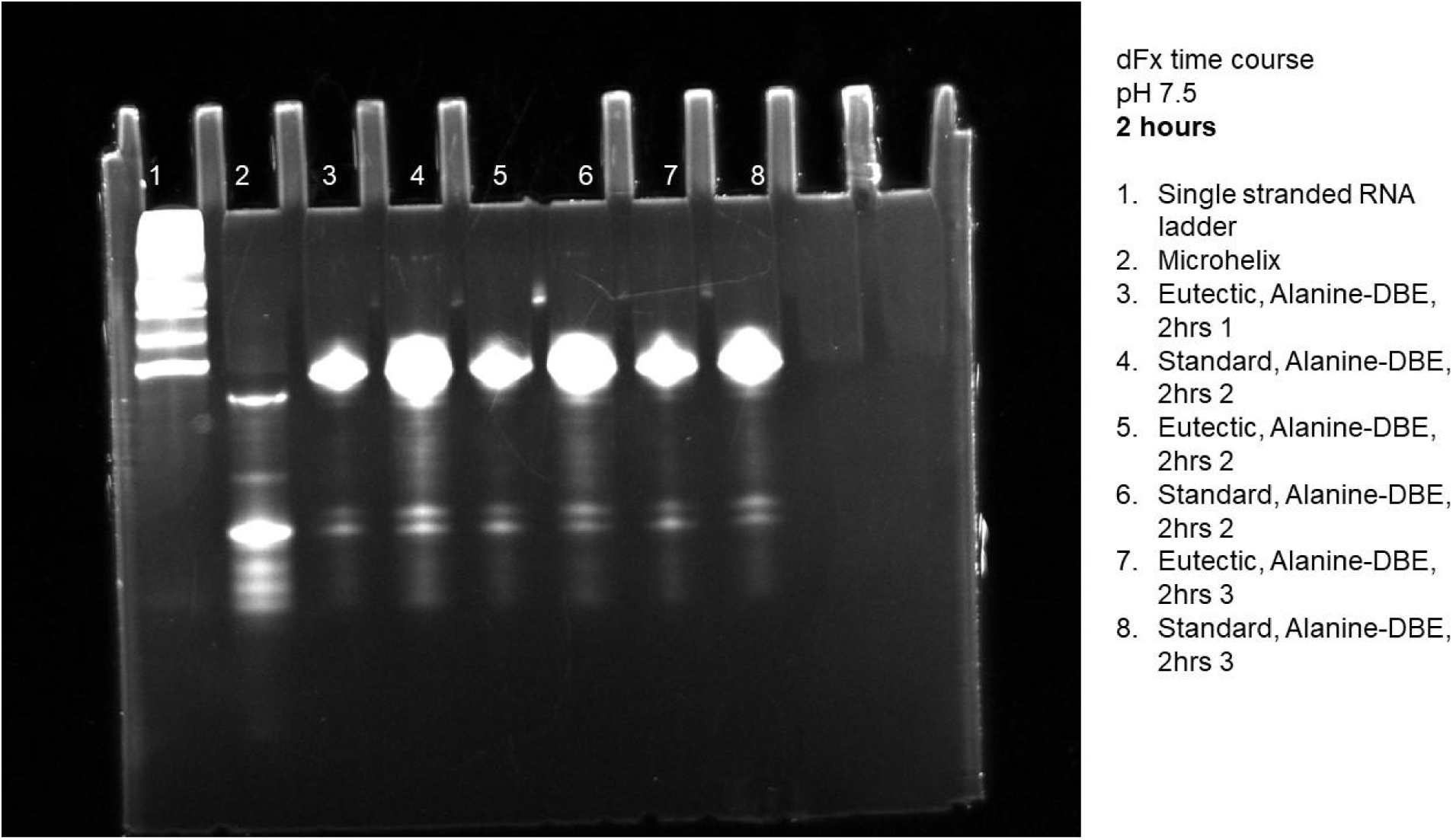
Gel of eutectic and standard reactions for dFx at pH 7.5 using L-Ala-DBE at 2 hours.

**Figure S12.**
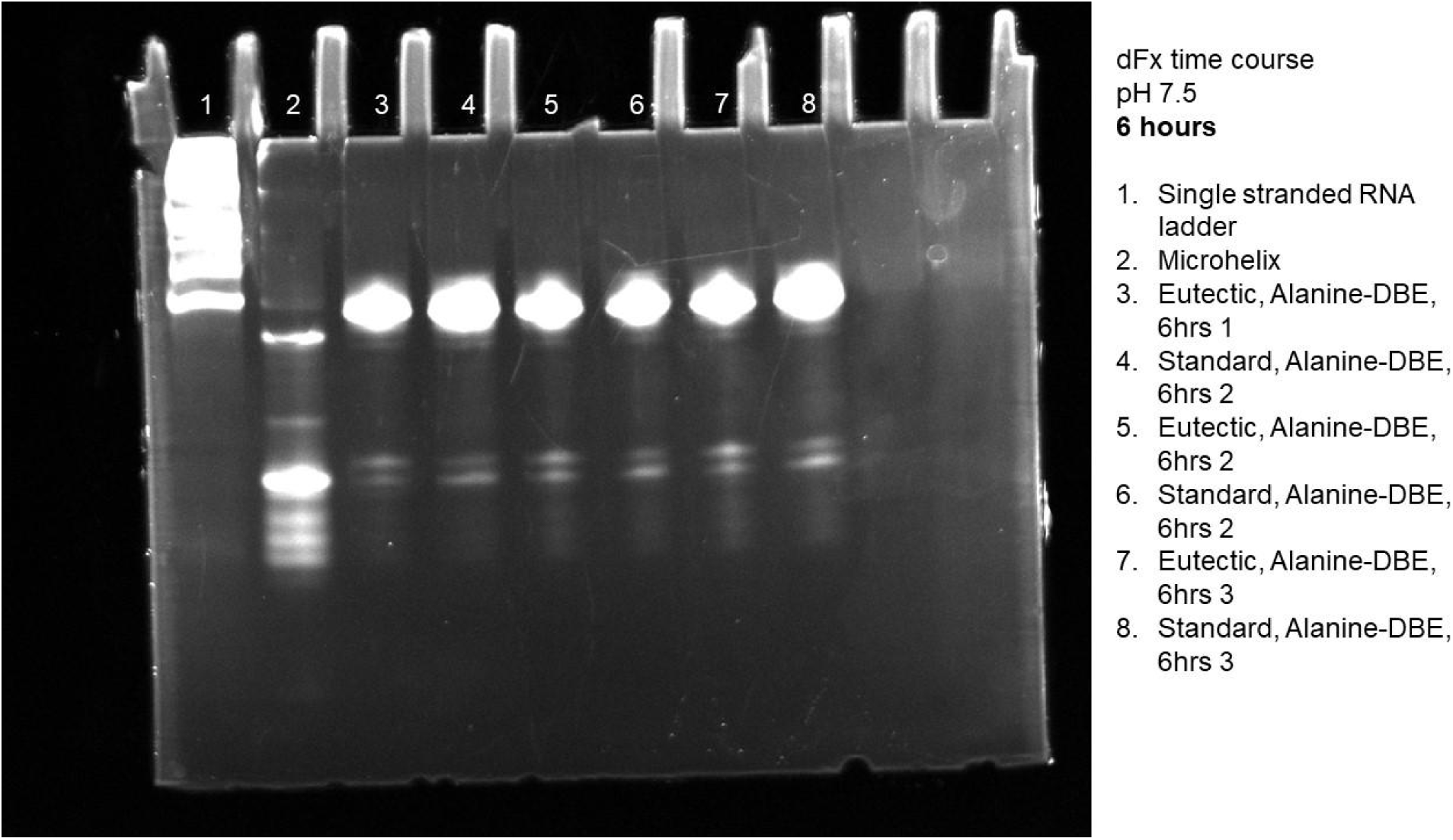
Gel of eutectic and standard reactions for dFx at pH 7.5 using L-Ala-DBE at 6 hours.

**Figure S13.**
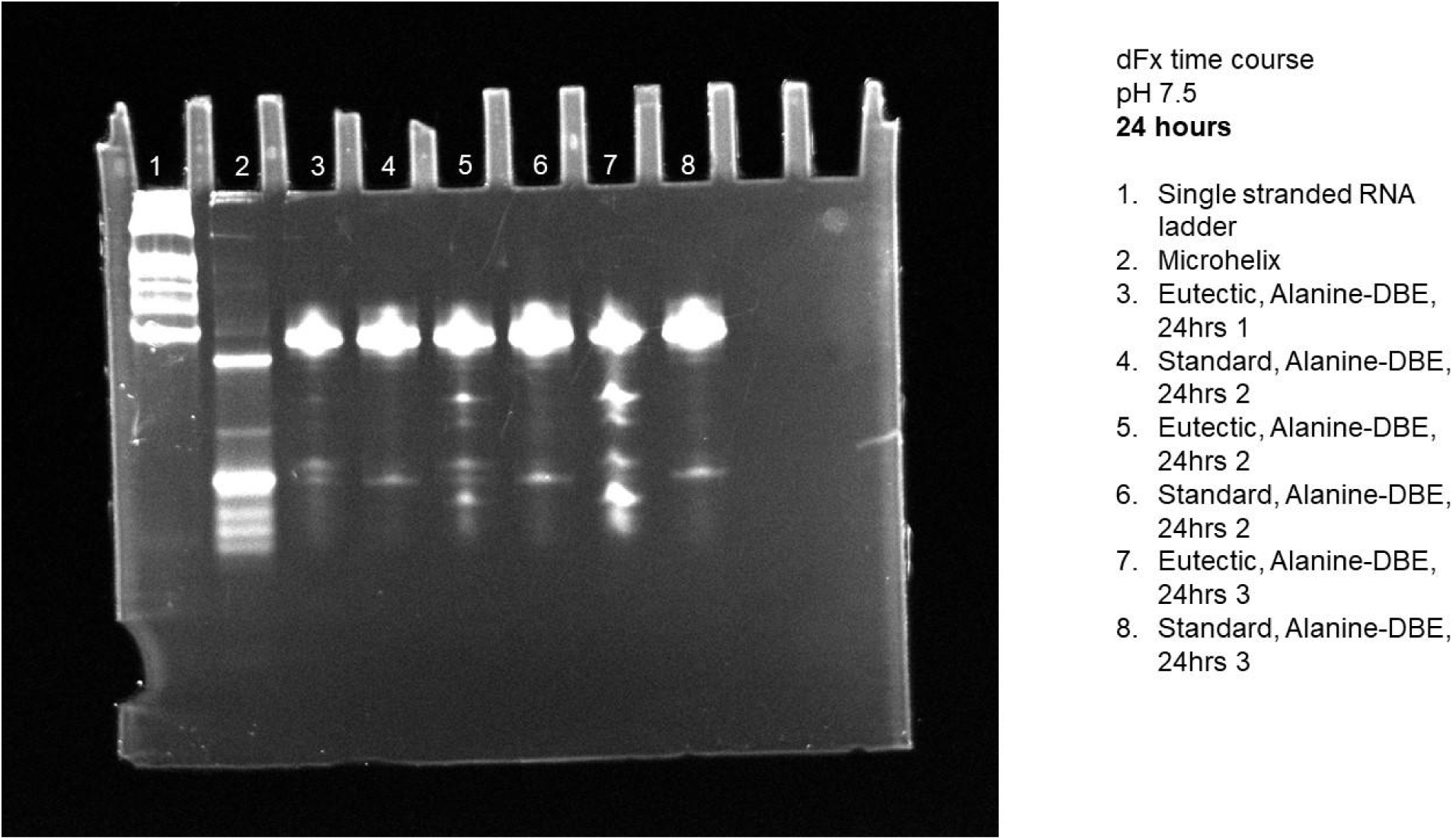
Gel of eutectic and standard reactions for dFx at pH 7.5 using L-Ala-DBE at 24 hours.

**Figure S14.**
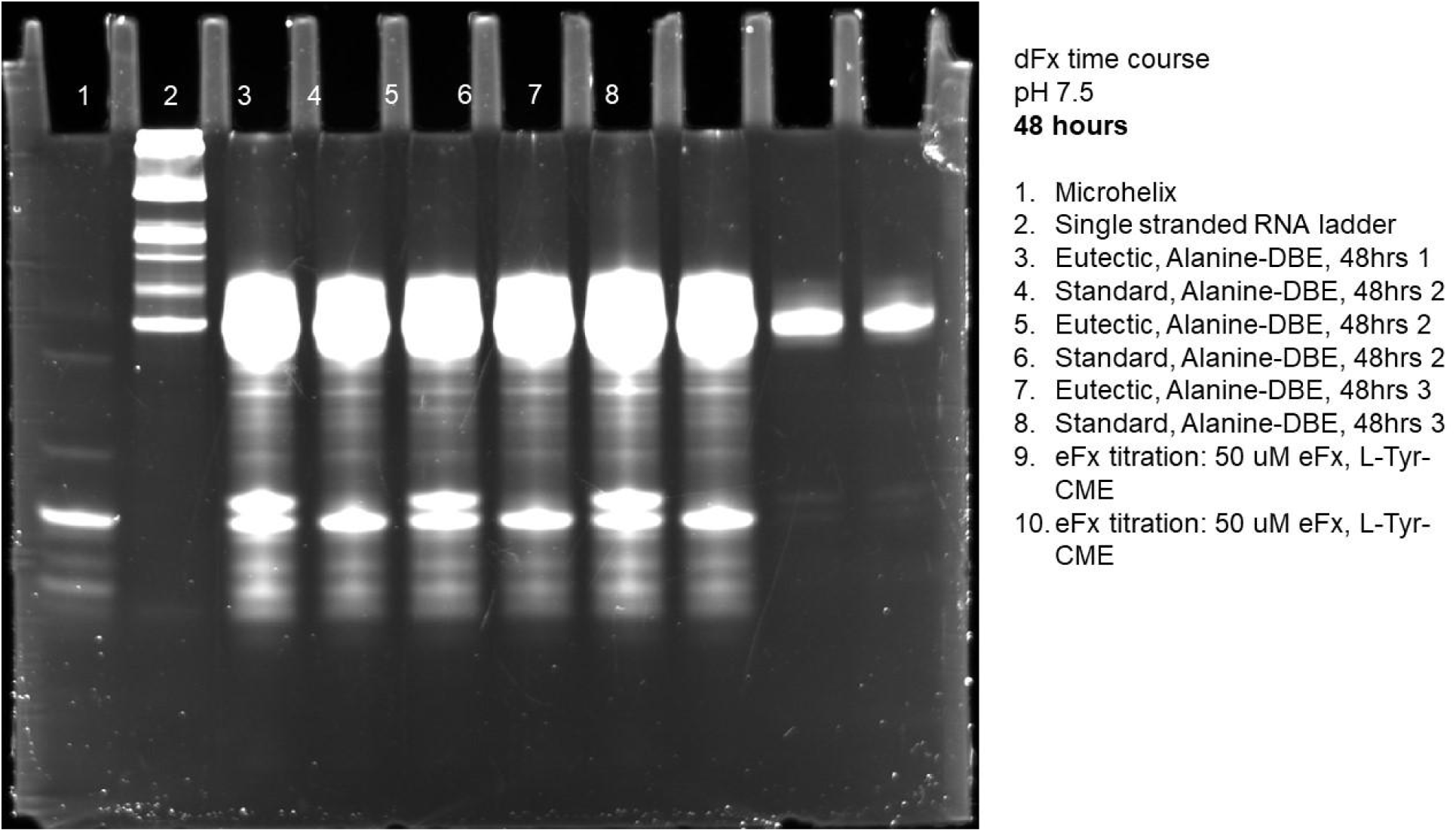
Gel of eutectic and standard reactions for dFx at pH 7.5 using L-Ala-DBE at 48 hours.

**Figure S15.**
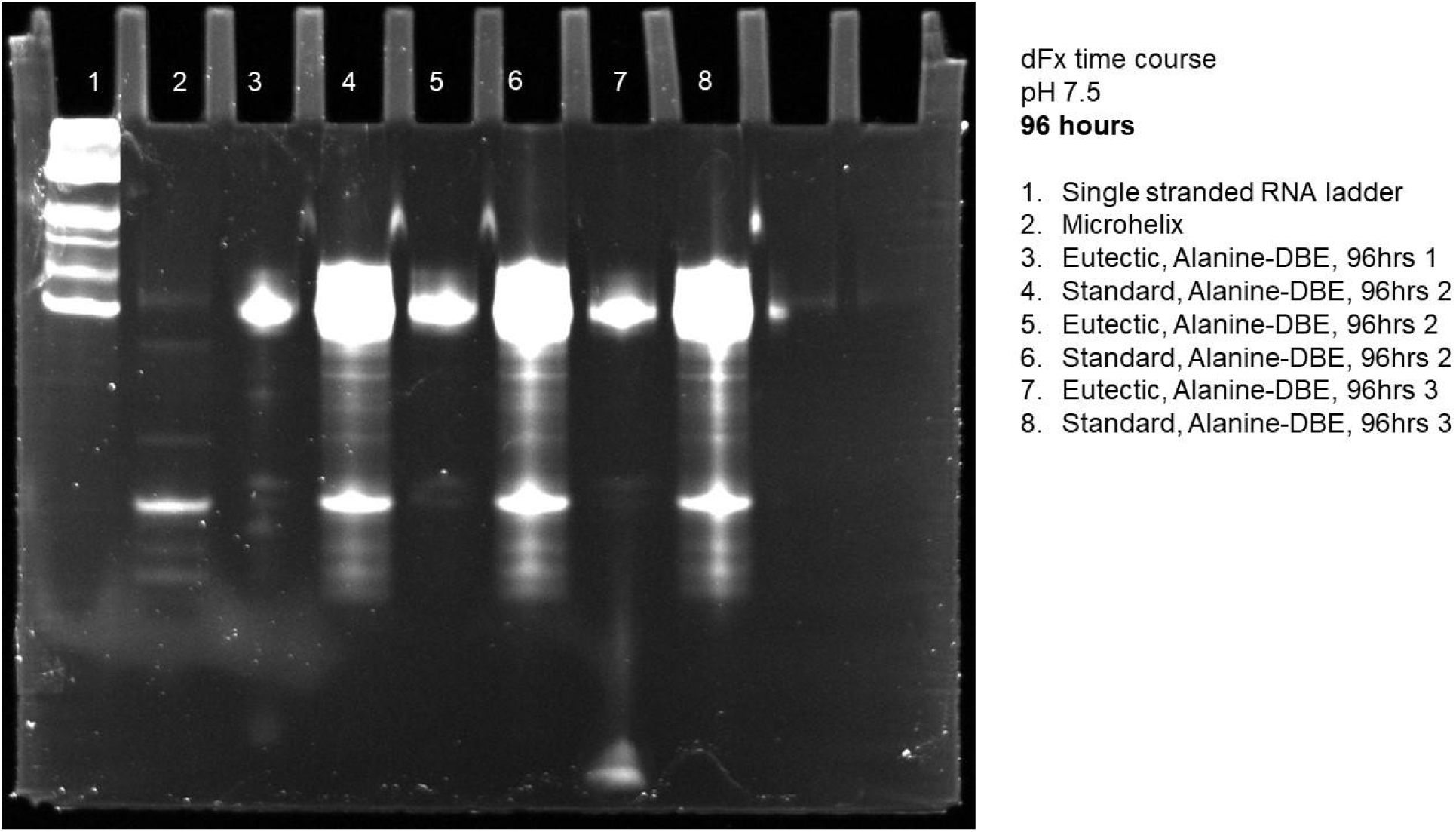
Gel of eutectic and standard reactions for dFx at pH 7.5 using L-Ala-DBE at 96 hours.

**Figure S16.**
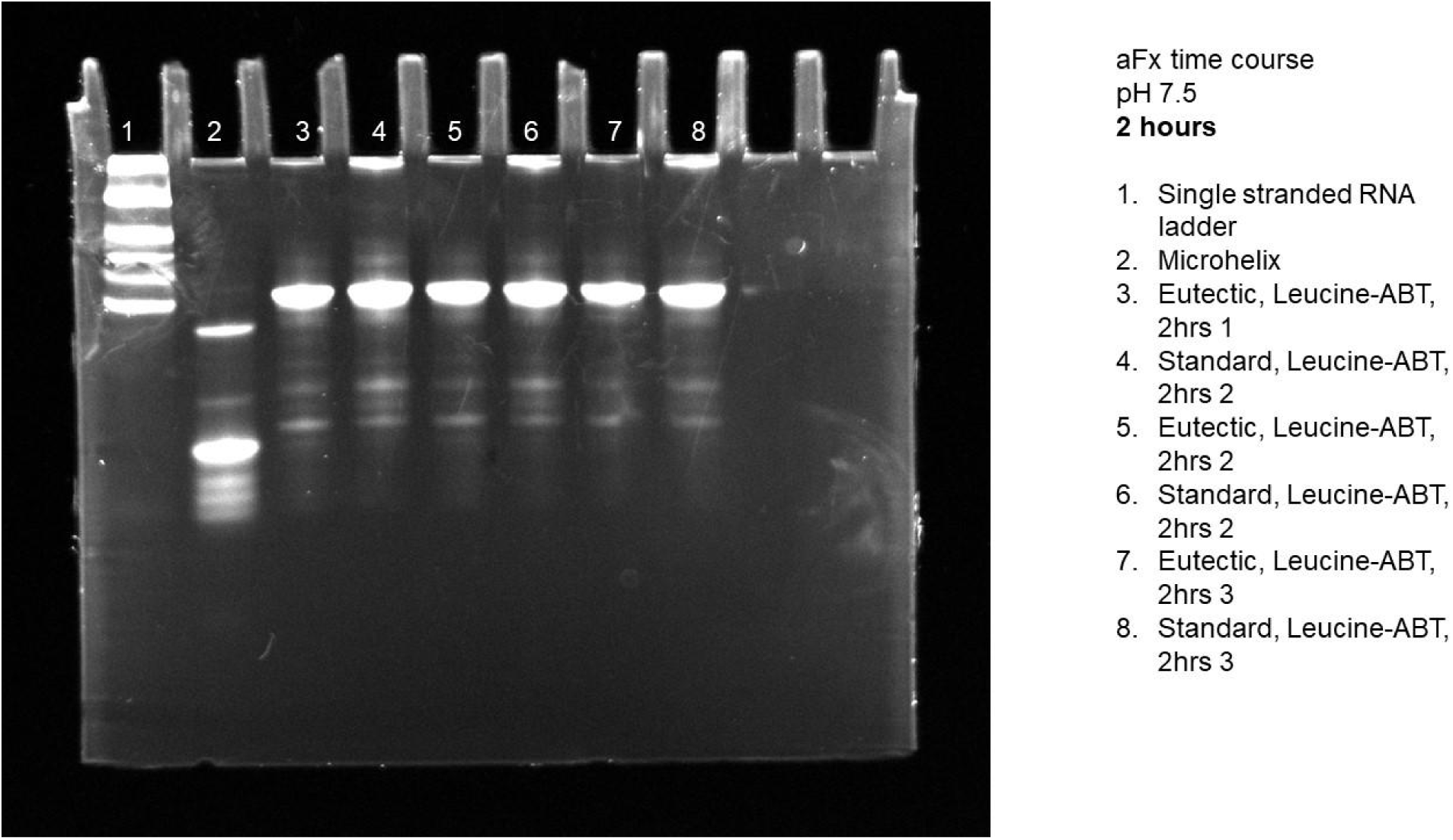
Gel of eutectic and standard reactions for aFx at pH 7.5 using L-Leu-ABT at 2 hours.

**Figure S17.**
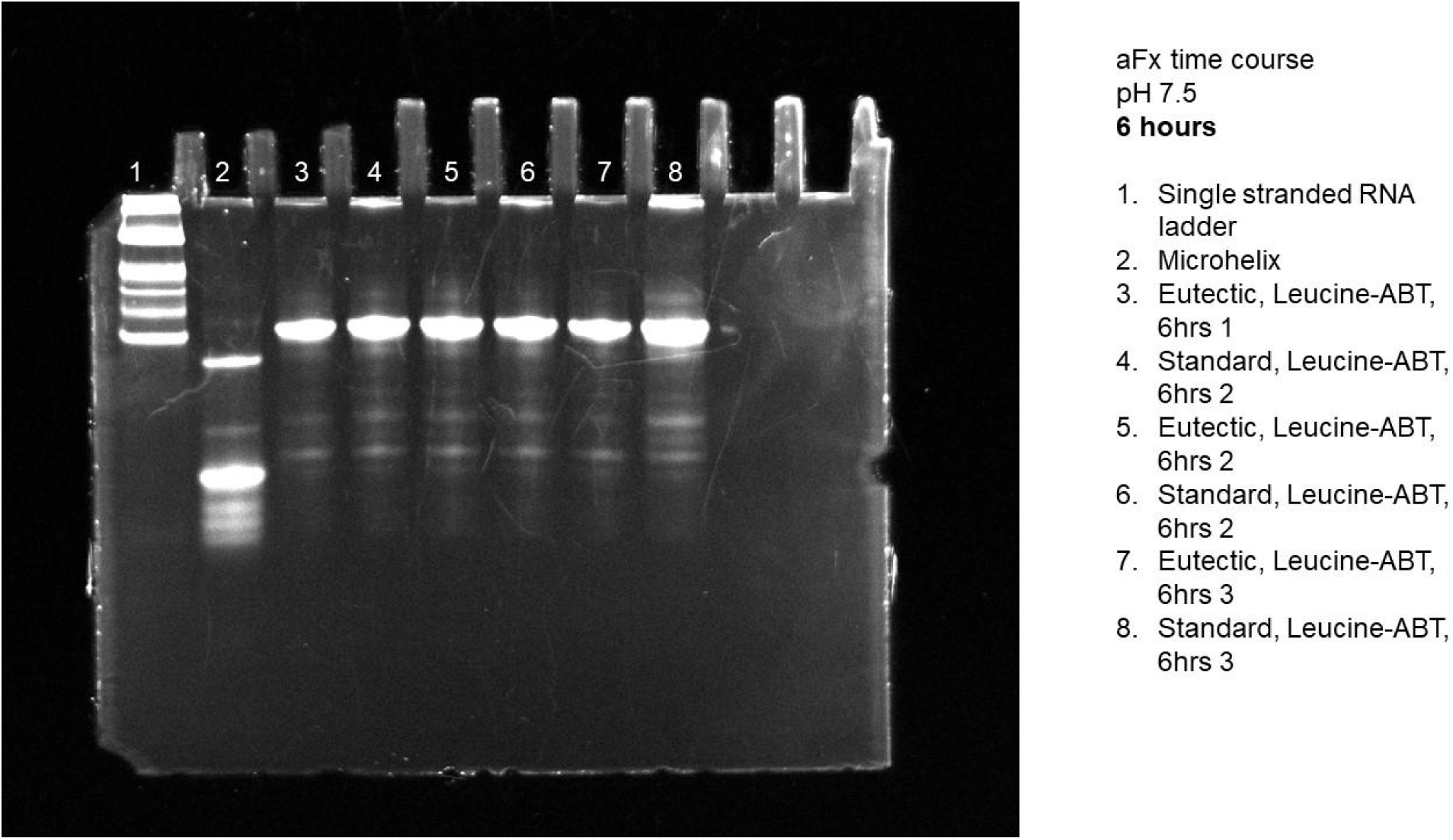
Gel of eutectic and standard reactions for aFx at pH 7.5 using L-Leu-ABT at 6 hours.

**Figure S18.**
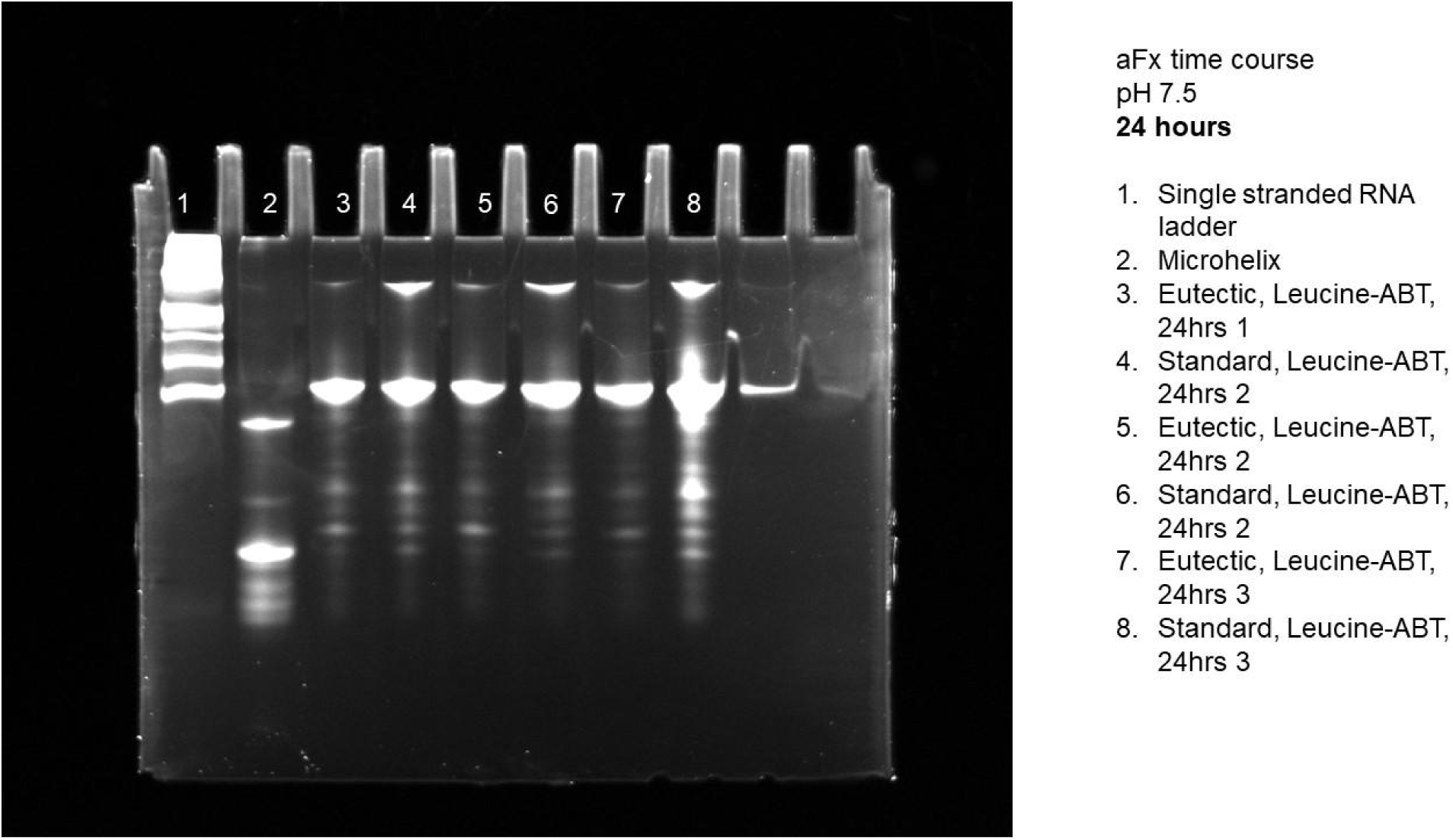
Gel of eutectic and standard reactions for aFx at pH 7.5 using L-Leu-ABT at 24 hours.

**Figure S19.**
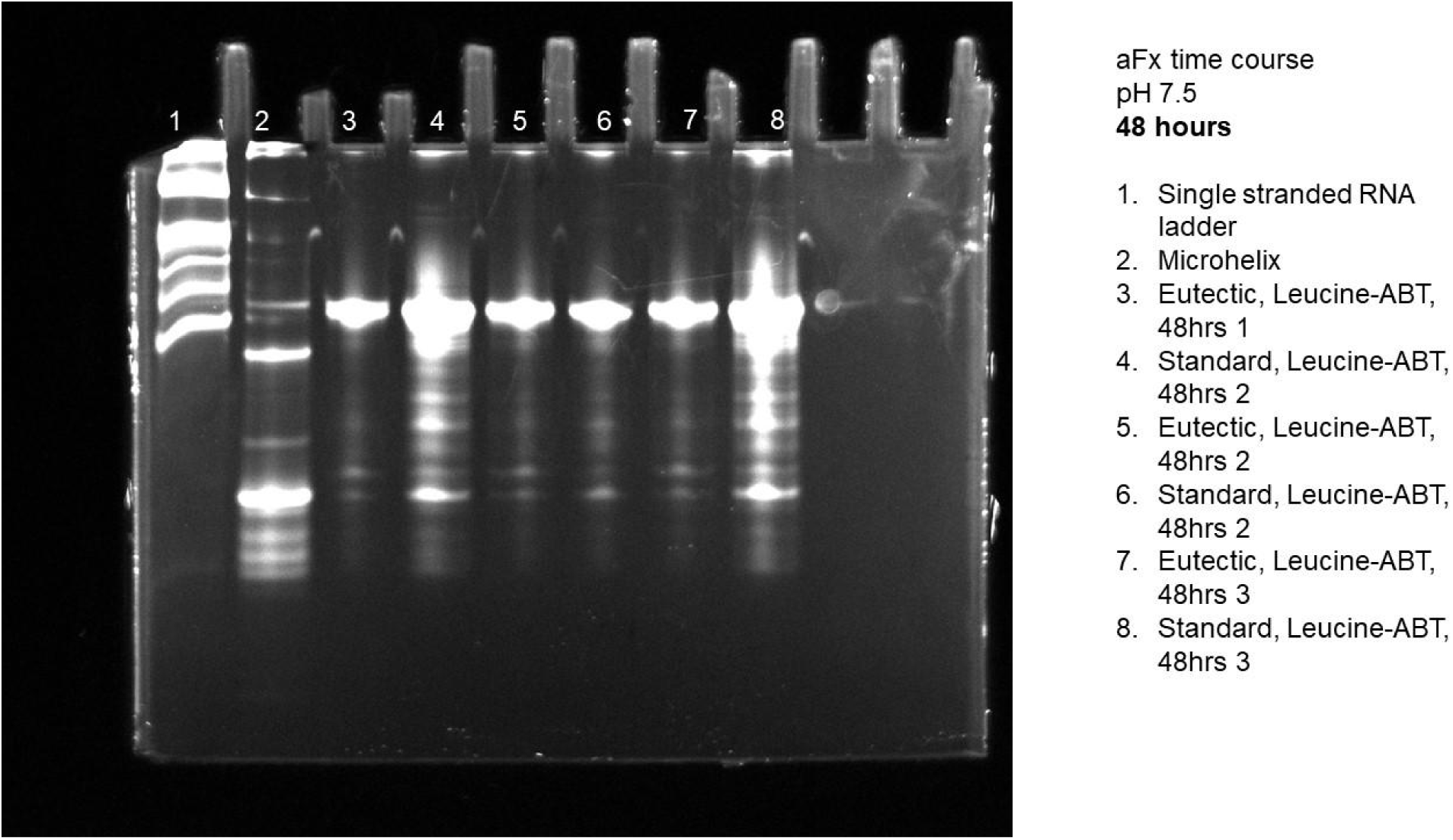
Gel of eutectic and standard reactions for aFx at pH 7.5 using L-Leu-ABT at 48 hours.

**Figure S20.**
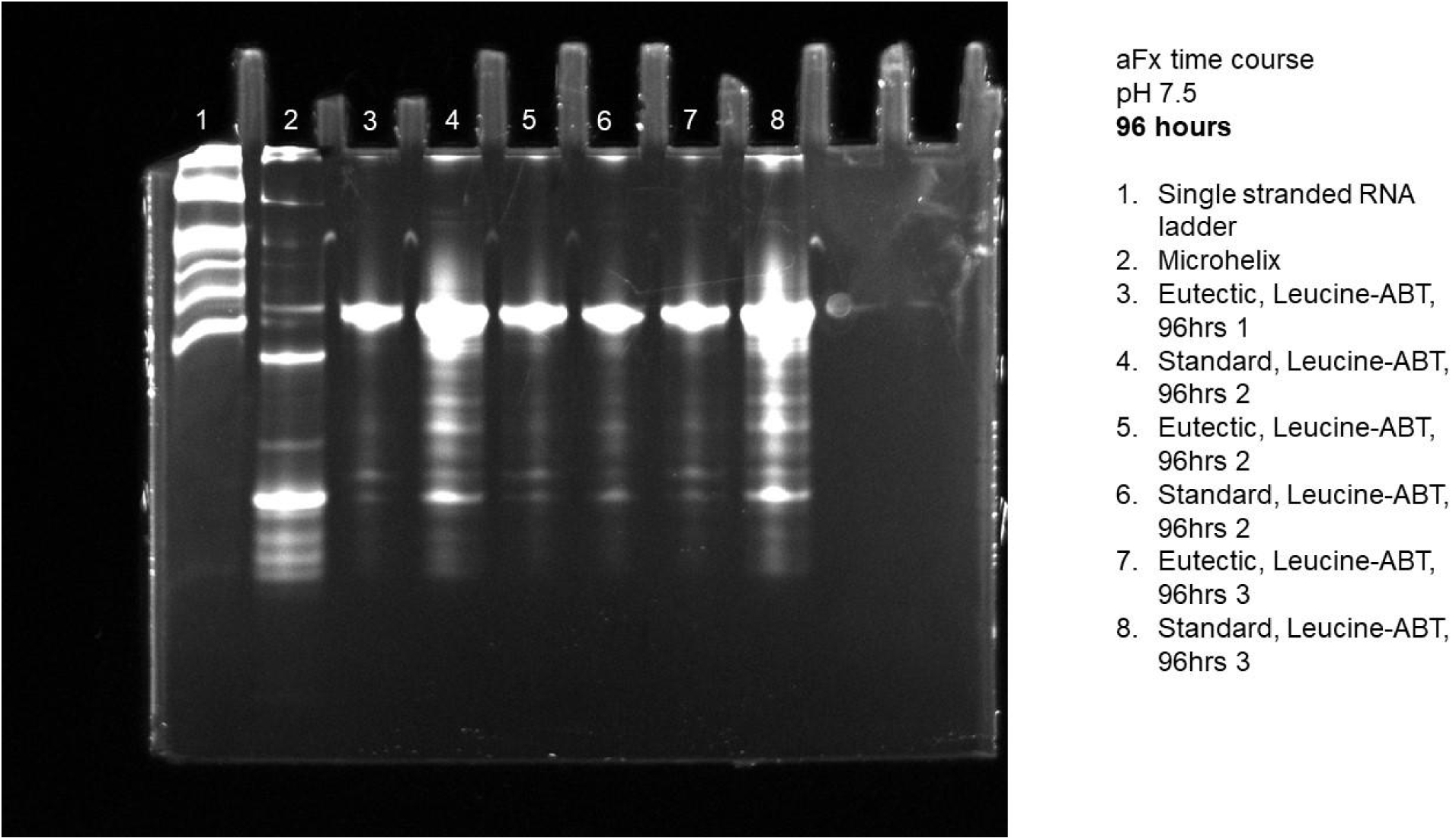
Gel of eutectic and standard reactions for aFx at pH 7.5 using L-Leu-ABT at 96 hours.

**Figure S21.**
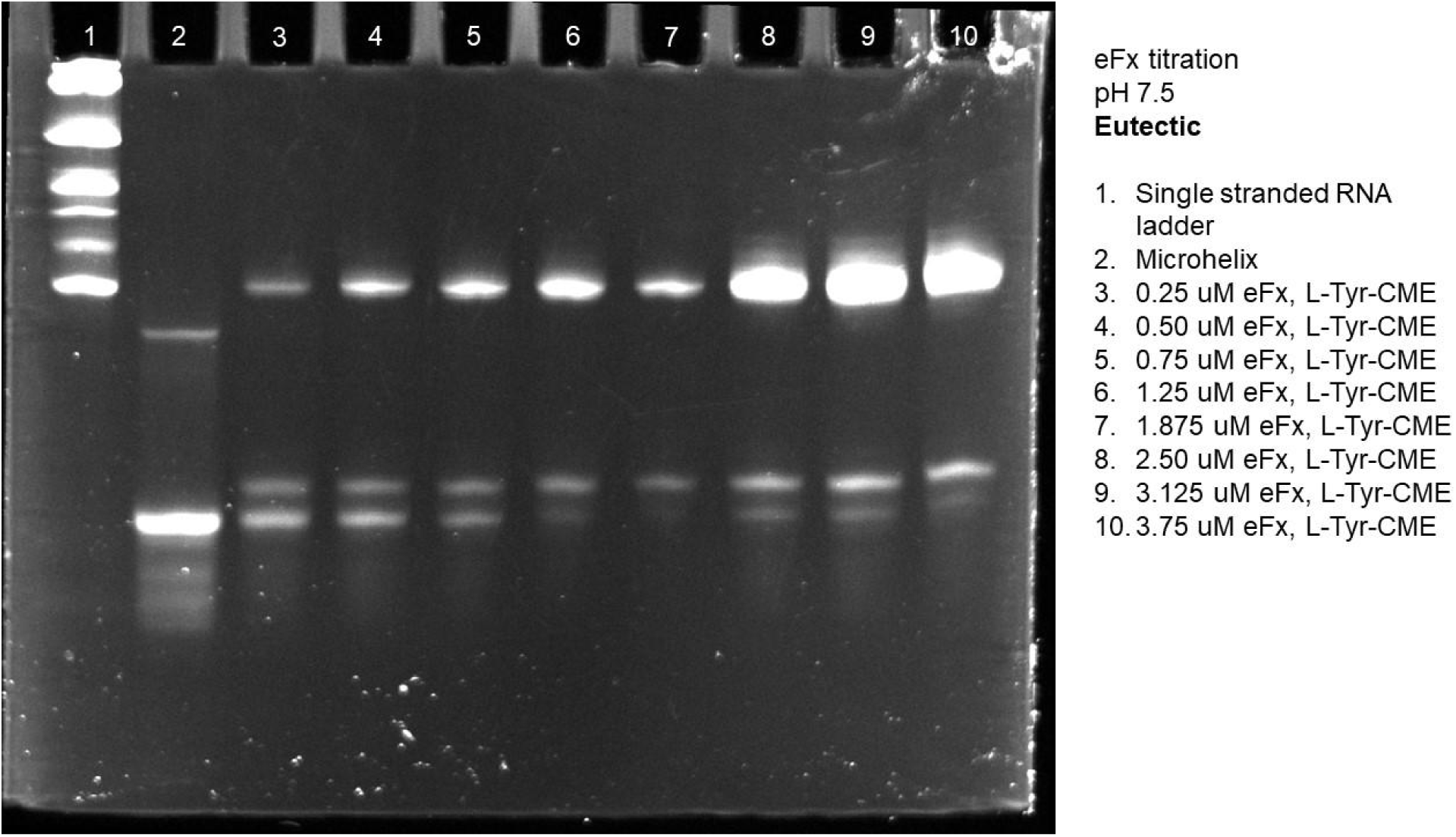
Gel of eutectic eFx titration at pH 7.5 using L-Tyr-CME.

**Figure S22.**
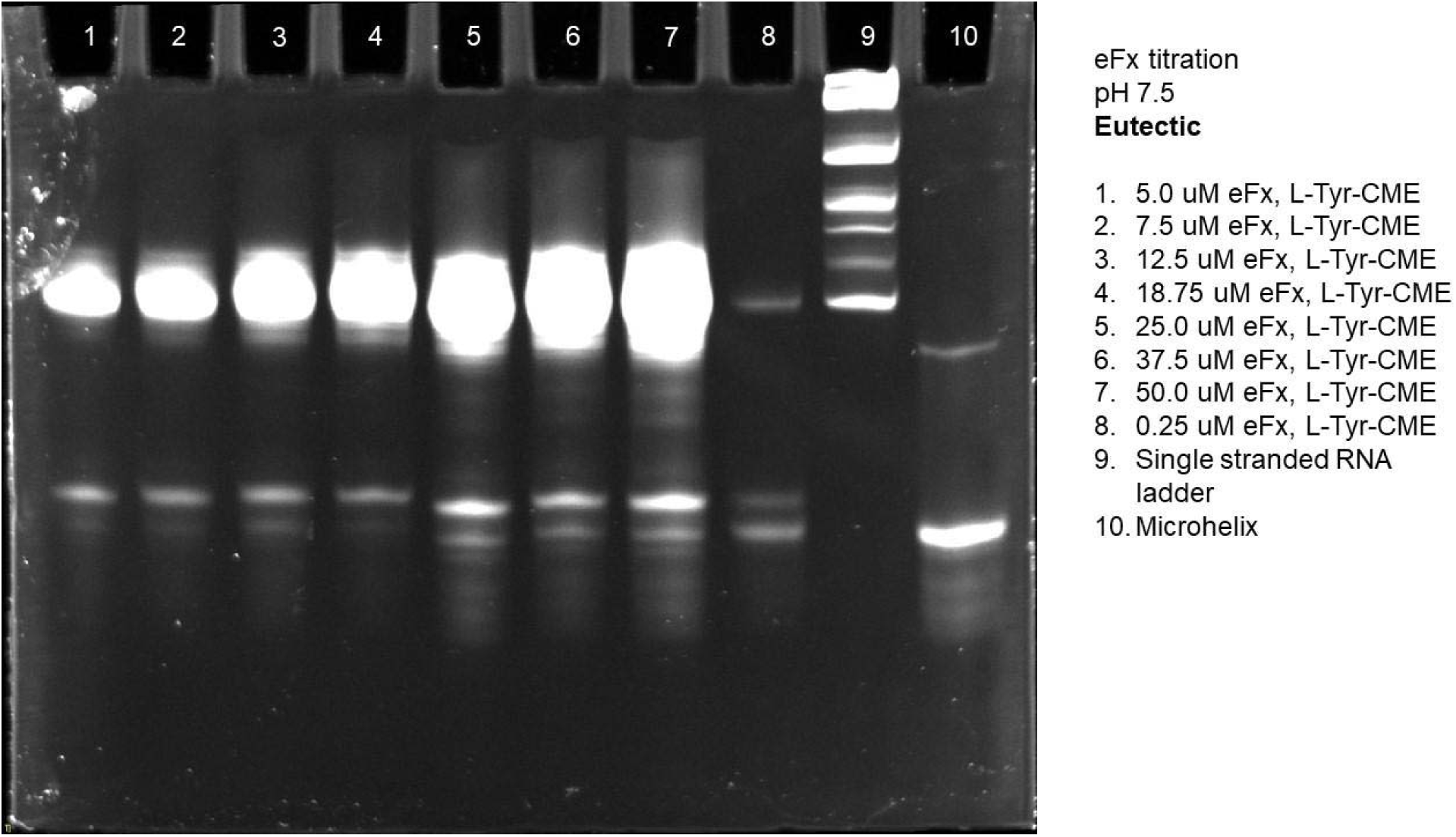
Gel of eutectic eFx titration at pH 7.5 using L-Tyr-CME.

**Figure S23.**
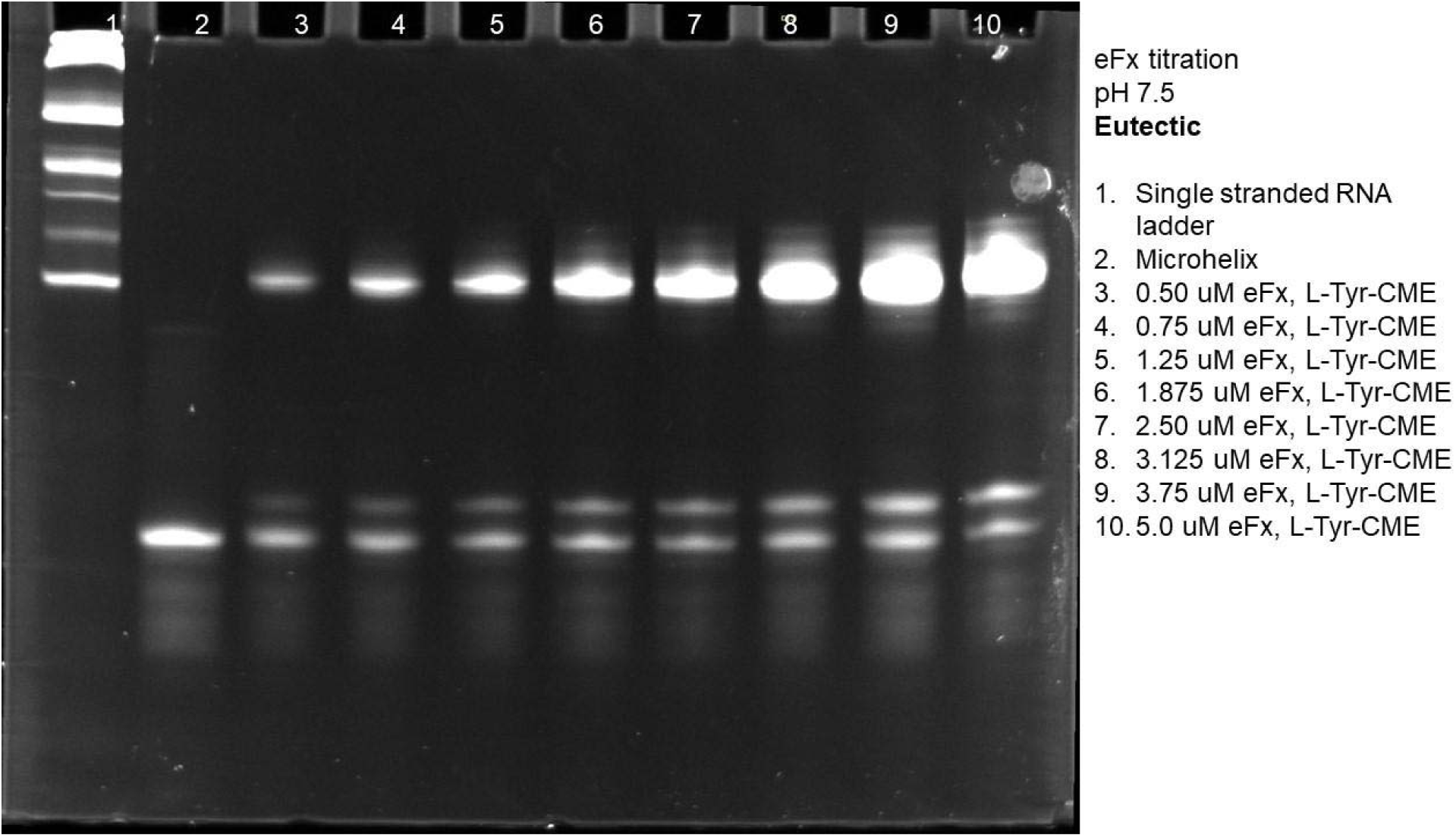
Gel of eutectic eFx titration at pH 7.5 using L-Tyr-CME.

**Figure S24.**
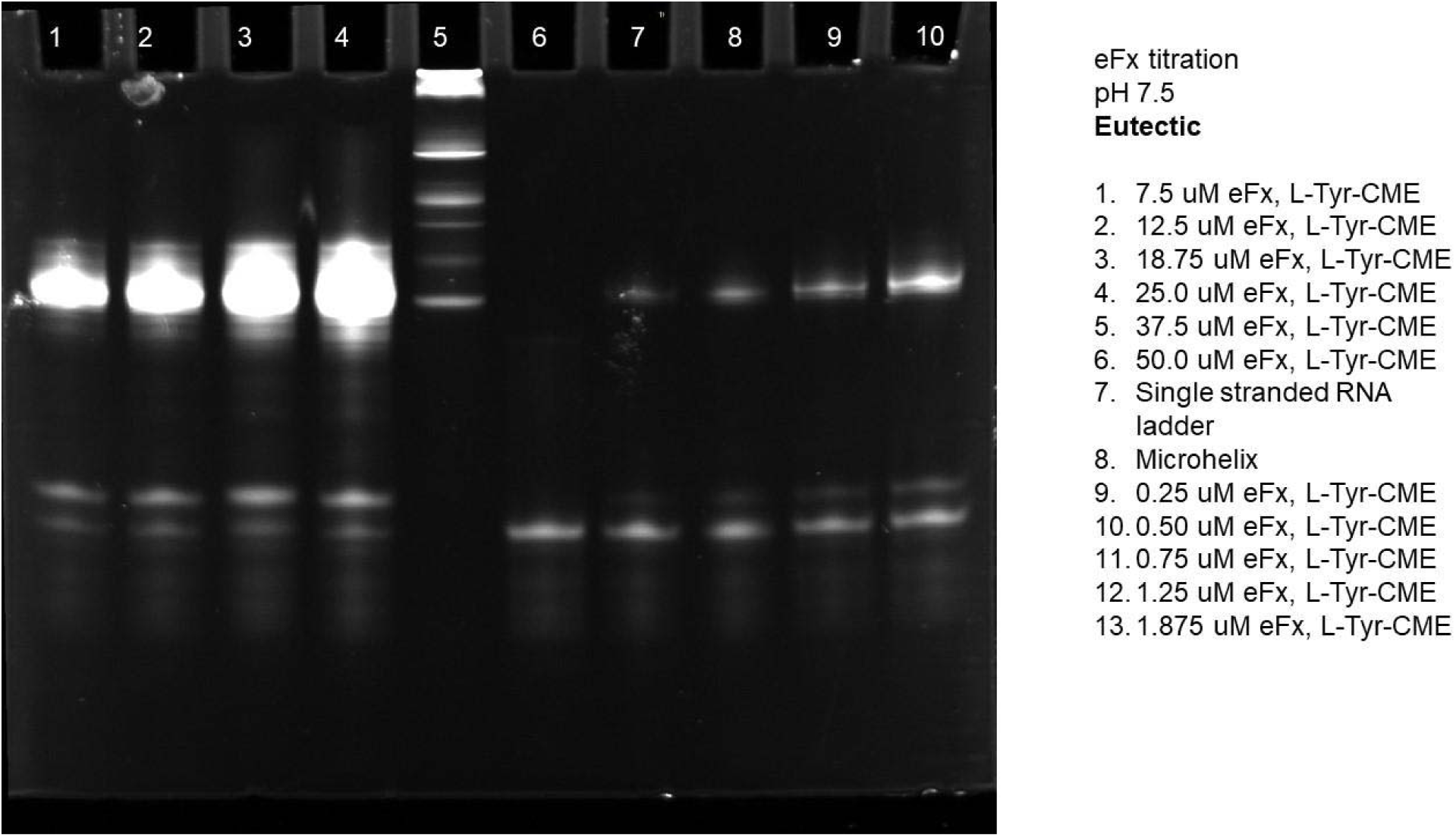
Gel of eutectic eFx titration at pH 7.5 using L-Tyr-CME.

**Figure S25.**
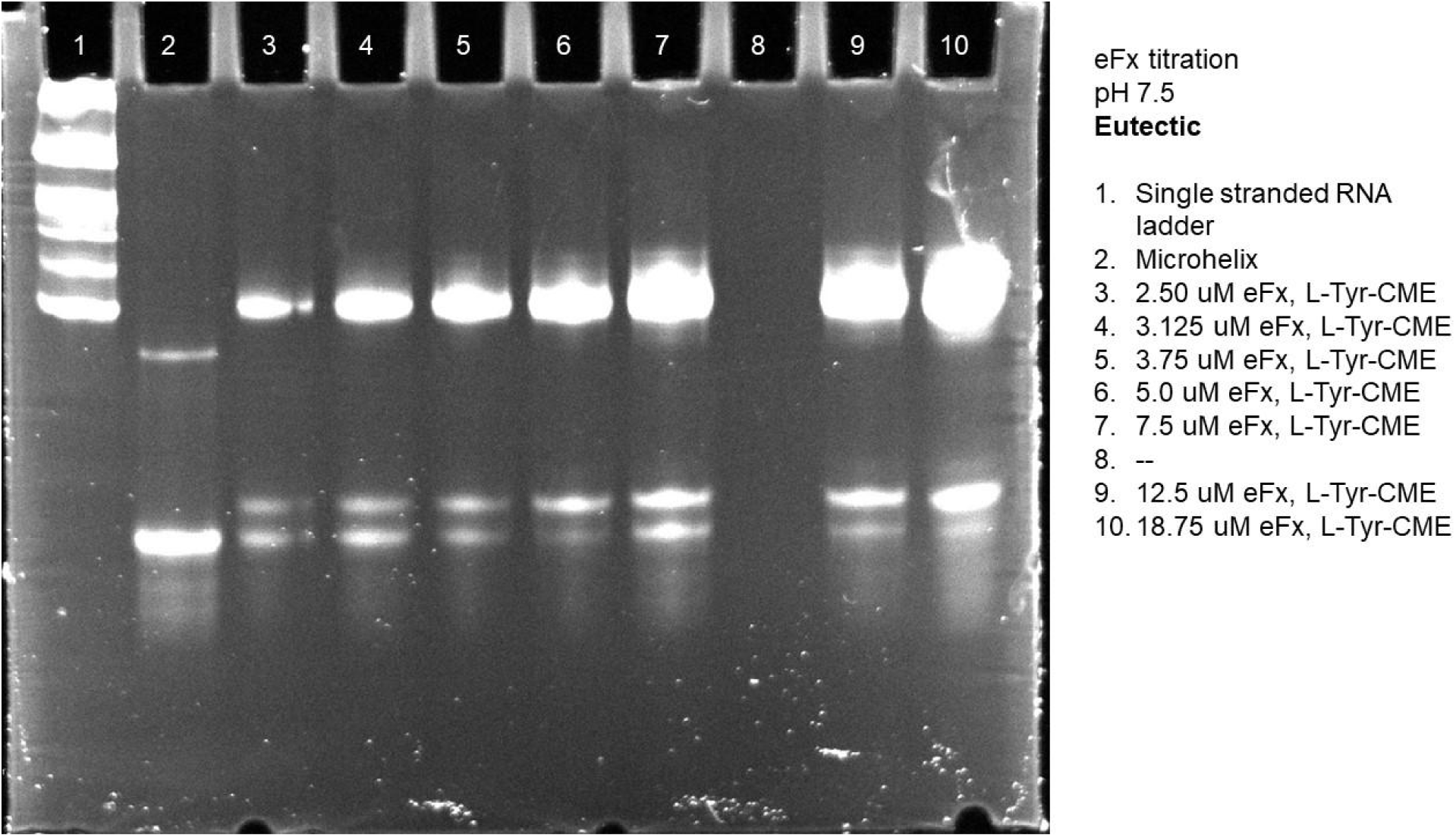
Gel of eutectic eFx titration at pH 7.5 using L-Tyr-CME.

**Figure S26.**
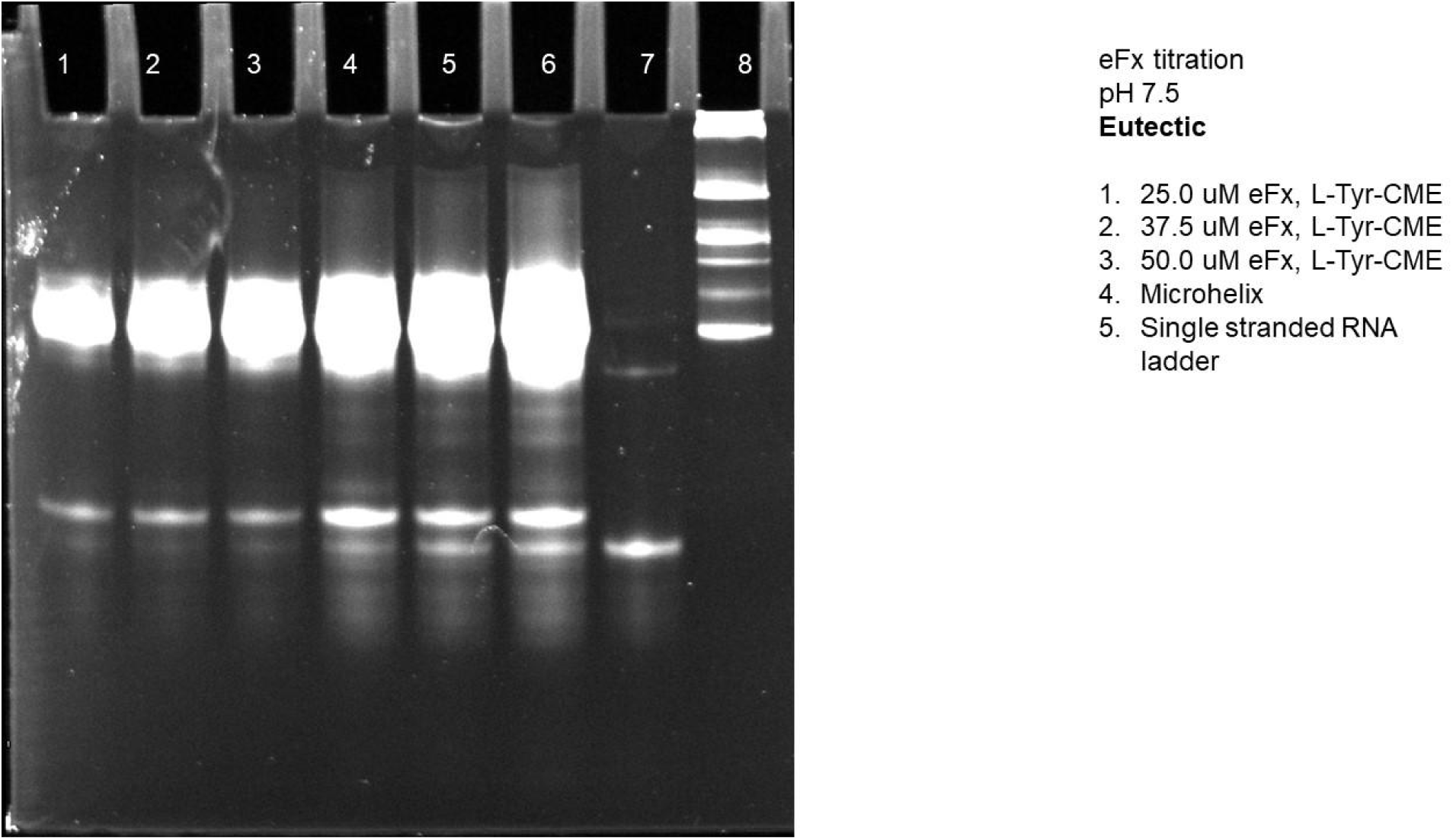
Gel of eutectic eFx titration at pH 7.5 using L-Tyr-CME.

**Figure S27.**
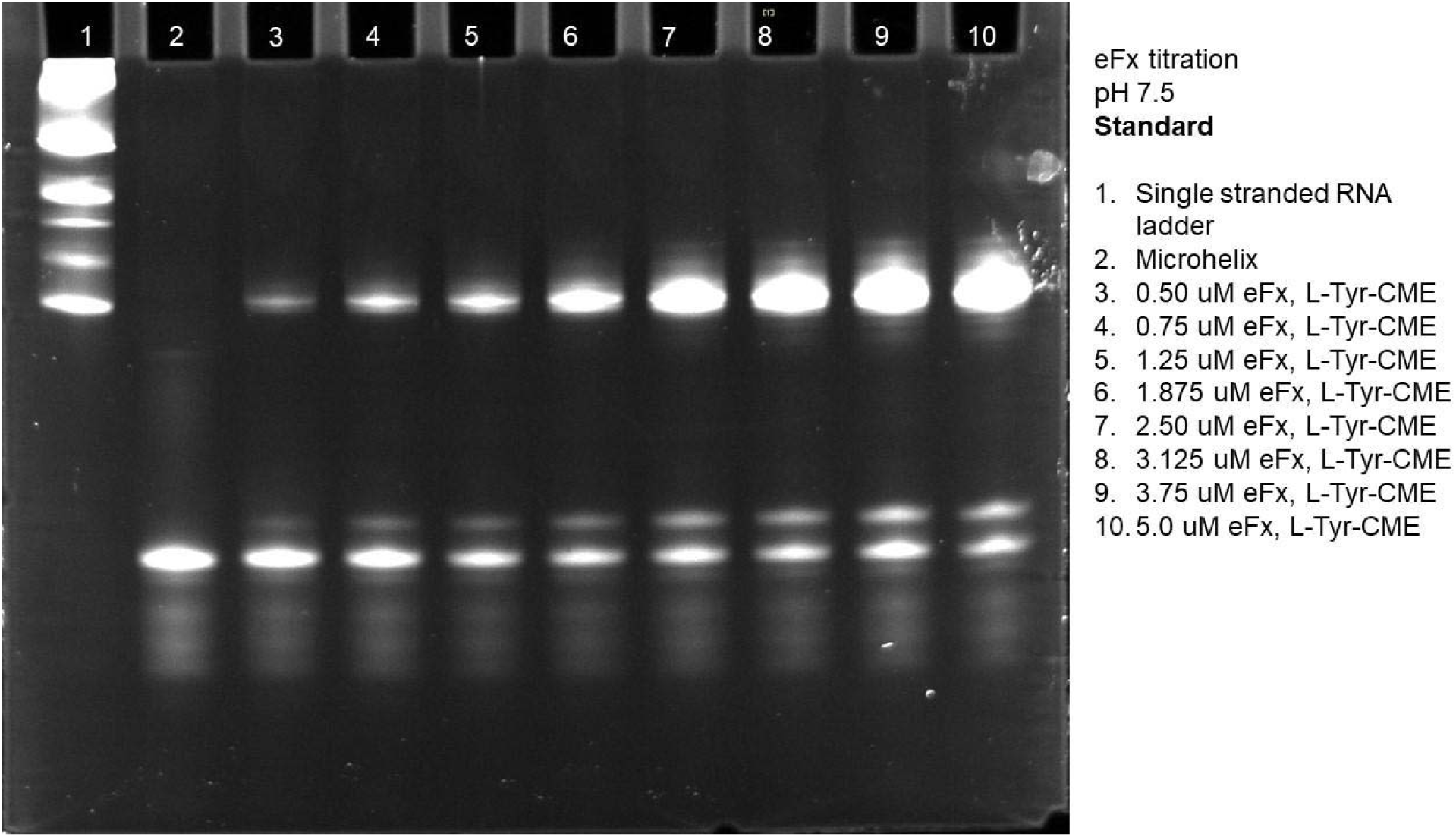
Gel of standard eFx titration at pH 7.5 using L-Tyr-CME.

**Figure S28.**
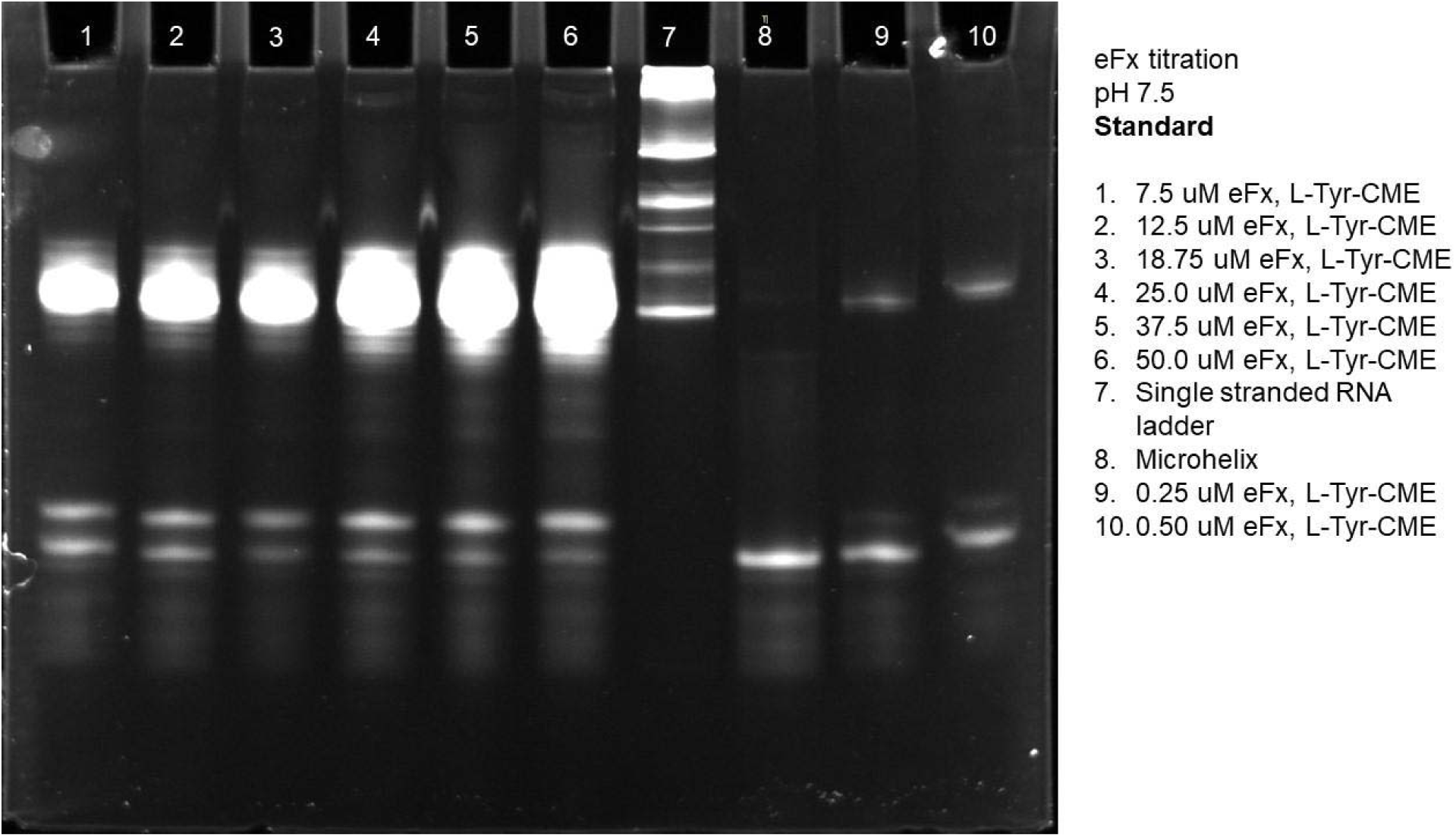
Gel of standard eFx titration at pH 7.5 using L-Tyr-CME.

**Figure S29.**
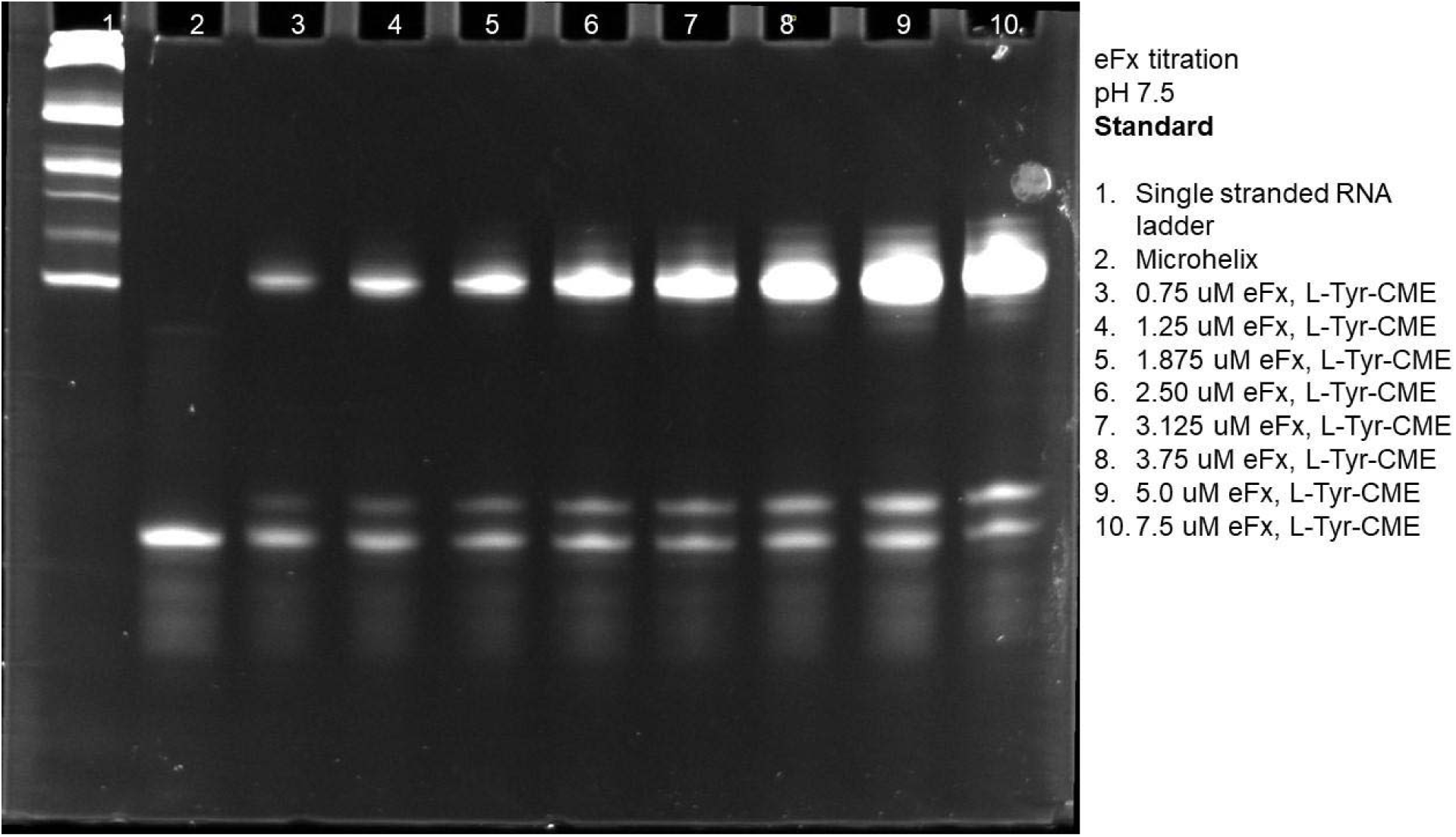
Gel of standard eFx titration at pH 7.5 using L-Tyr-CME.

**Figure S30.**
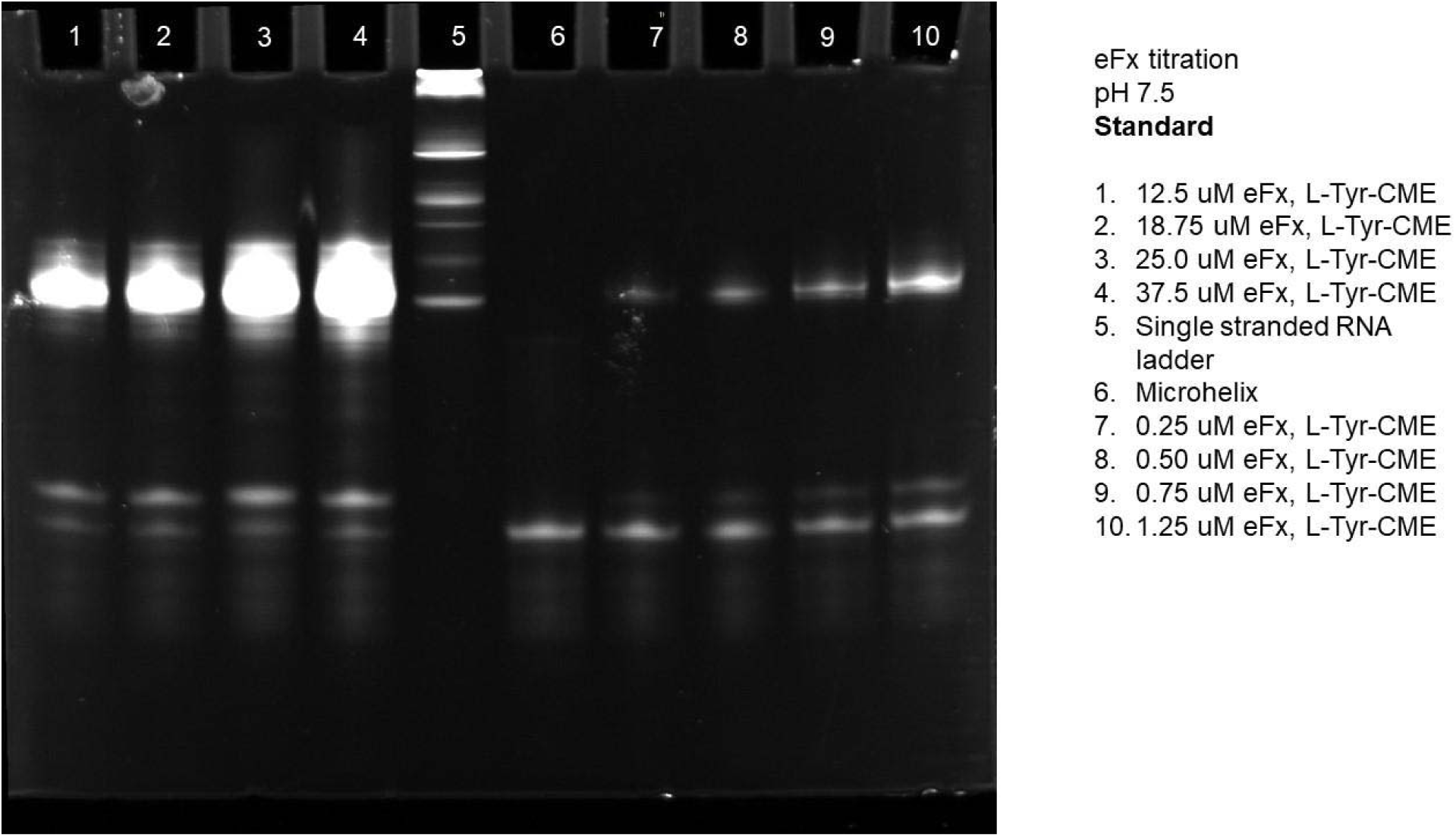
Gel of standard eFx titration at pH 7.5 using L-Tyr-CME.

**Figure S31.**
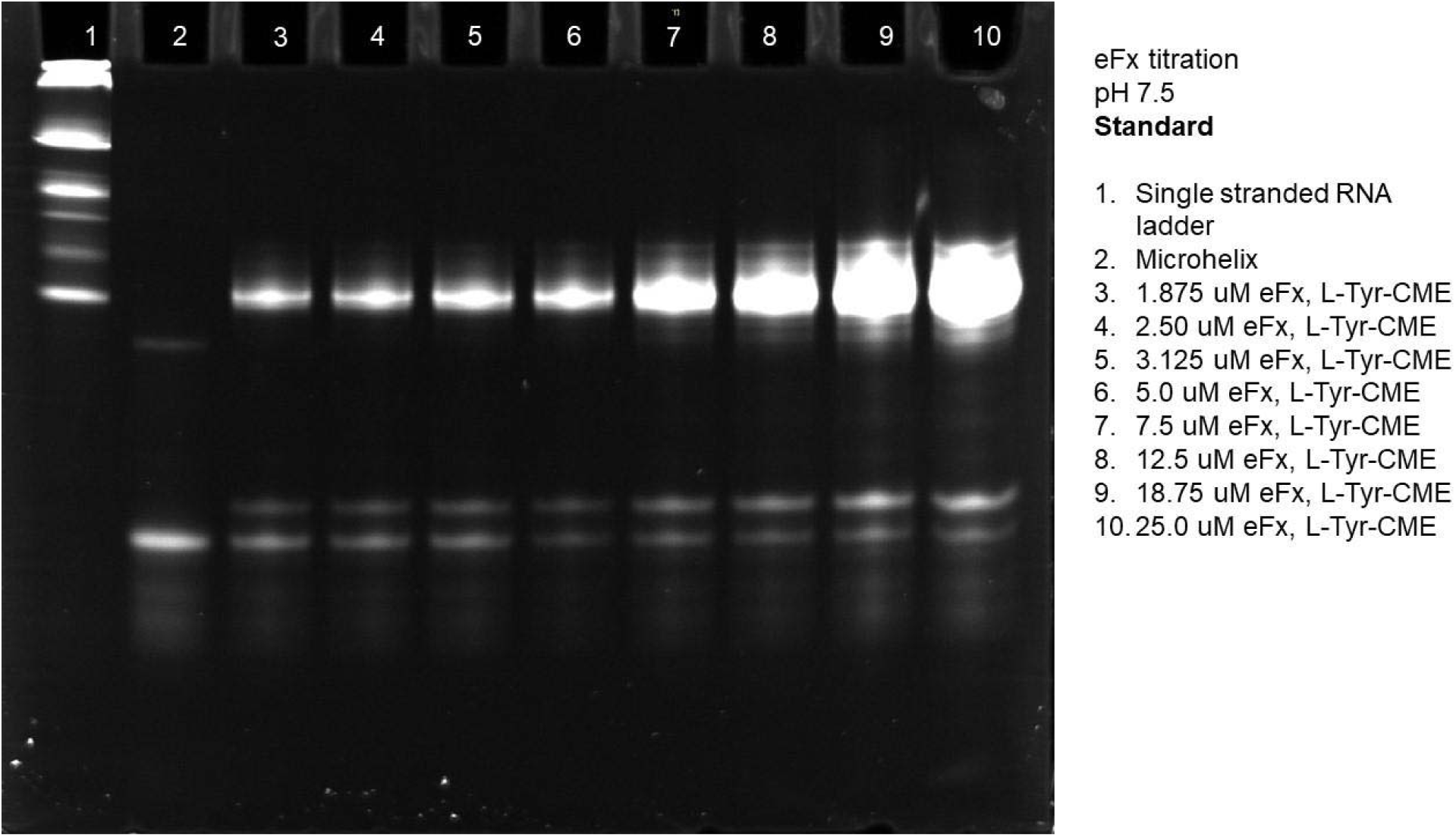
Gel of standard eFx titration at pH 7.5 using L-Tyr-CME.

**Figure S32.**
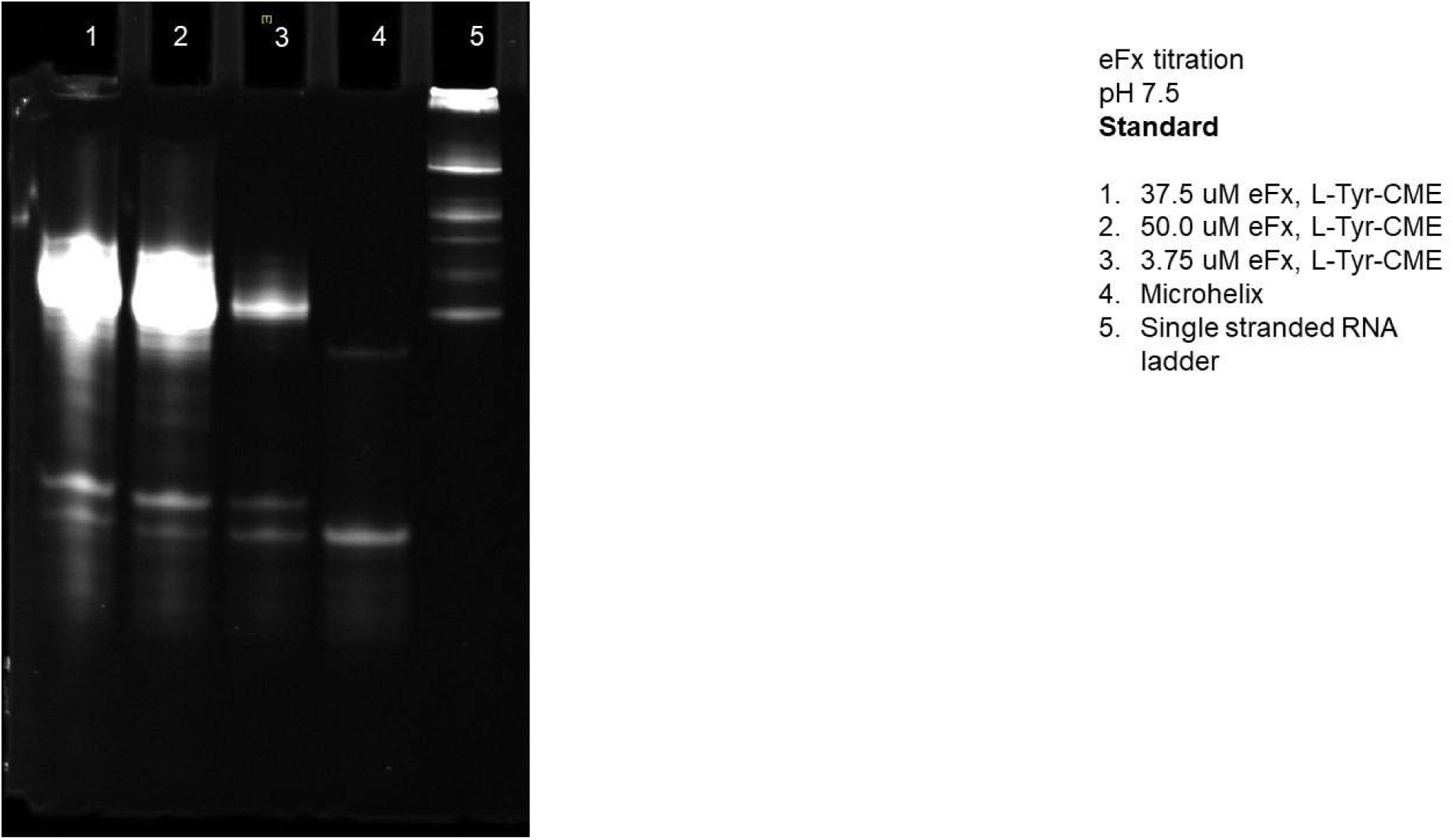
Gel of standard eFx titration at pH 7.5 using L-Tyr-CME.

**Figure S33.**
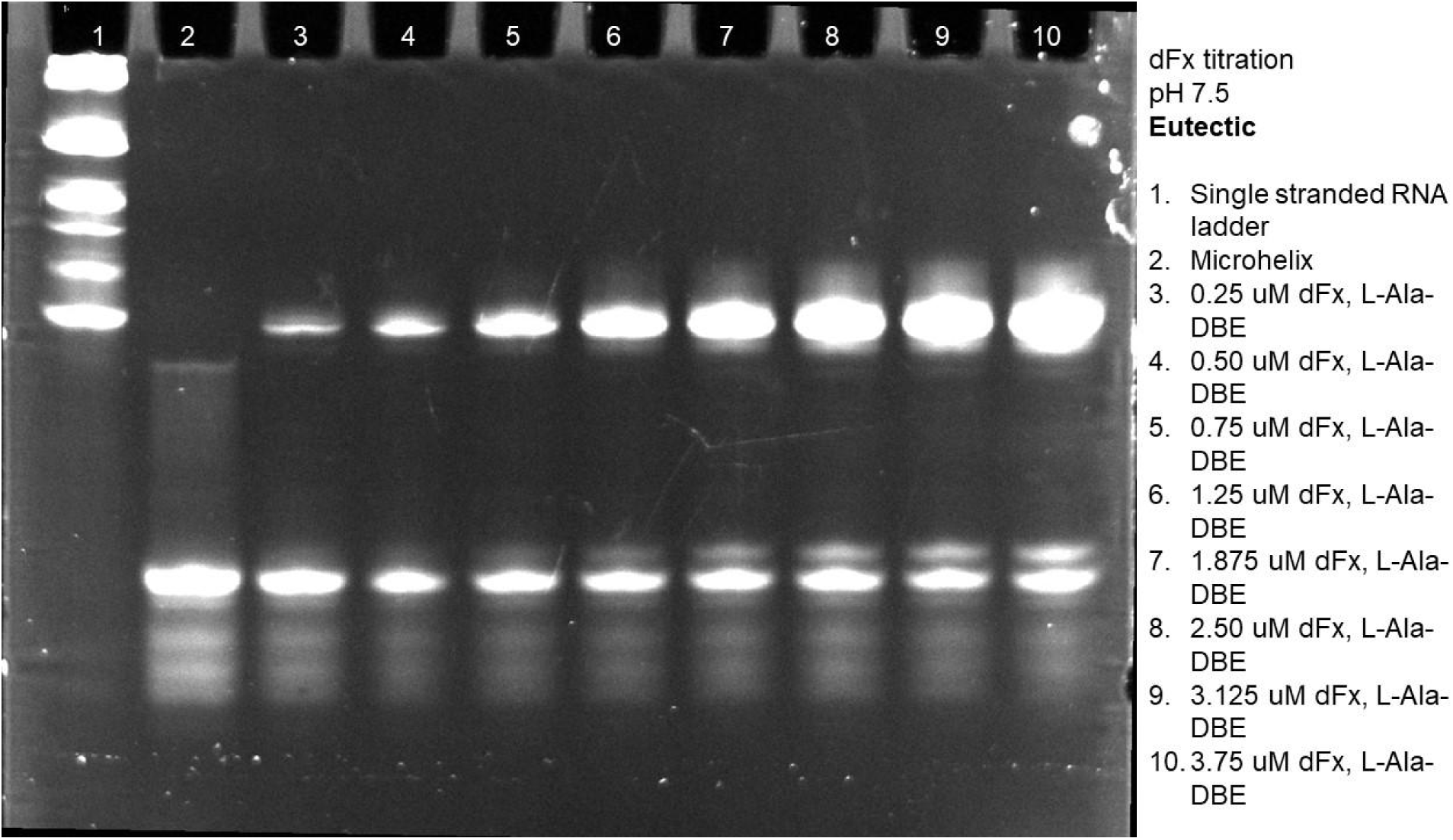
Gel of eutectic dFx titration at pH 7.5 using L-Ala-DBE.

**Figure S34.**
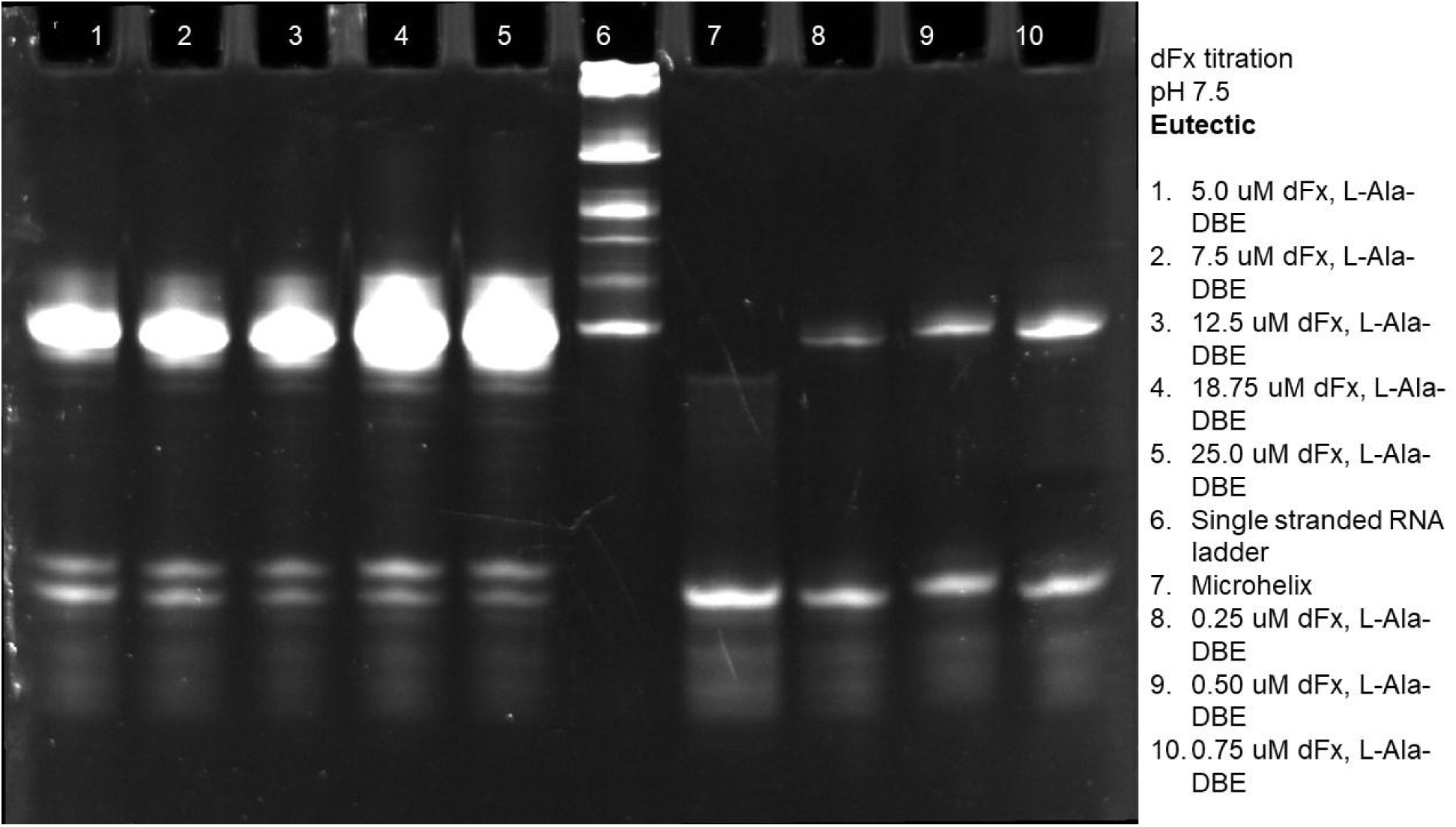
Gel of eutectic dFx titration at pH 7.5 using L-Ala-DBE.

**Figure S35.**
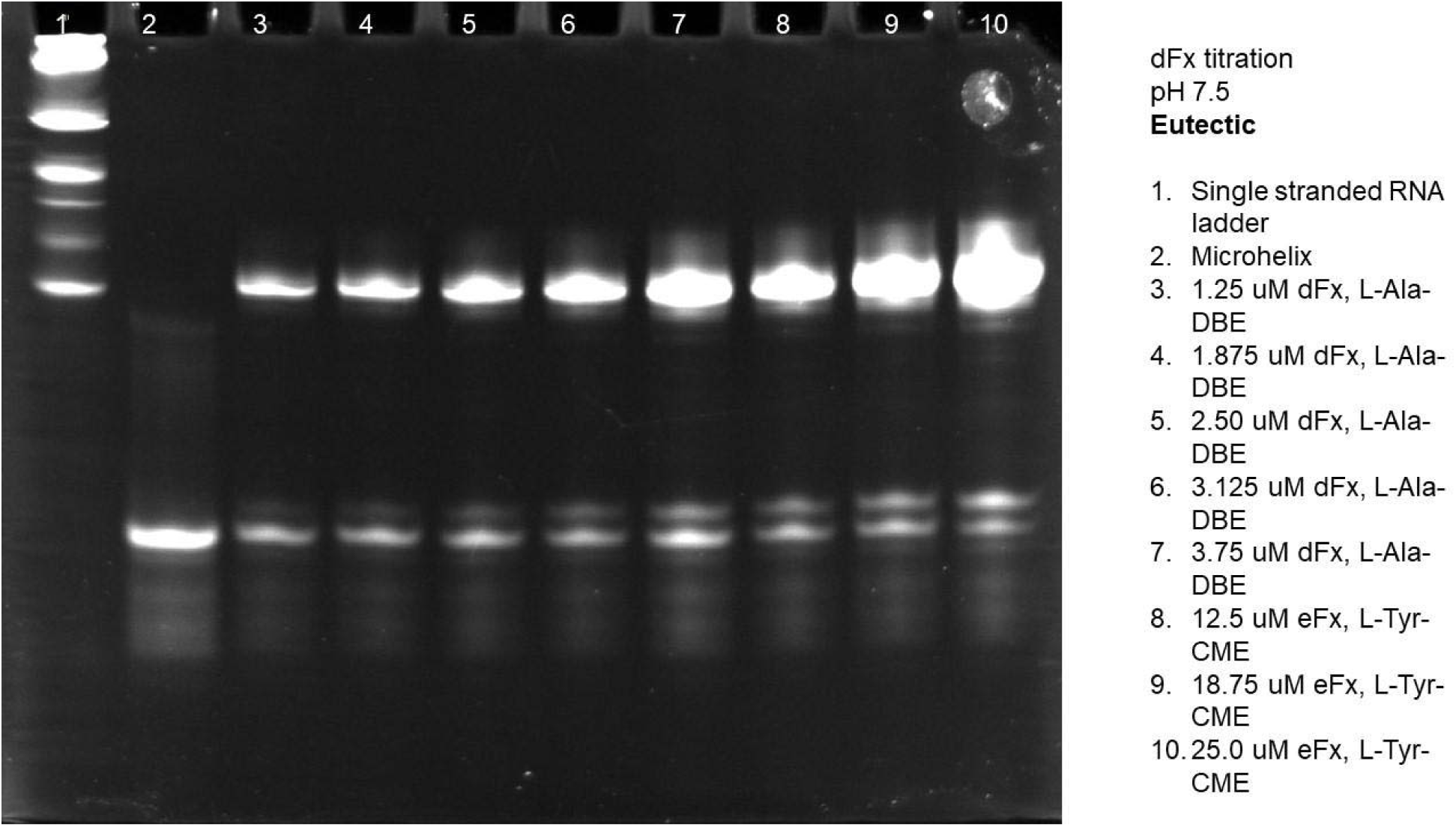
Gel of eutectic dFx titration at pH 7.5 using L-Ala-DBE.

**Figure S36.**
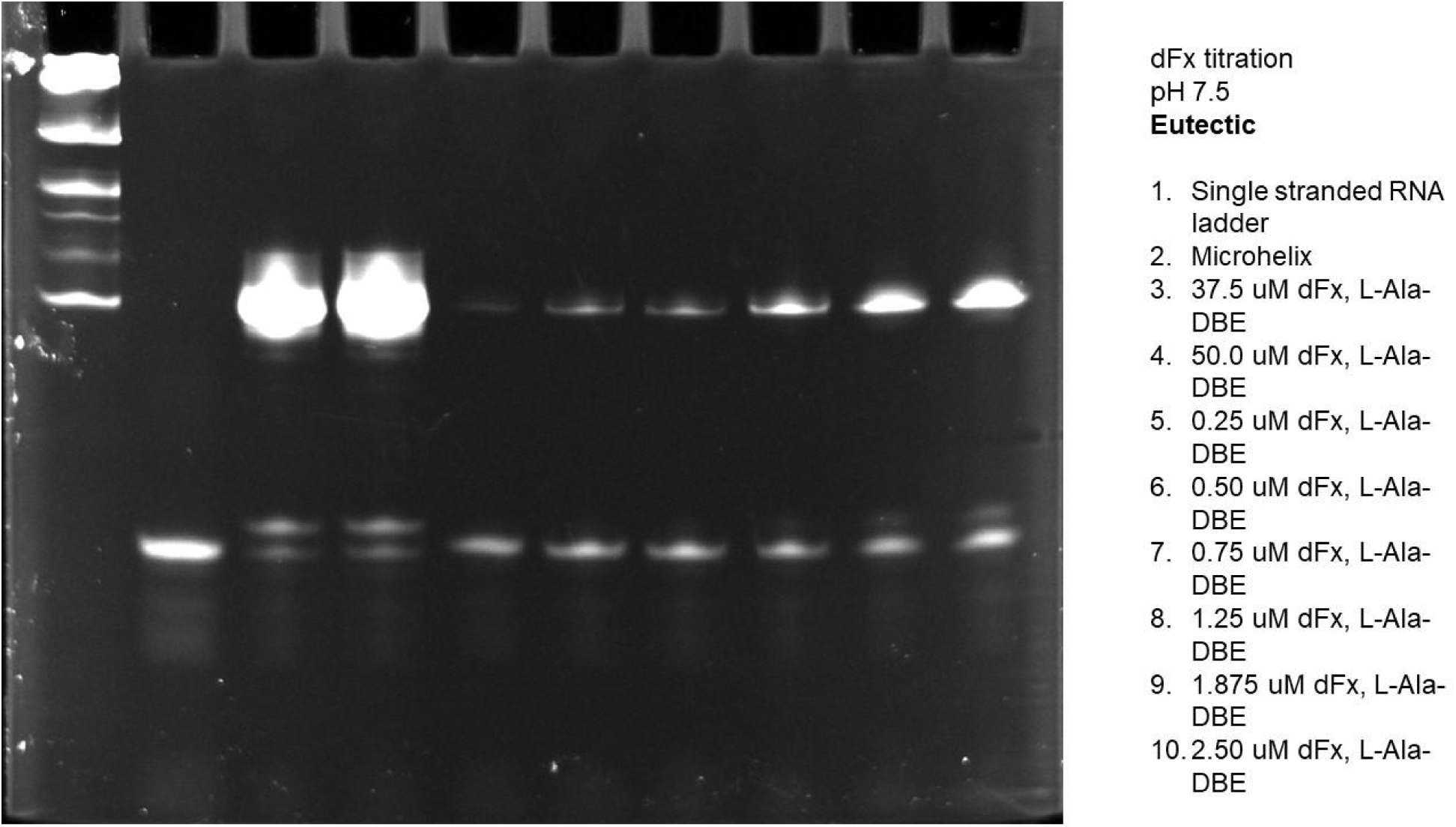
Gel of eutectic dFx titration at pH 7.5 using L-Ala-DBE.

**Figure S37.**
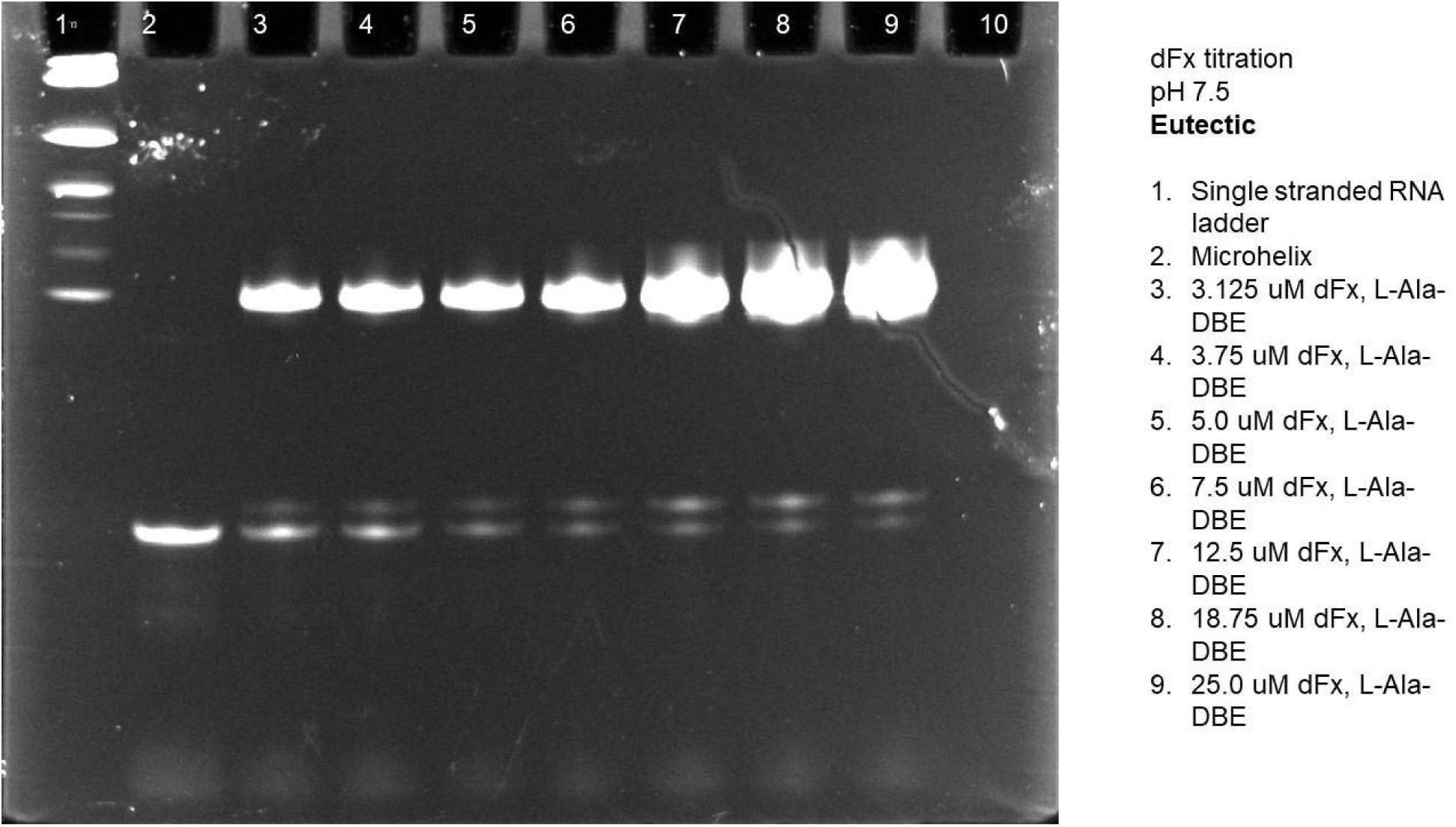
Gel of eutectic dFx titration at pH 7.5 using L-Ala-DBE.

**Figure S38.**
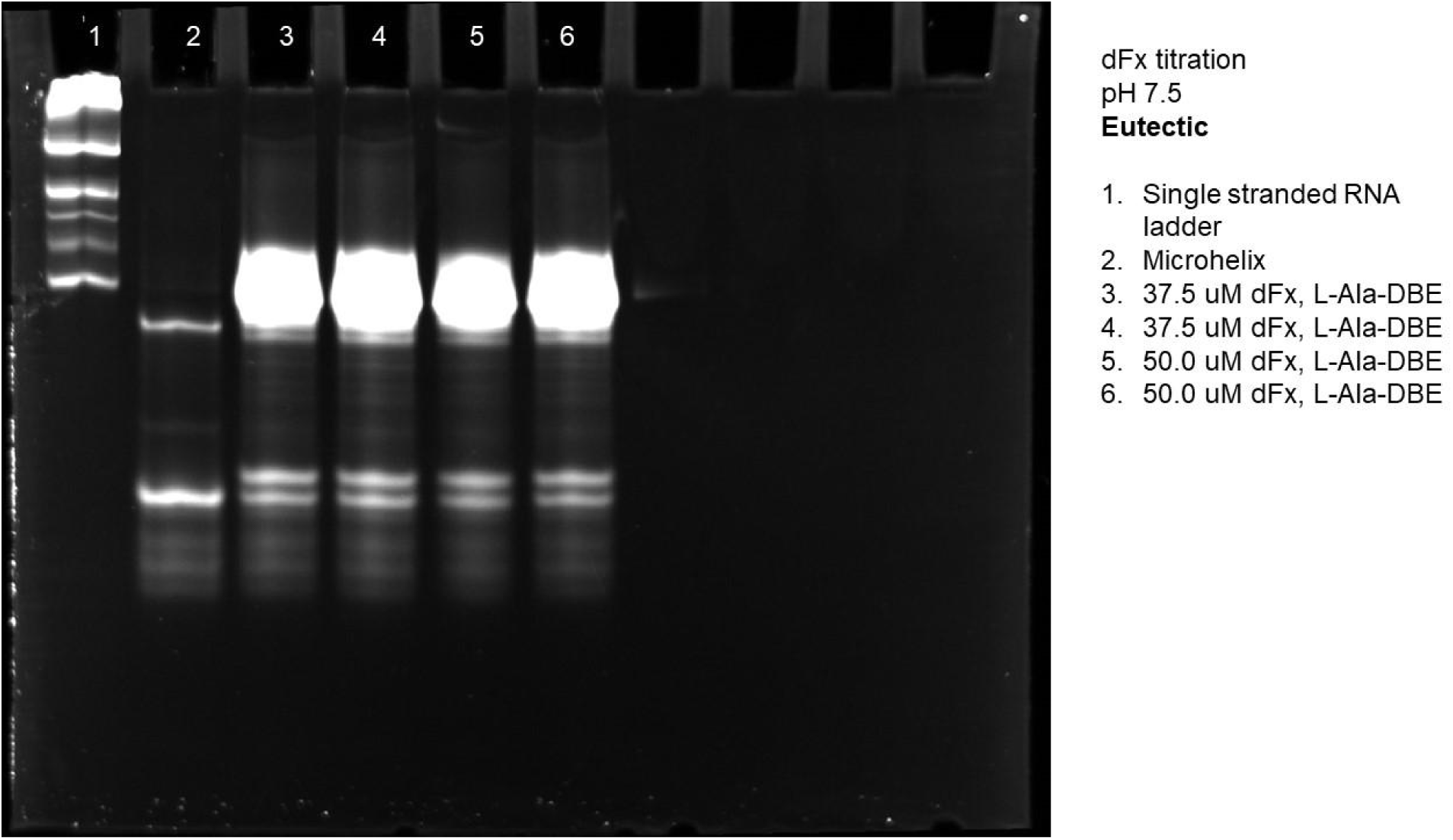
Gel of eutectic dFx titration at pH 7.5 using L-Ala-DBE.

**Figure S39.**
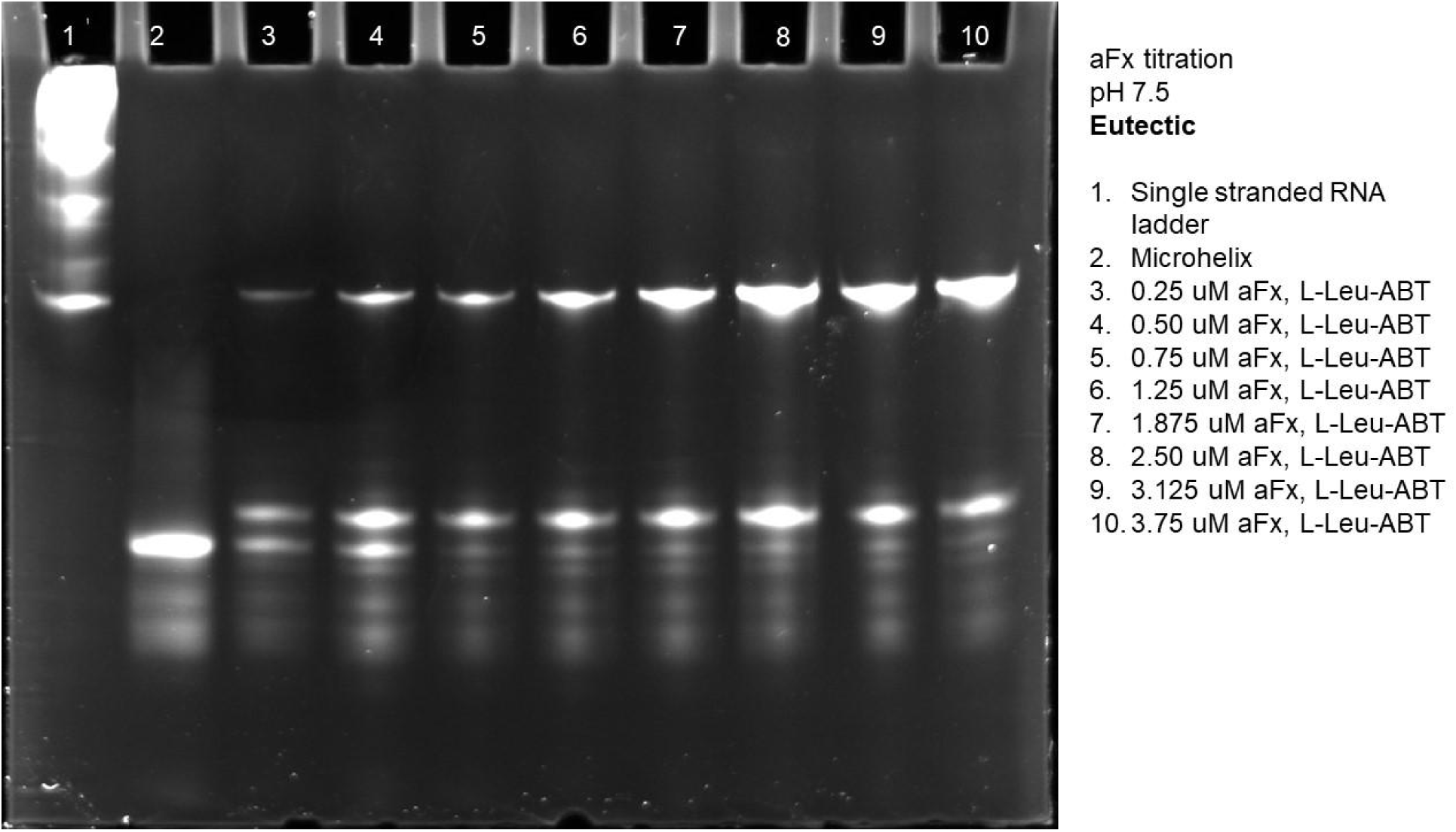
Gel of eutectic aFx titration at pH 7.5 using L-Leu-ABT.

**Figure S40.**
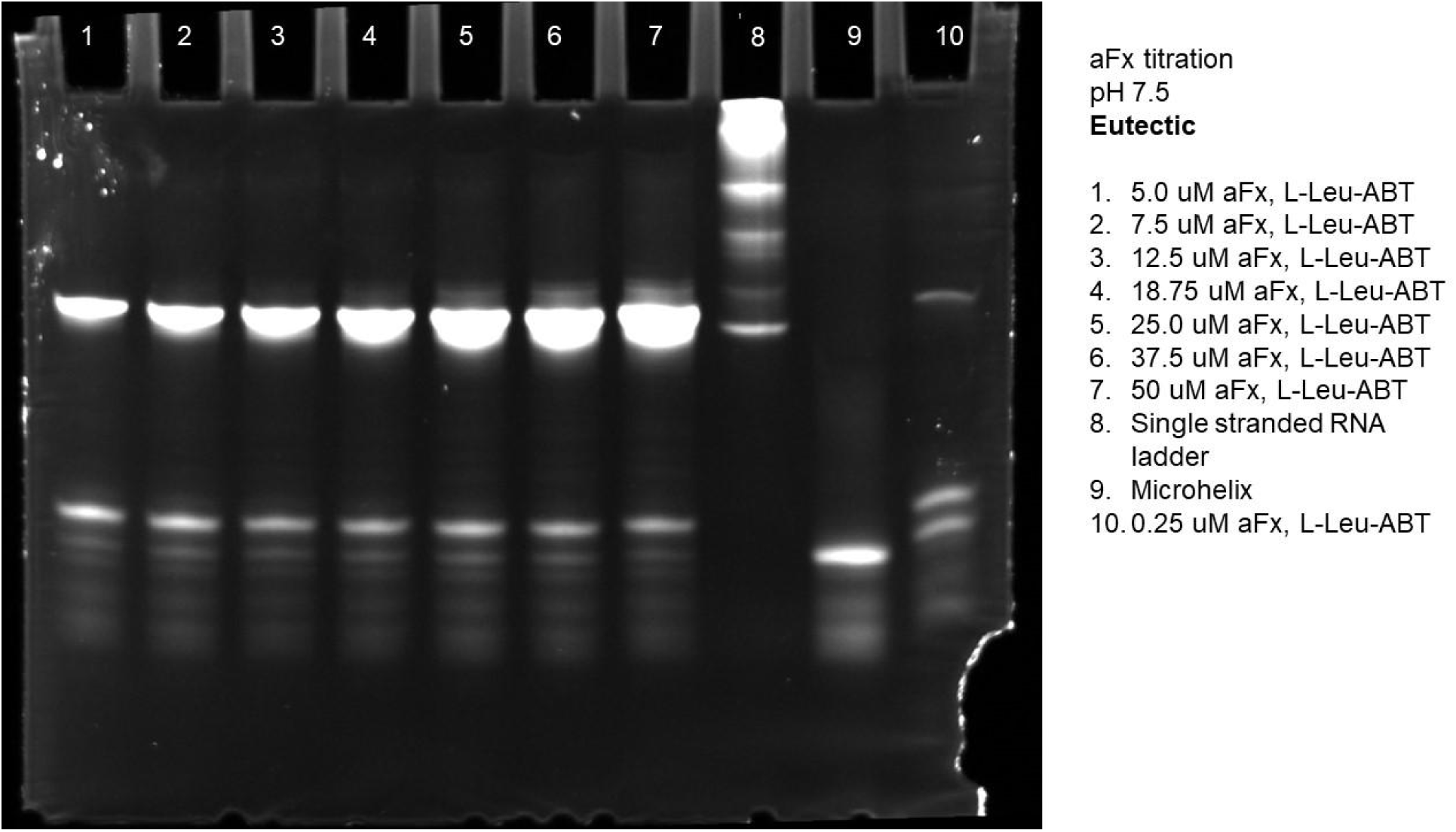
Gel of eutectic aFx titration at pH 7.5 using L-Leu-ABT.

**Figure S41.**
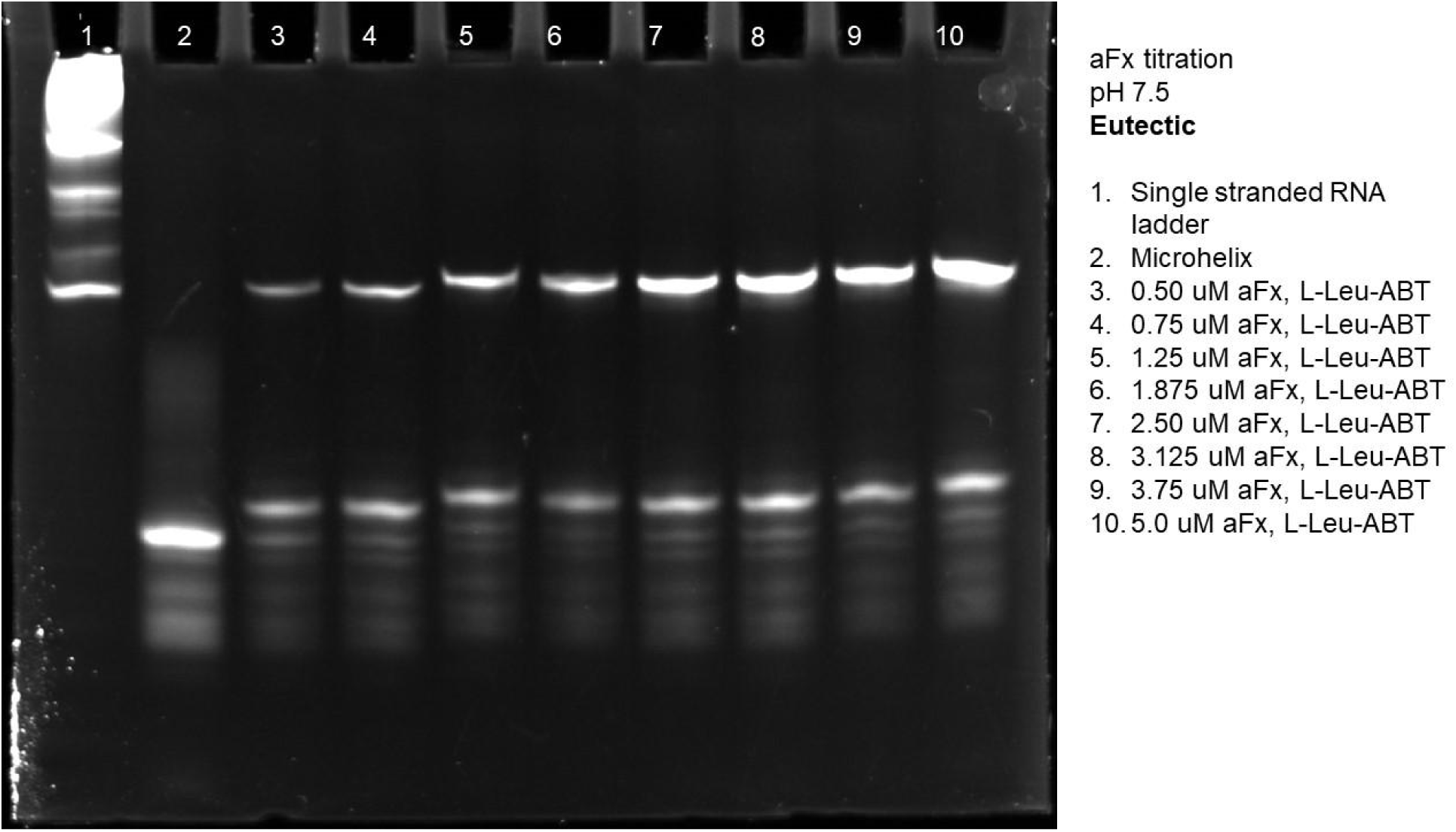
Gel of eutectic aFx titration at pH 7.5 using L-Leu-ABT.

**Figure S42.**
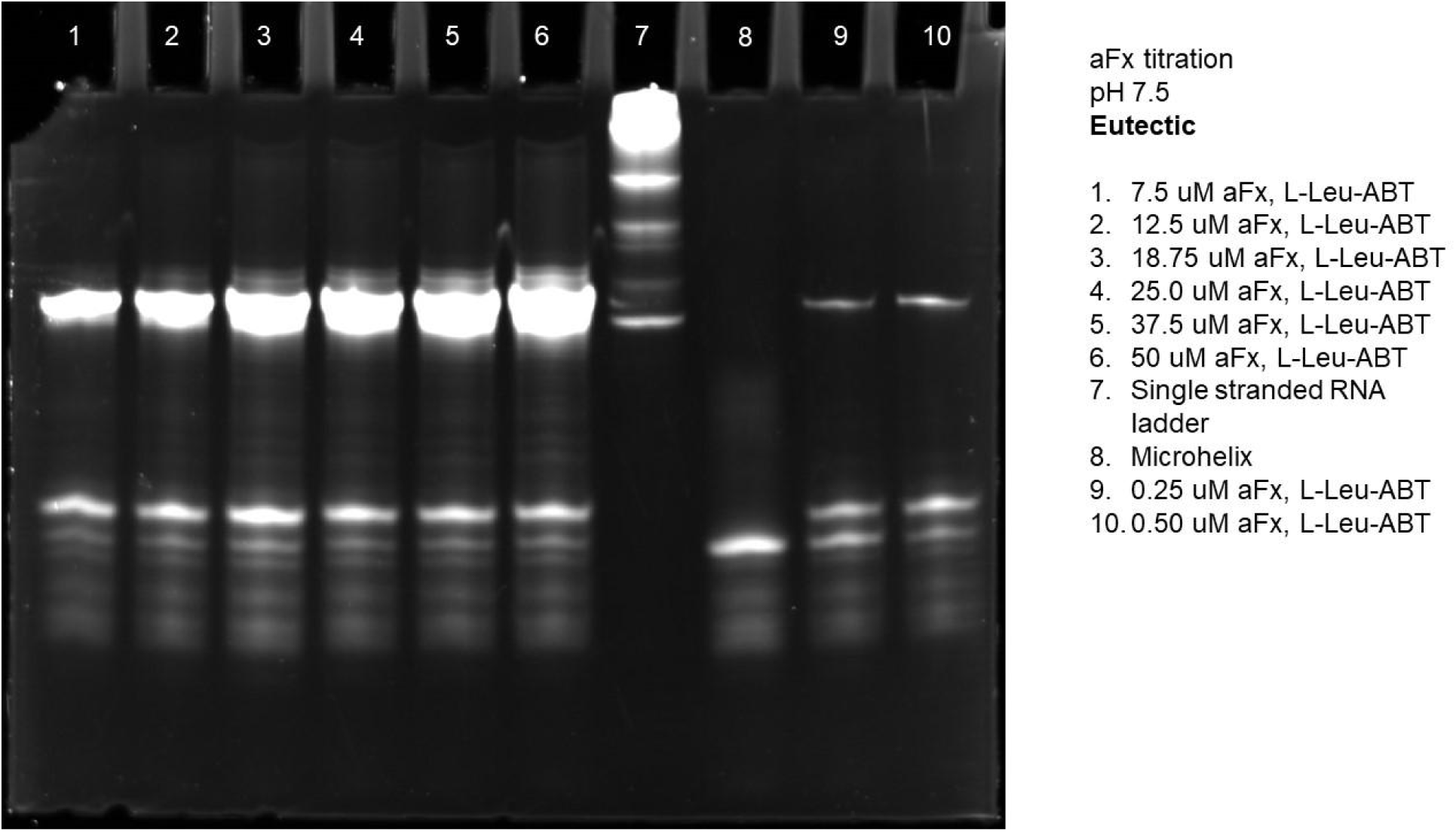
Gel of eutectic aFx titration at pH 7.5 using L-Leu-ABT.

**Figure S43.**
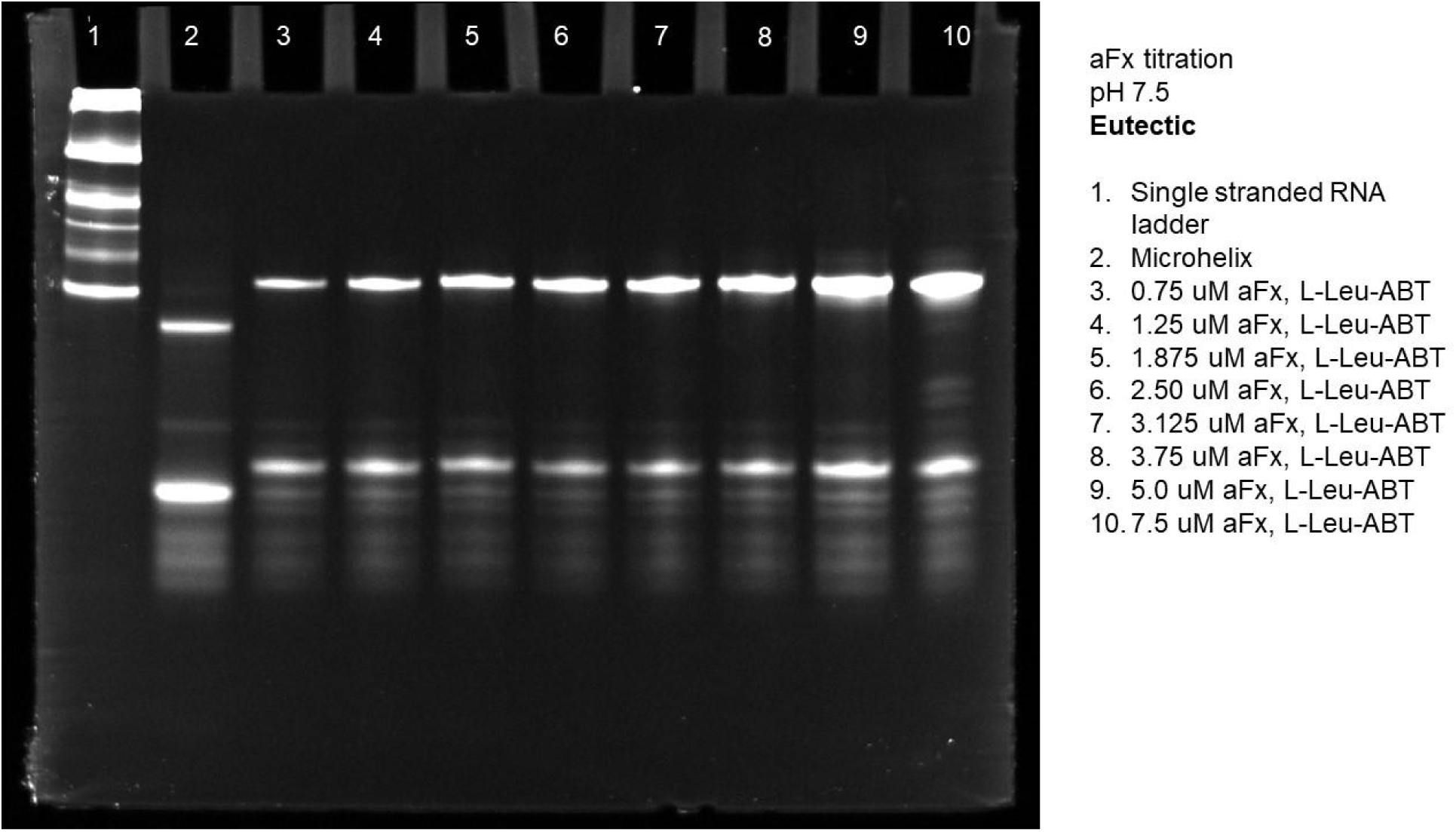
Gel of eutectic aFx titration at pH 7.5 using L-Leu-ABT.

**Figure S44.**
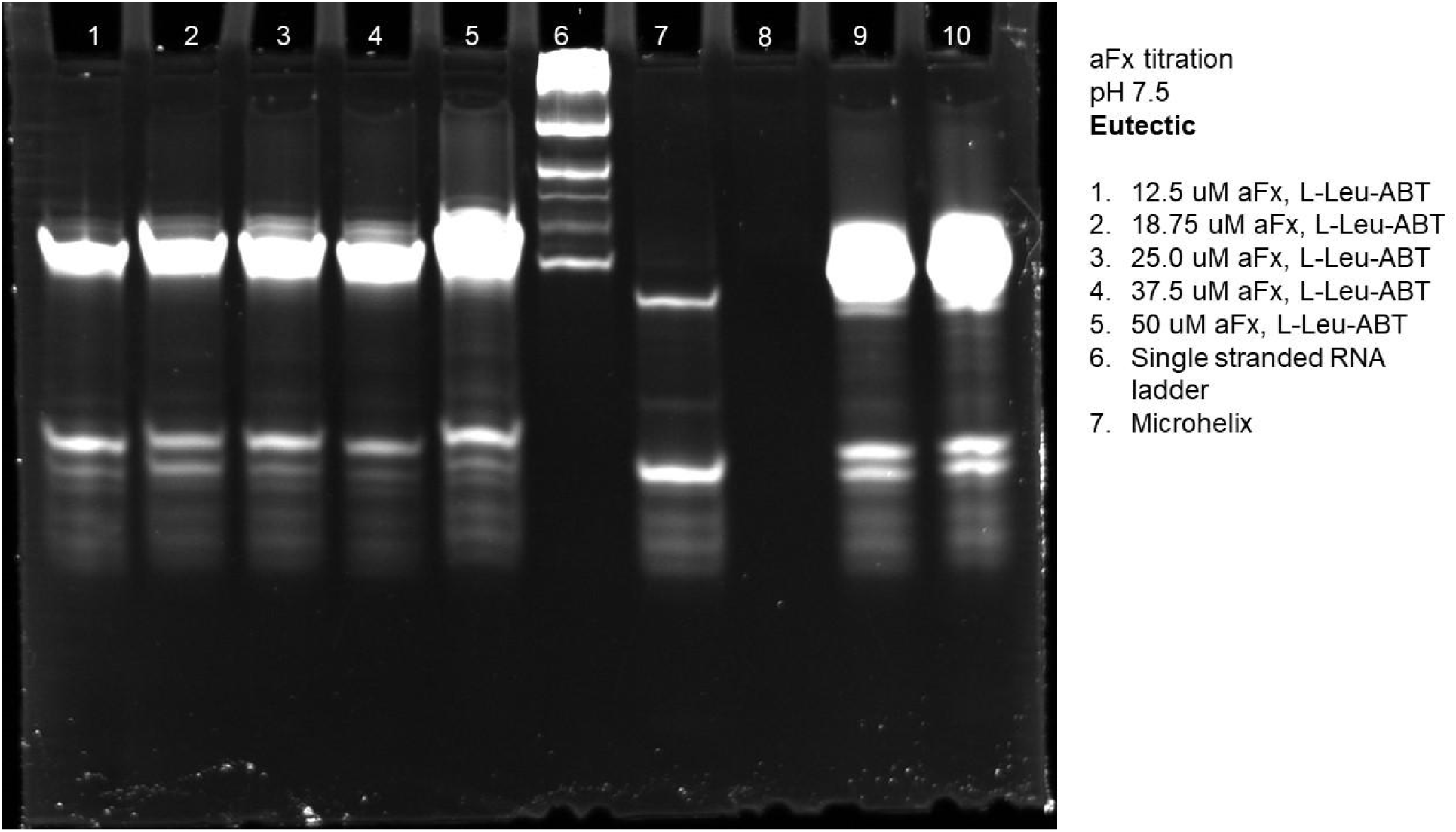
Gel of eutectic aFx titration at pH 7.5 using L-Leu-ABT.

**Figure S45.**
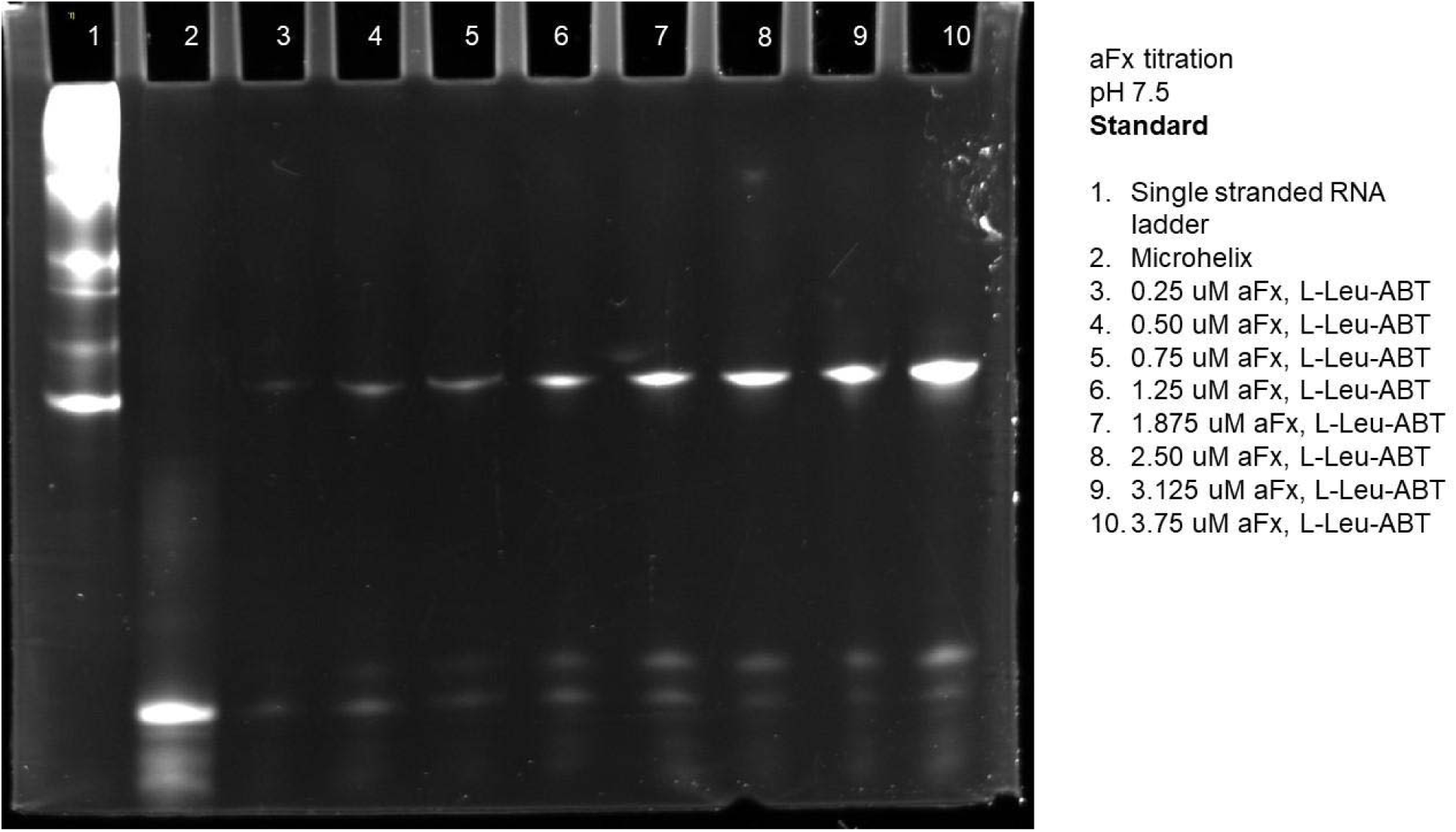
Gel of standard aFx titration at pH 7.5 using L-Leu-ABT.

**Figure S46.**
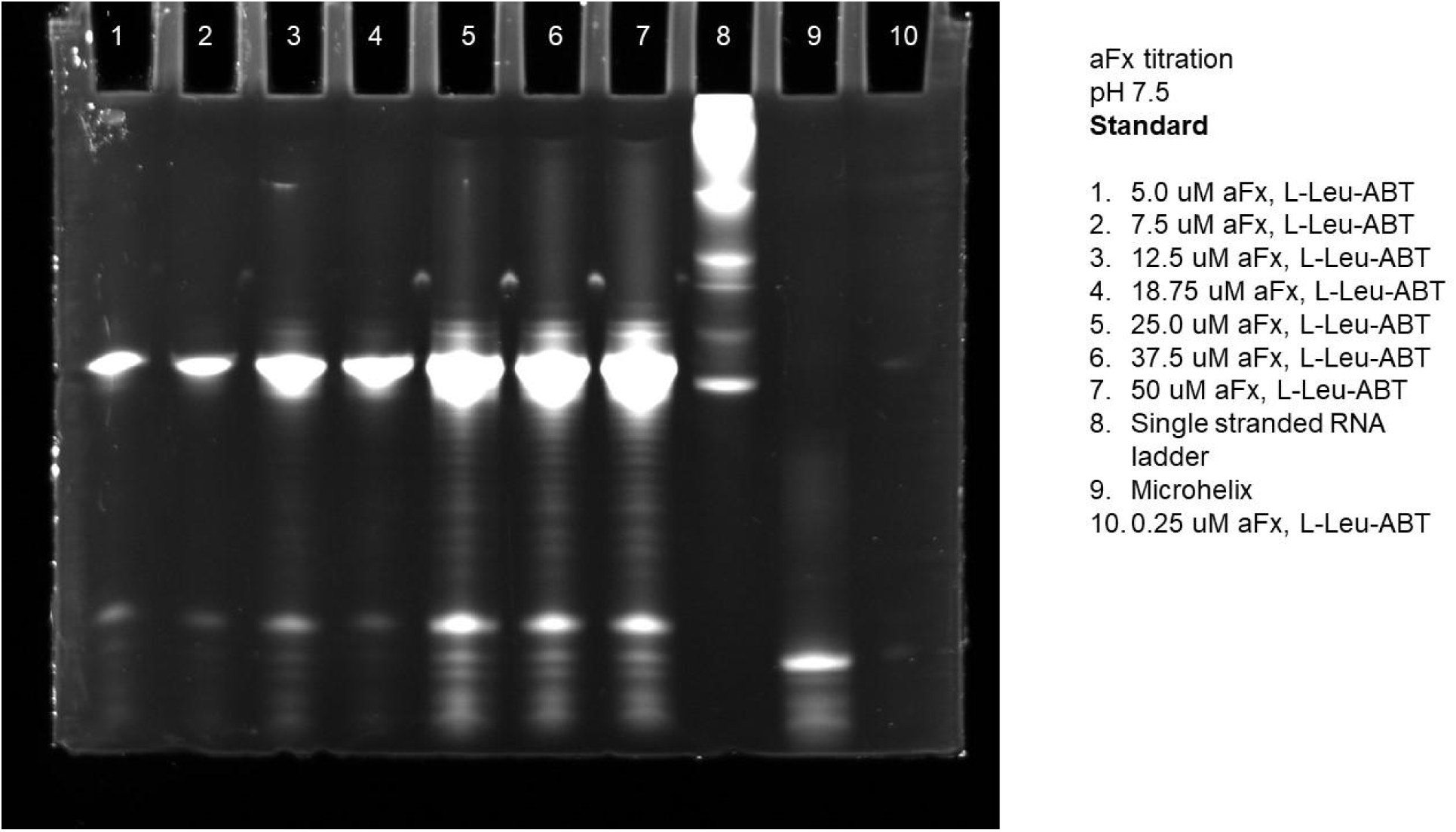
Gel of standard aFx titration at pH 7.5 using L-Leu-ABT.

**Figure S47.**
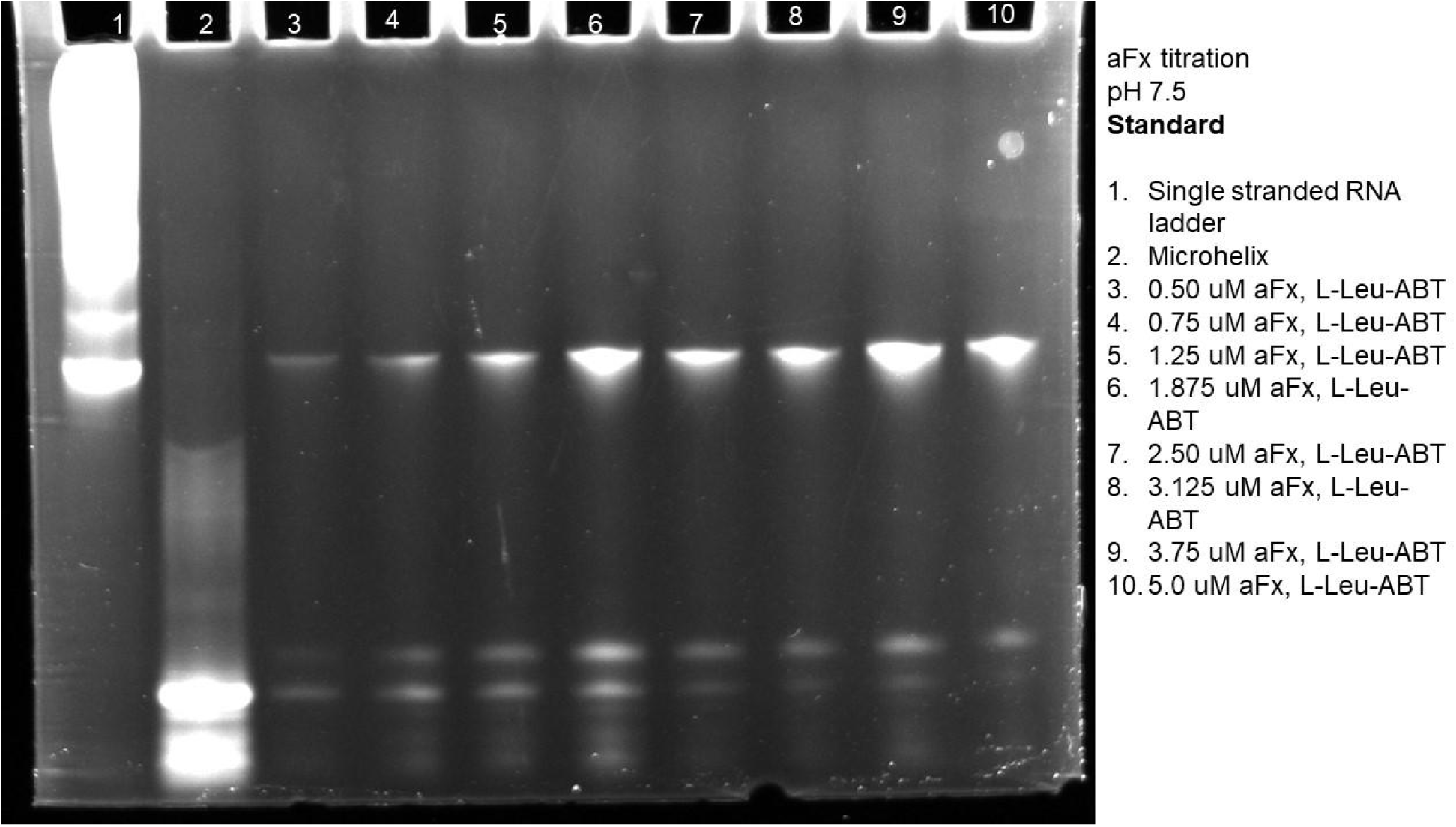
Gel of standard aFx titration at pH 7.5 using L-Leu-ABT.

**Figure S48.**
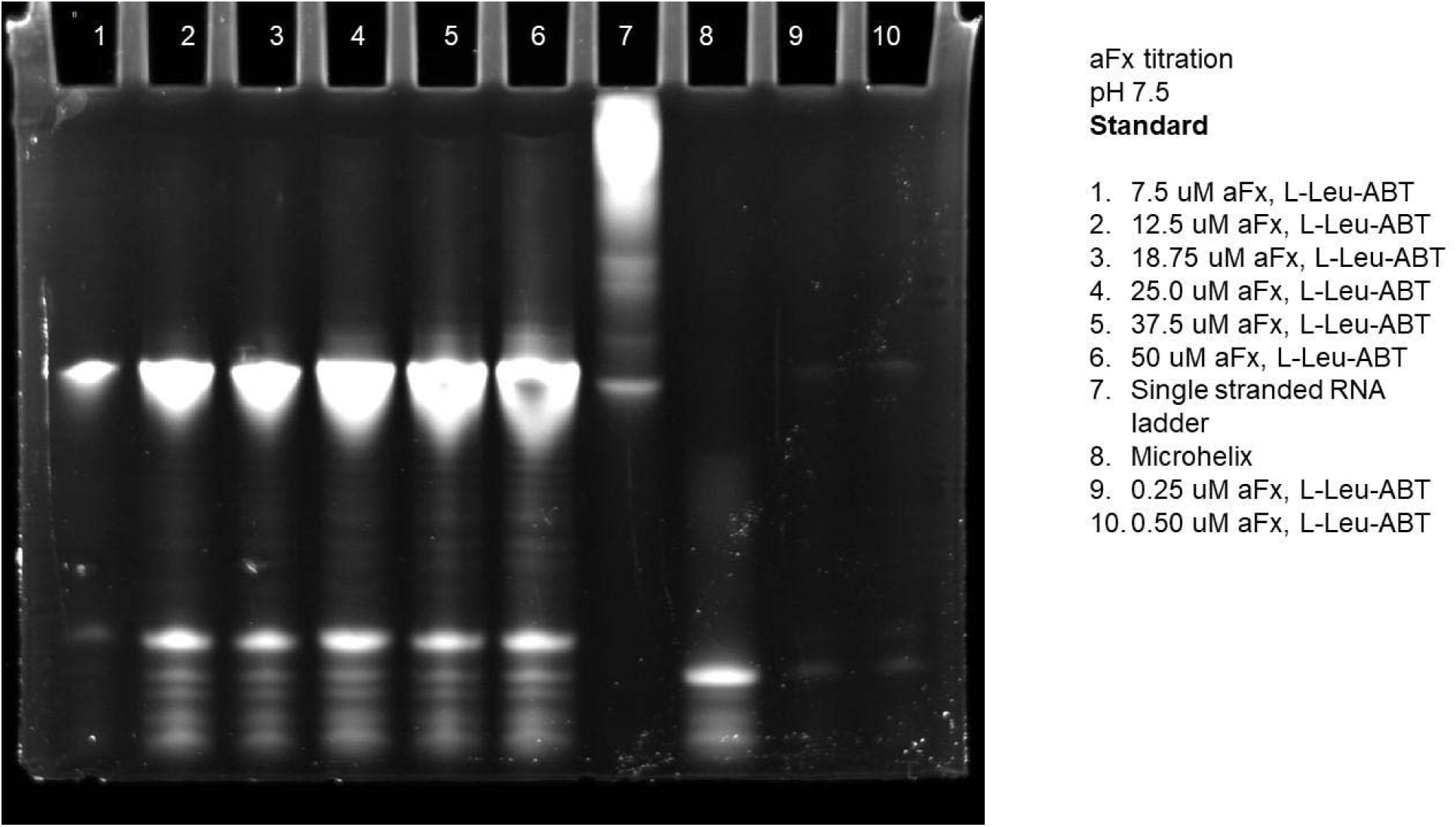
Gel of standard aFx titration at pH 7.5 using L-Leu-ABT.

**Figure S49.**
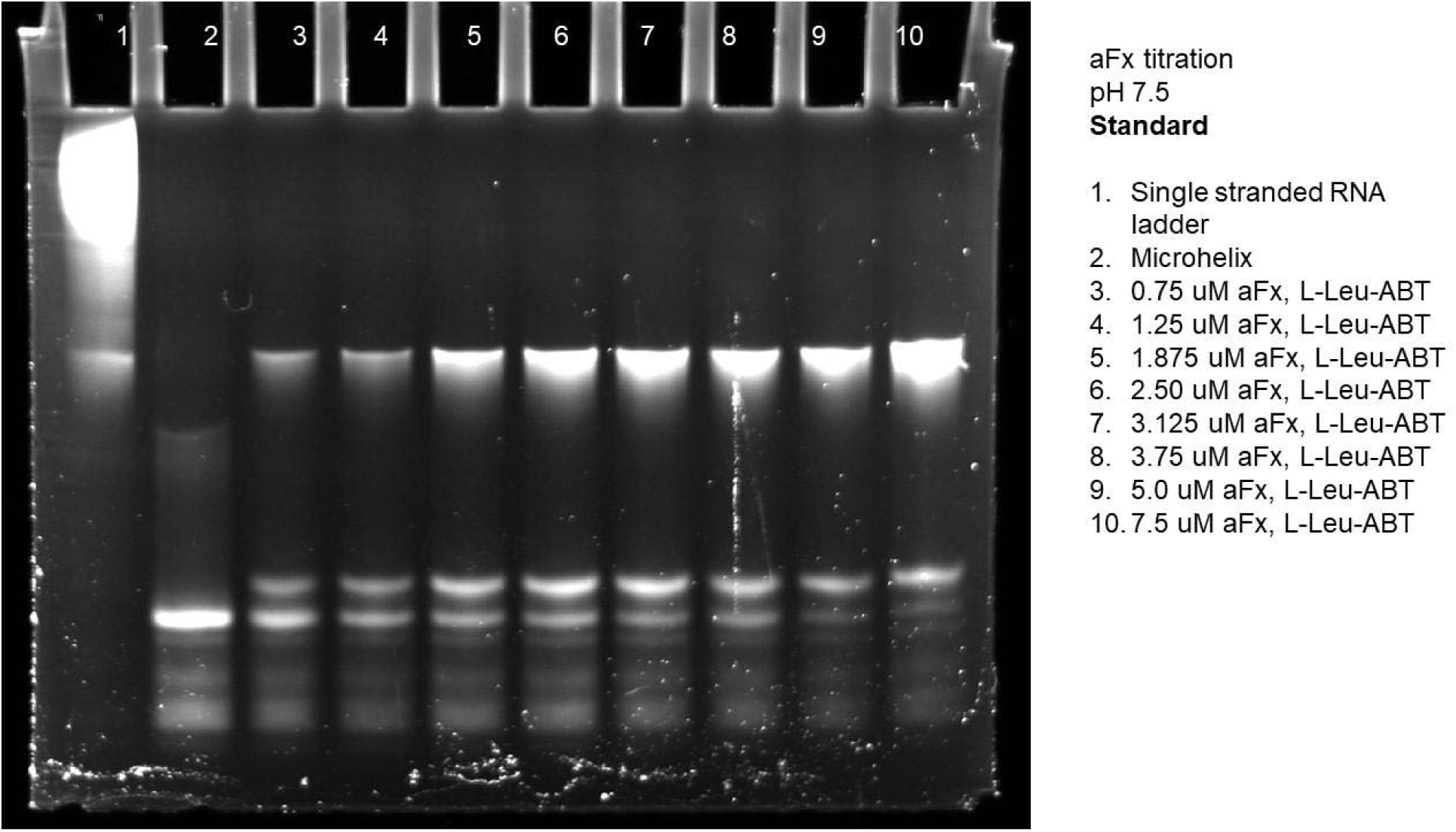
Gel of standard aFx titration at pH 7.5 using L-Leu-ABT.

**Figure S50.**
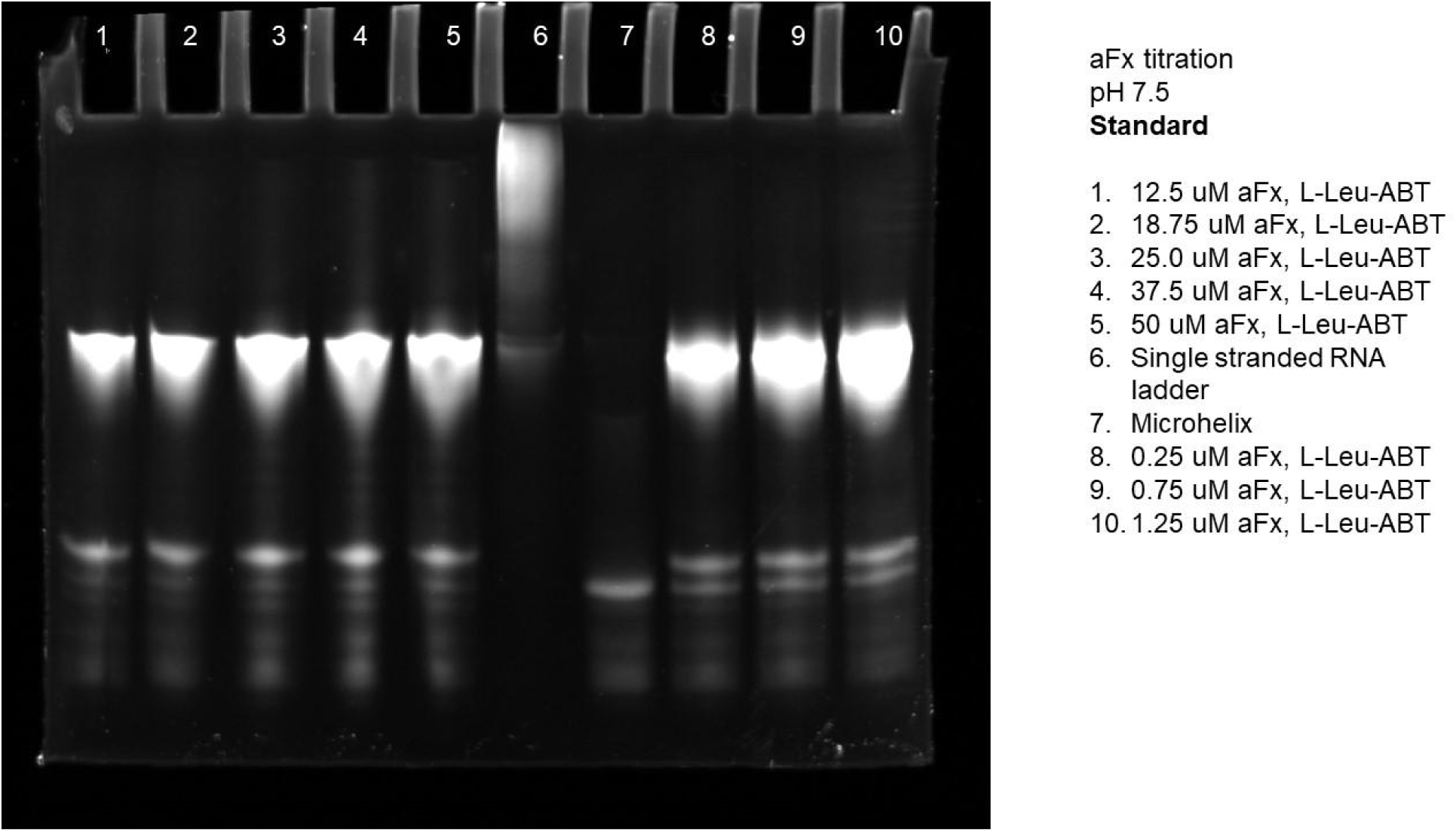
Gel of standard aFx titration at pH 7.5 using L-Leu-ABT.

**Figure S51.**
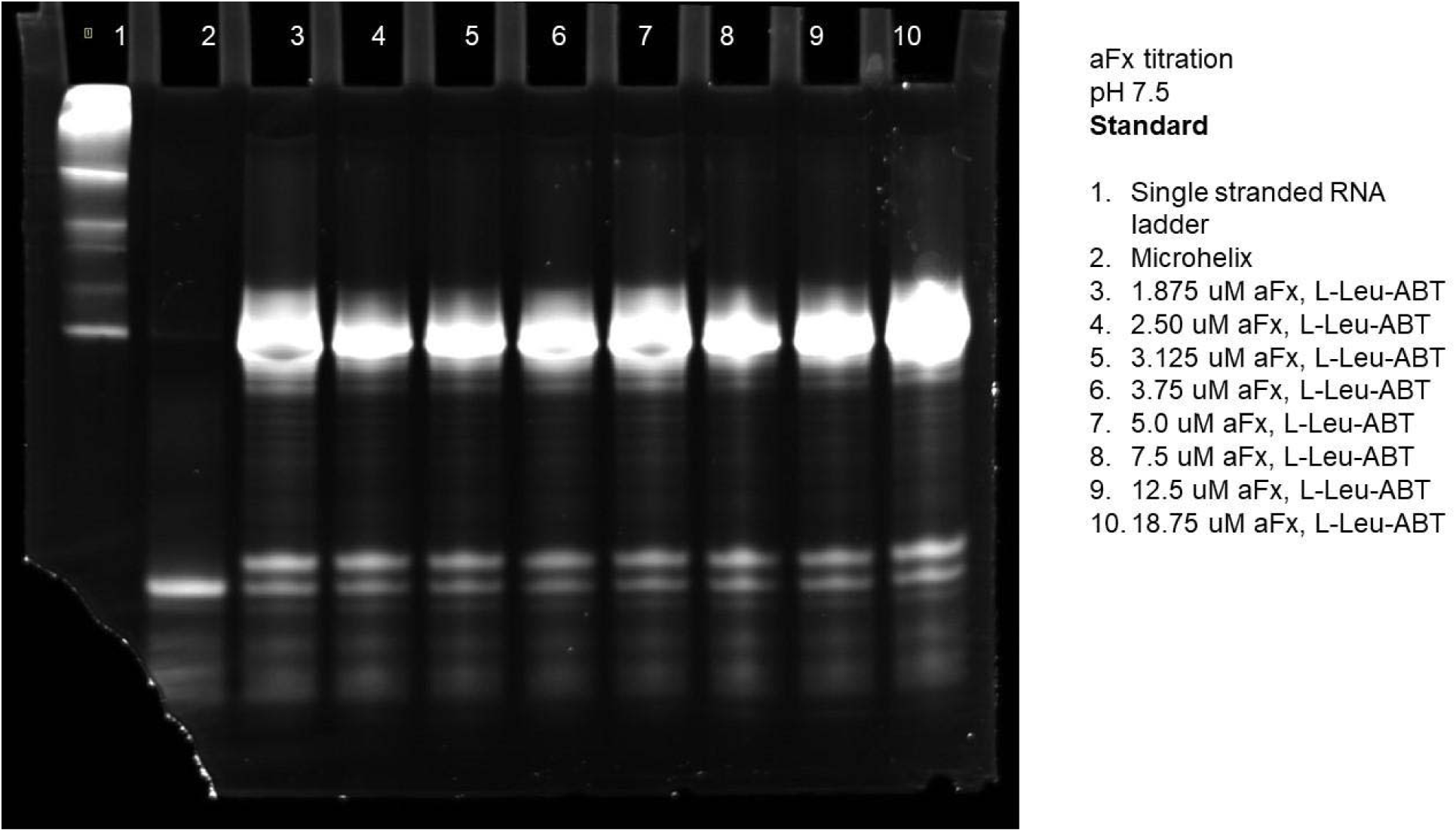
Gel of standard aFx titration at pH 7.5 using L-Leu-ABT.

**Figure S52.**
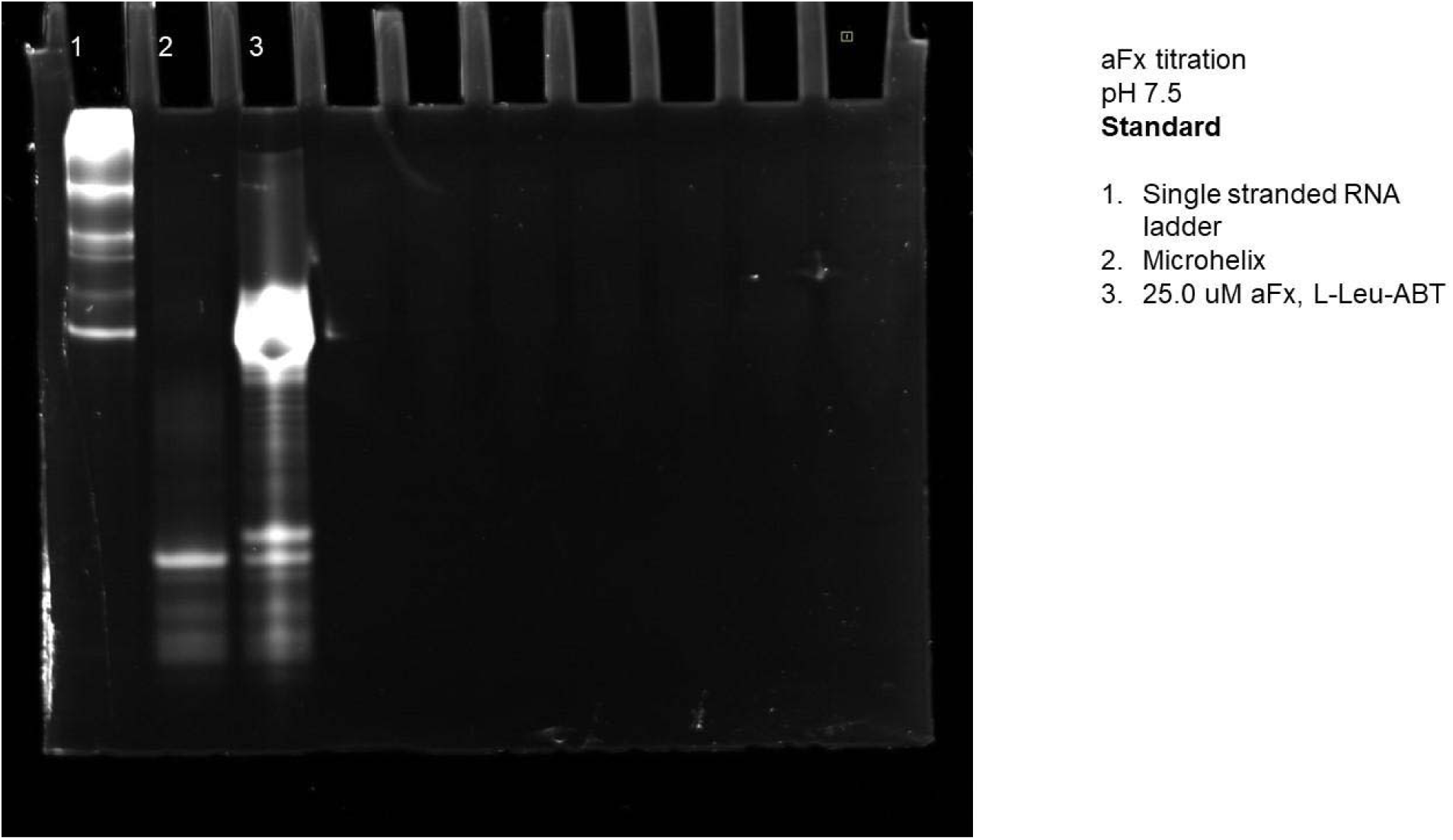
Gel of standard aFx titration at pH 7.5 using L-Leu-ABT.

**Figure S53.**
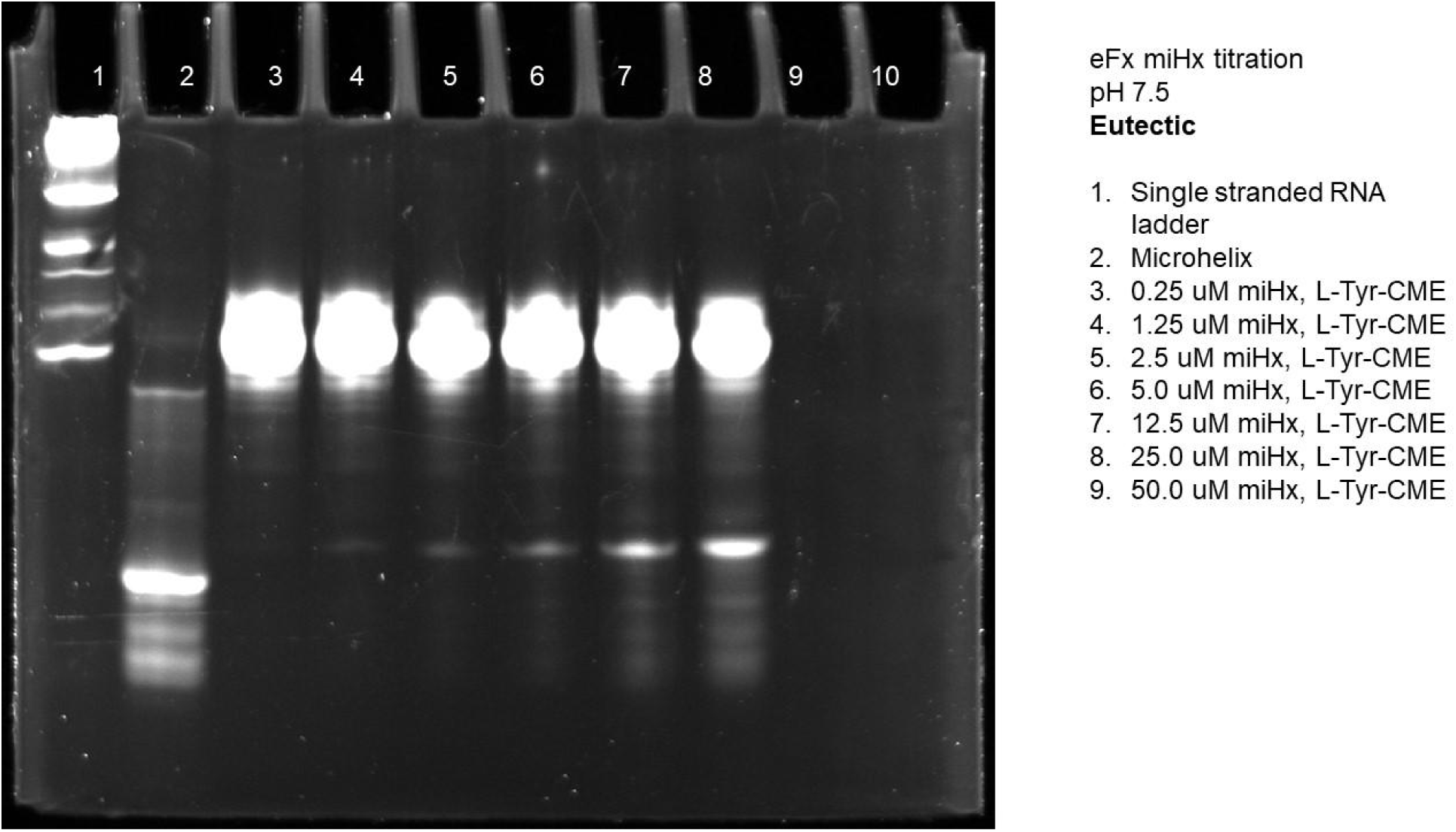
Gel of eutectic microhelix titration with eFx at pH 7.5 using L-Tyr-CME.

**Figure S54.**
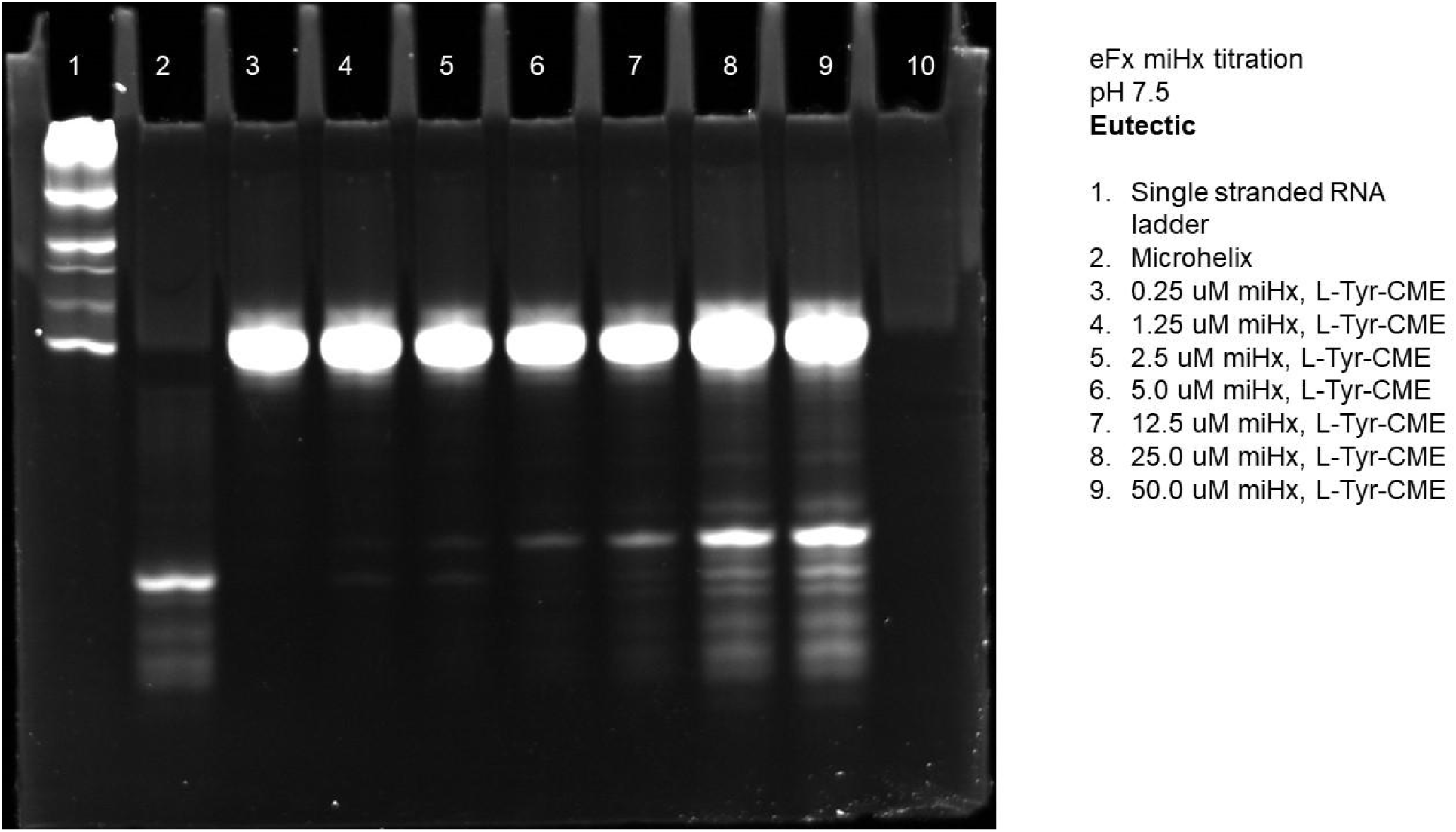
Gel of eutectic microhelix titration with eFx at pH 7.5 using L-Tyr-CME.

**Figure S55.**
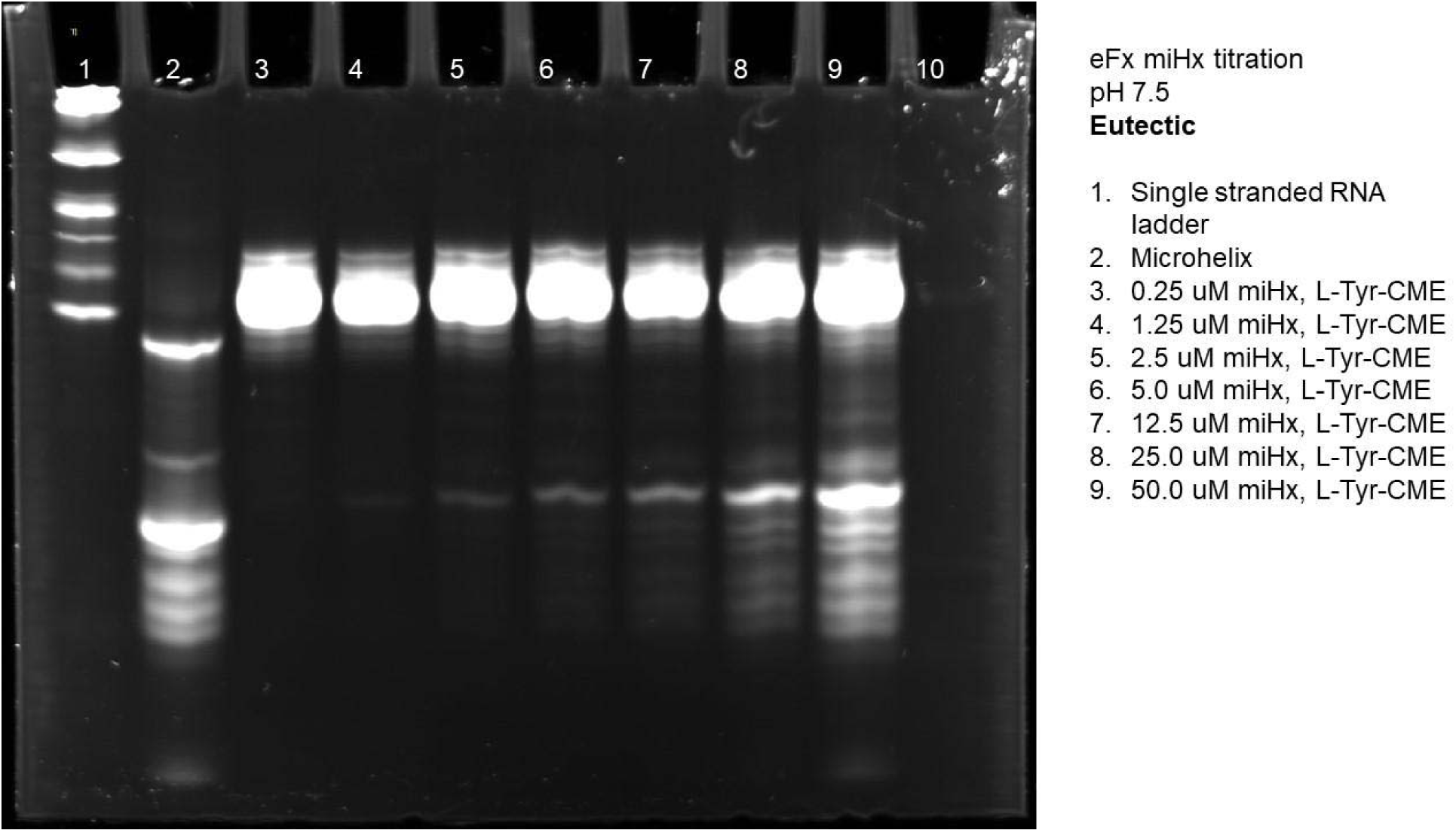
Gel of eutectic microhelix titration with eFx at pH 7.5 using L-Tyr-CME.

**Figure S56.**
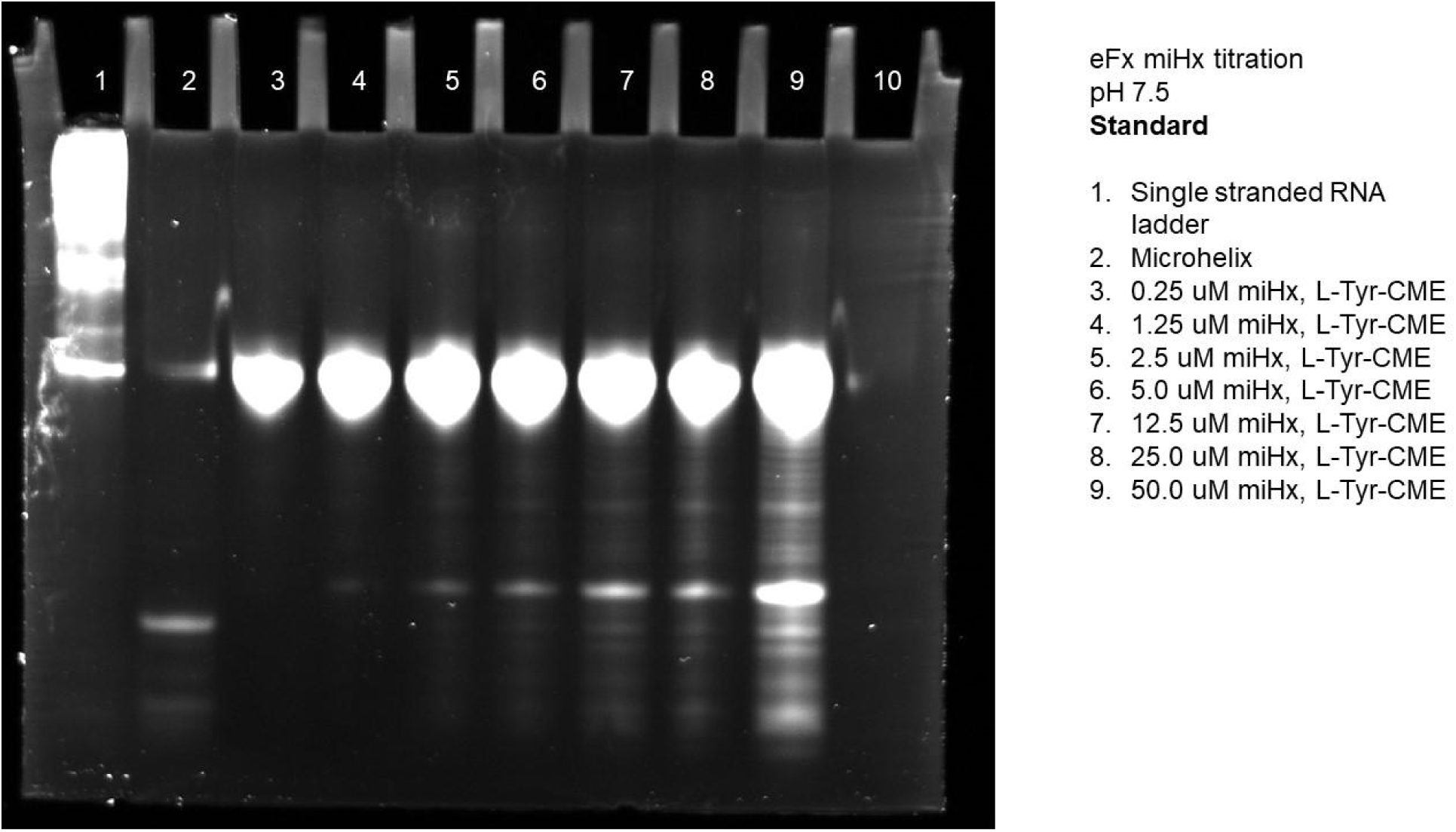
Gel of standard microhelix titration with eFx at pH 7.5 using L-Tyr-CME.

**Figure S57.**
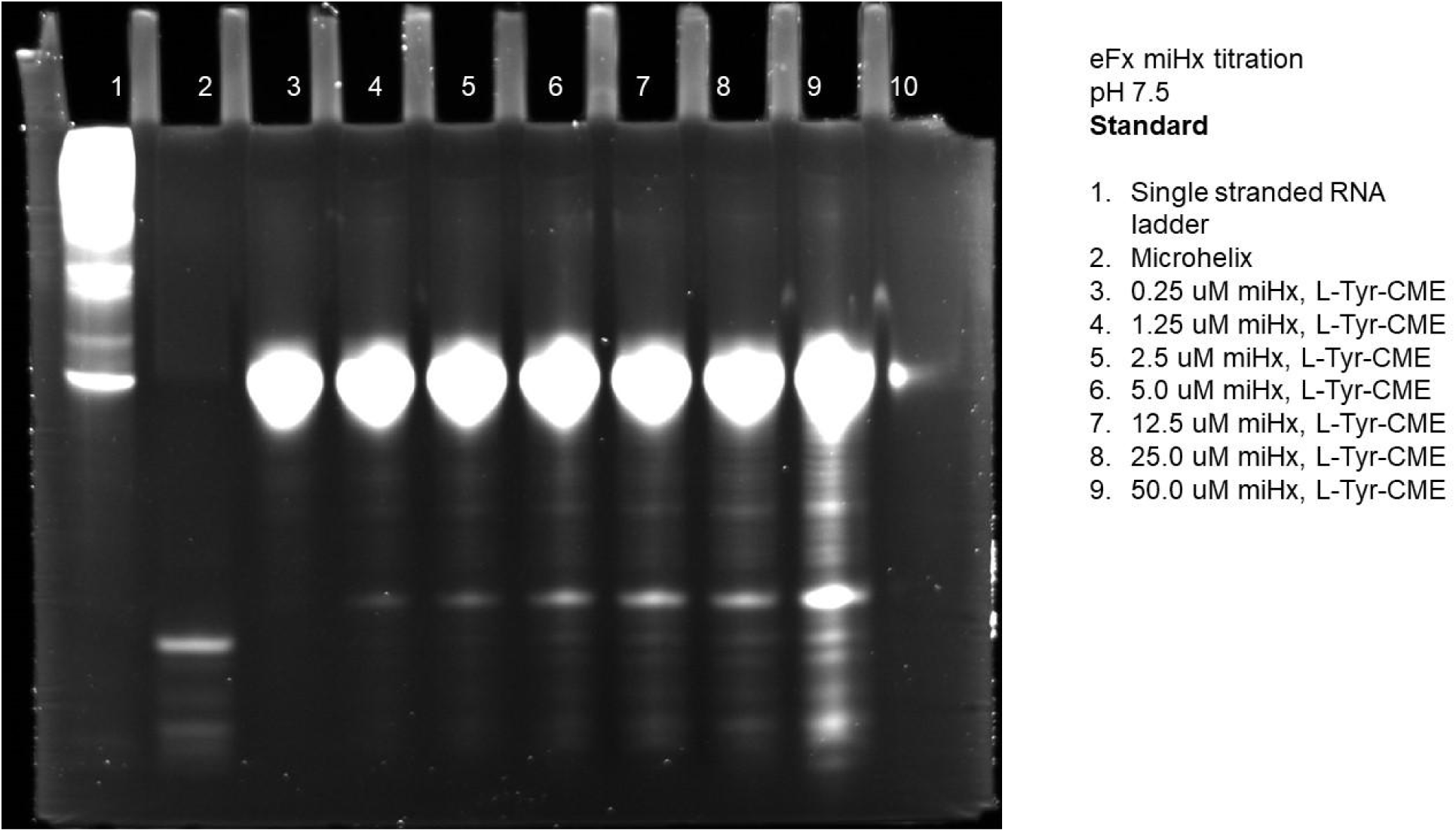
Gel of standard microhelix titration with eFx at pH 7.5 using L-Tyr-CME.

**Figure S58.**
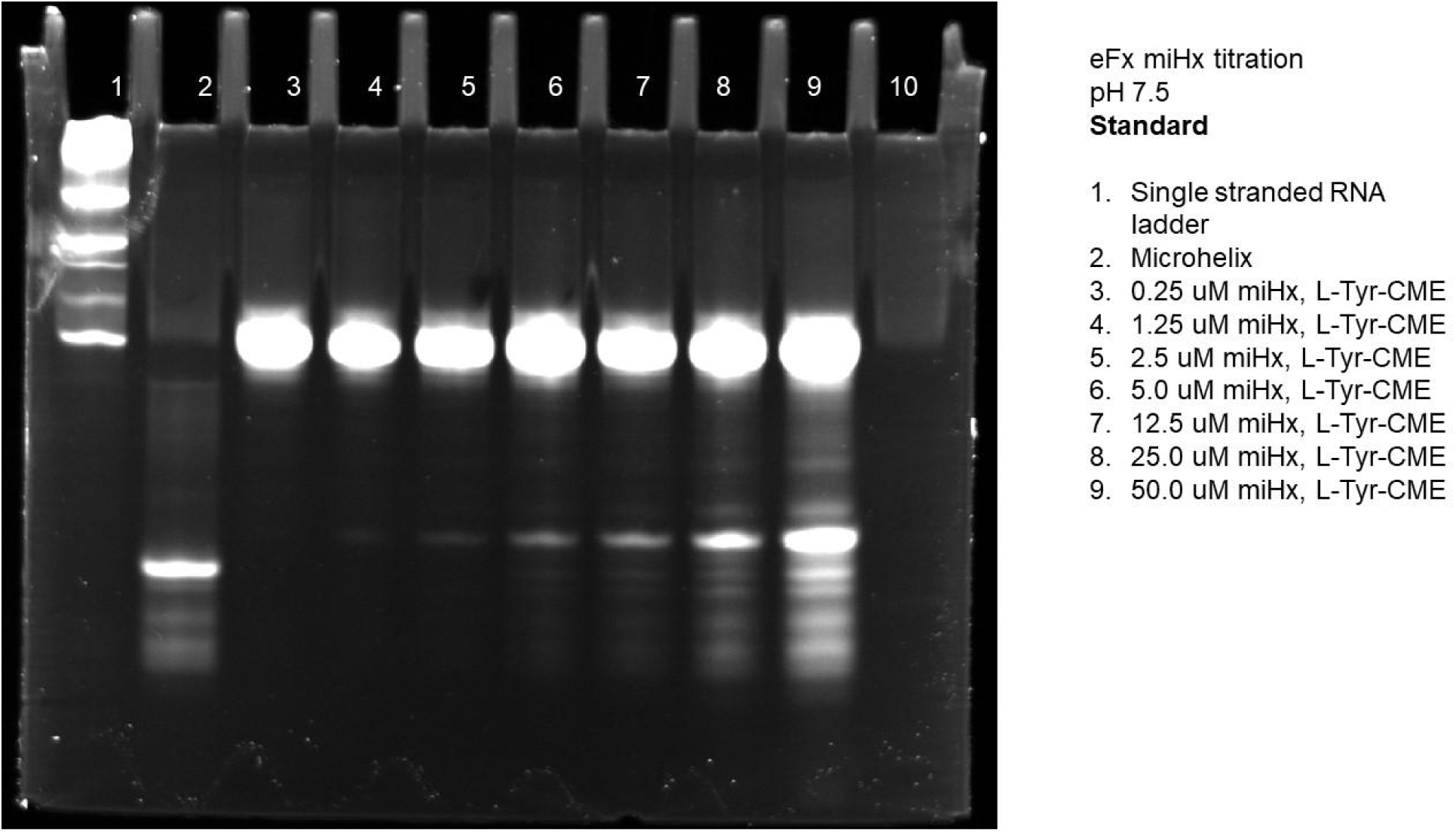
Gel of standard microhelix titration with eFx at pH 7.5 using L-Tyr-CME.

**Figure S59.**
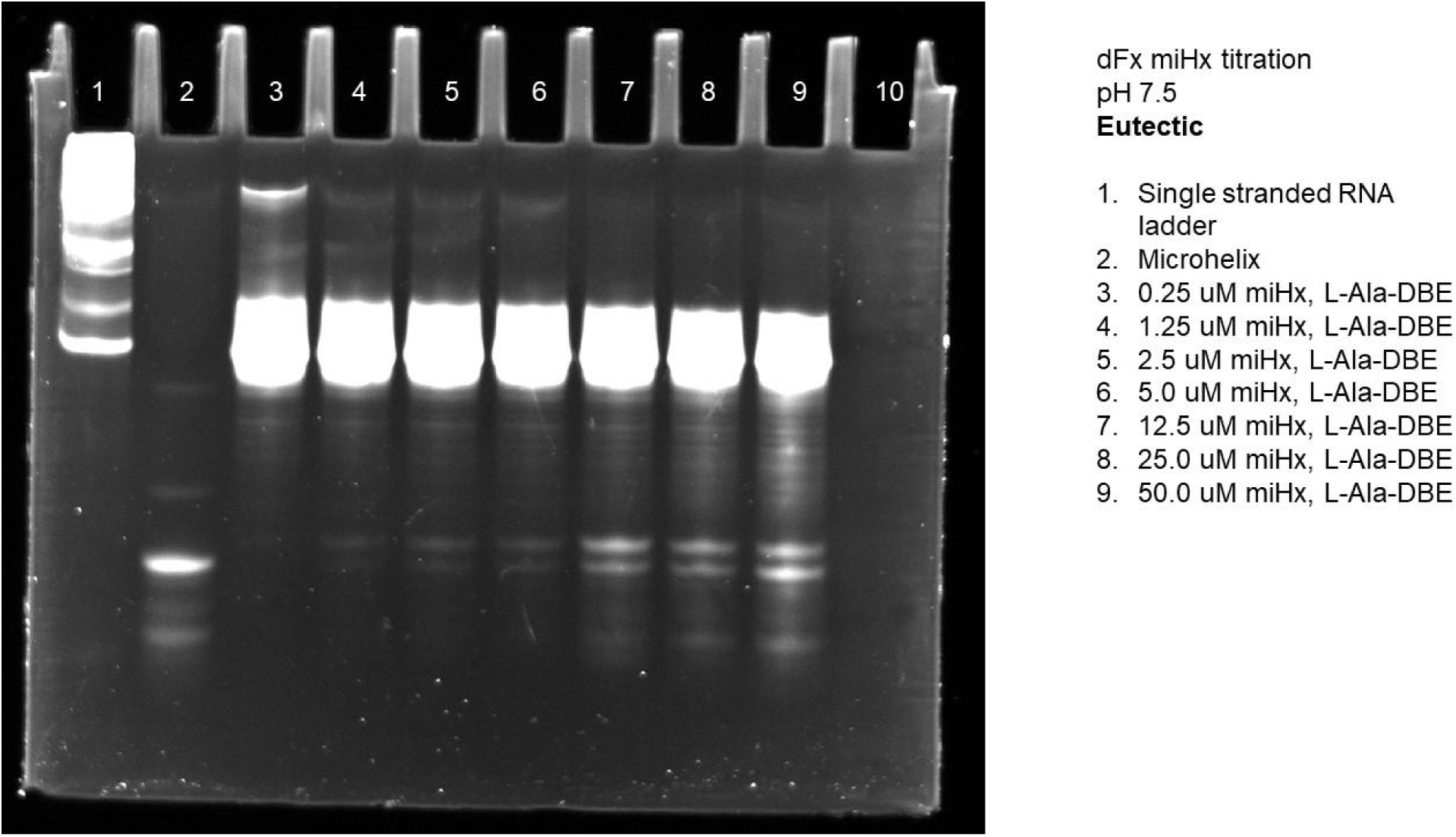
Gel of eutectic microhelix titration with dFx at pH 7.5 using L-Ala-DBE.

**Figure S60.**
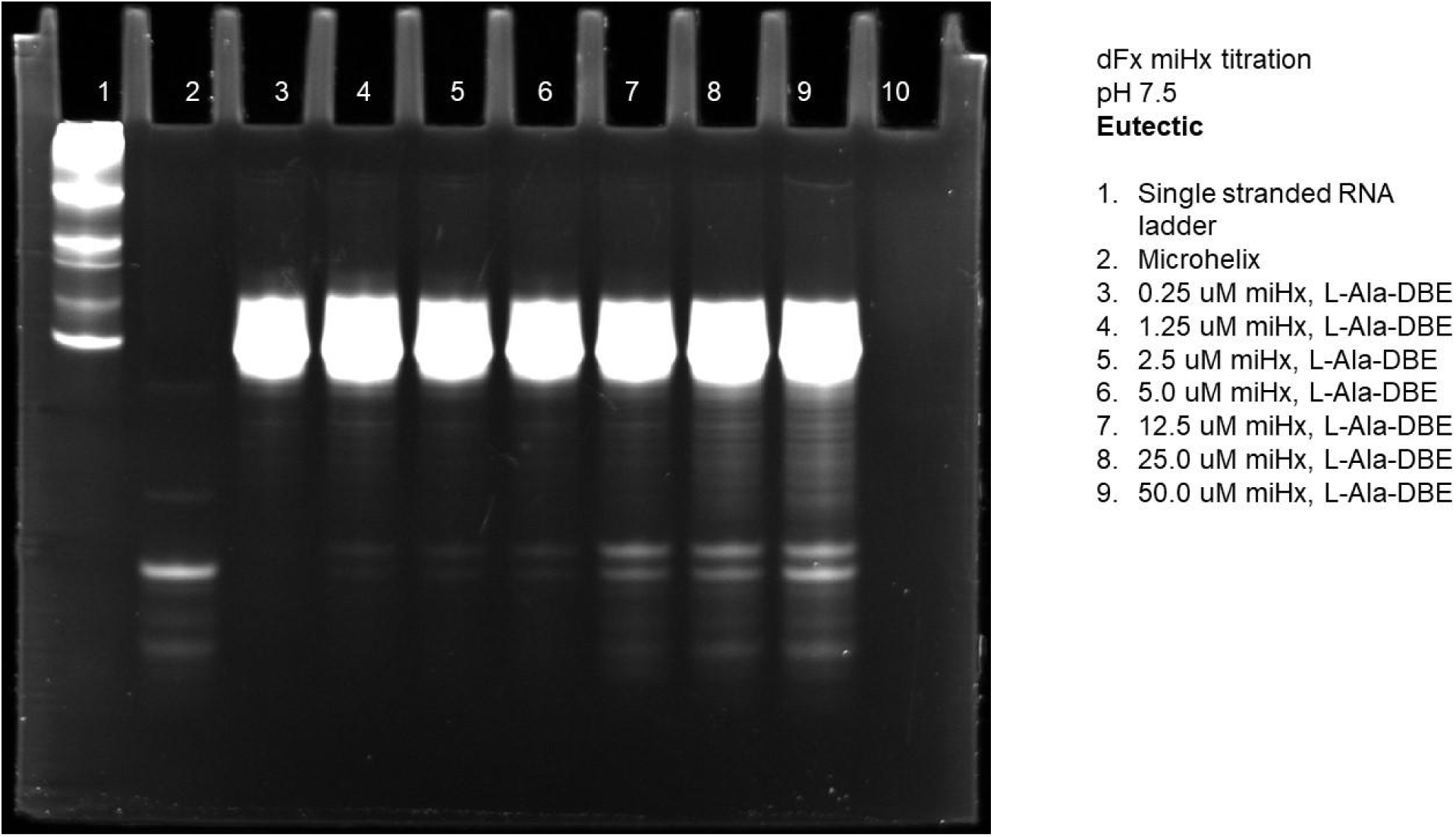
Gel of eutectic microhelix titration with dFx at pH 7.5 using L-Ala-DBE.

**Figure S61.**
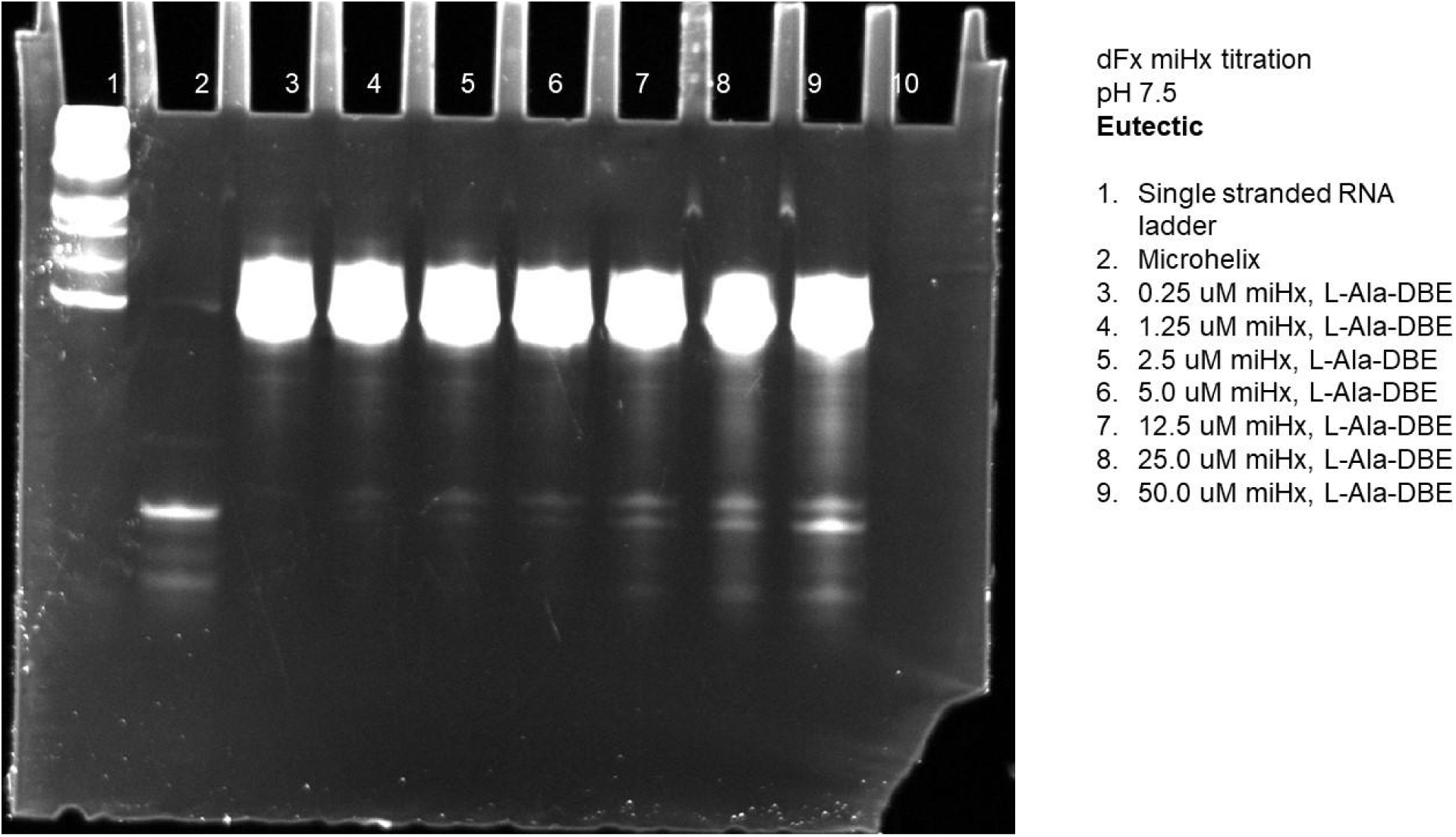
Gel of eutectic microhelix titration with dFx at pH 7.5 using L-Ala-DBE.

**Figure S62.**
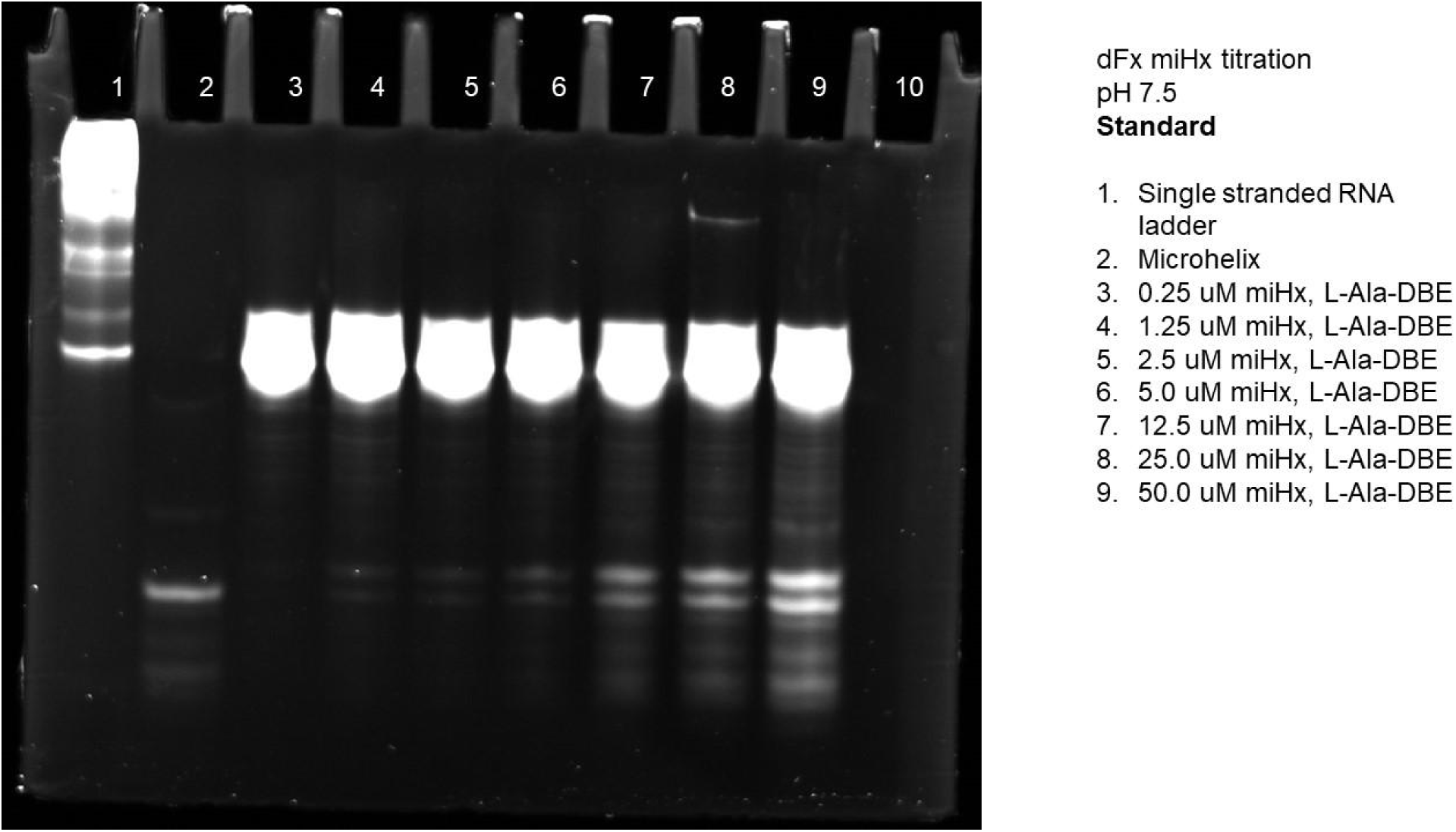
Gel of standard microhelix titration with dFx at pH 7.5 using L-Ala-DBE.

**Figure S63.**
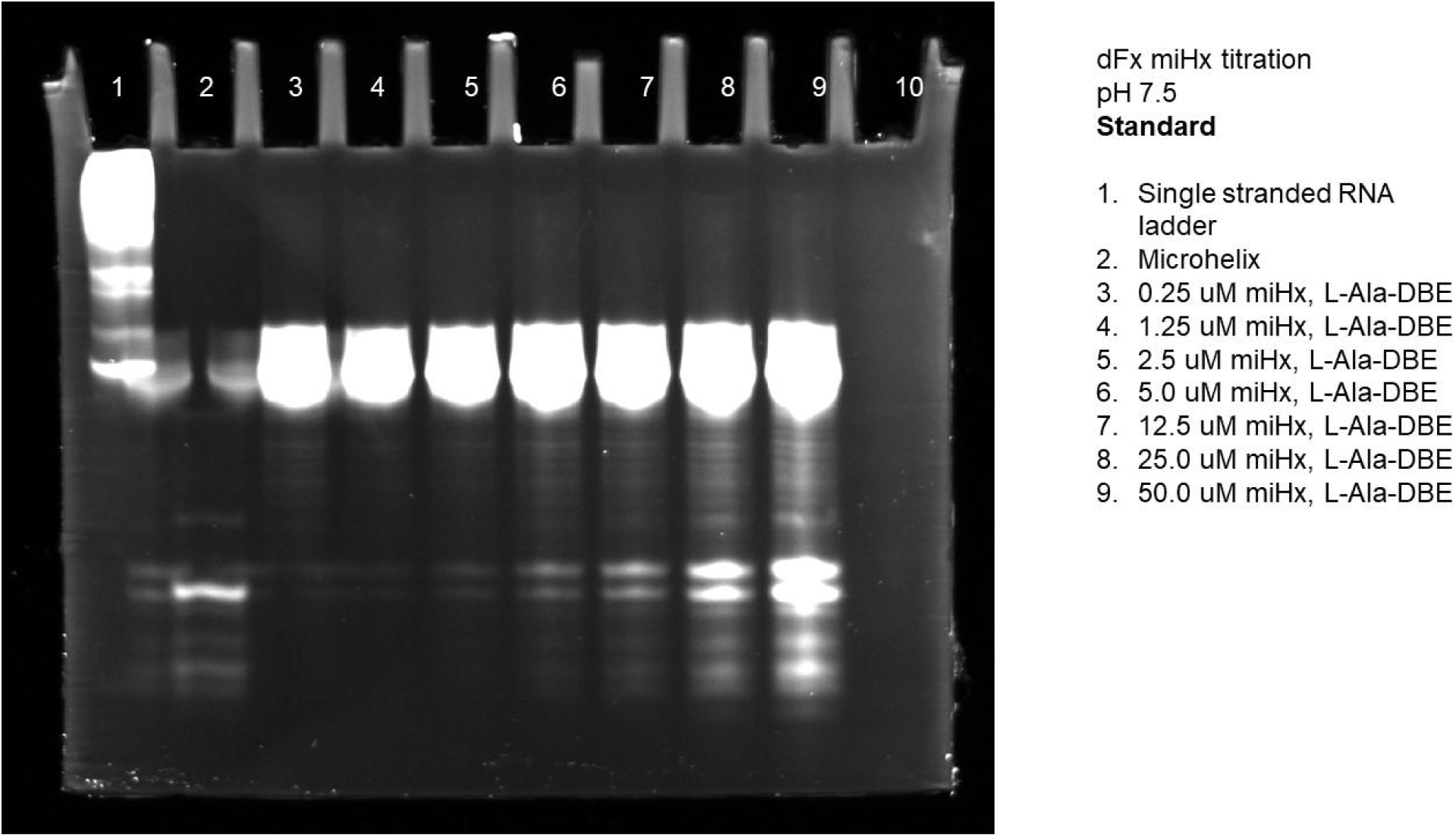
Gel of standard microhelix titration with dFx at pH 7.5 using L-Ala-DBE.

**Figure S64.**
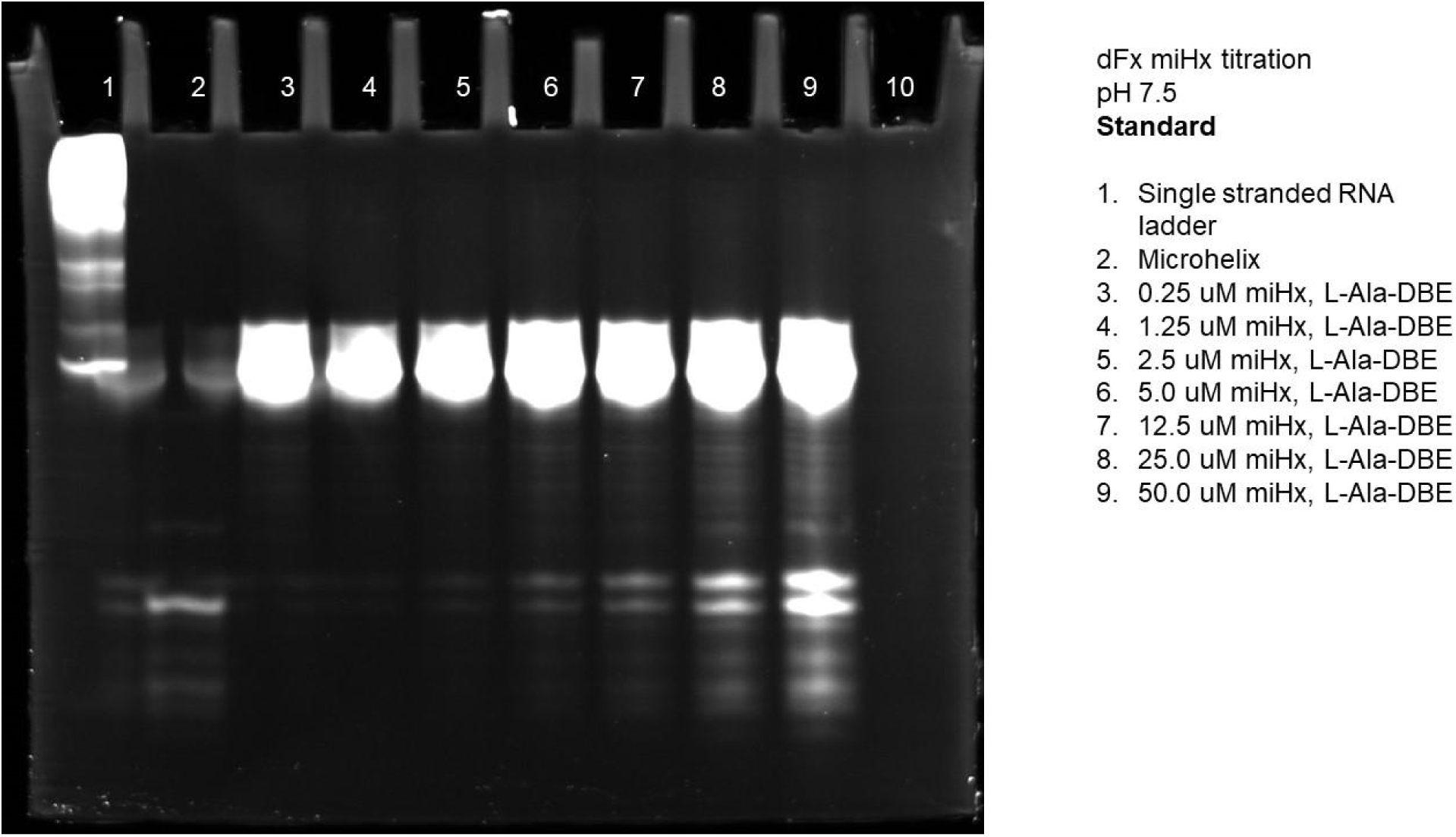
Gel of standard microhelix titration with dFx at pH 7.5 using L-Ala-DBE.

**Figure S65.**
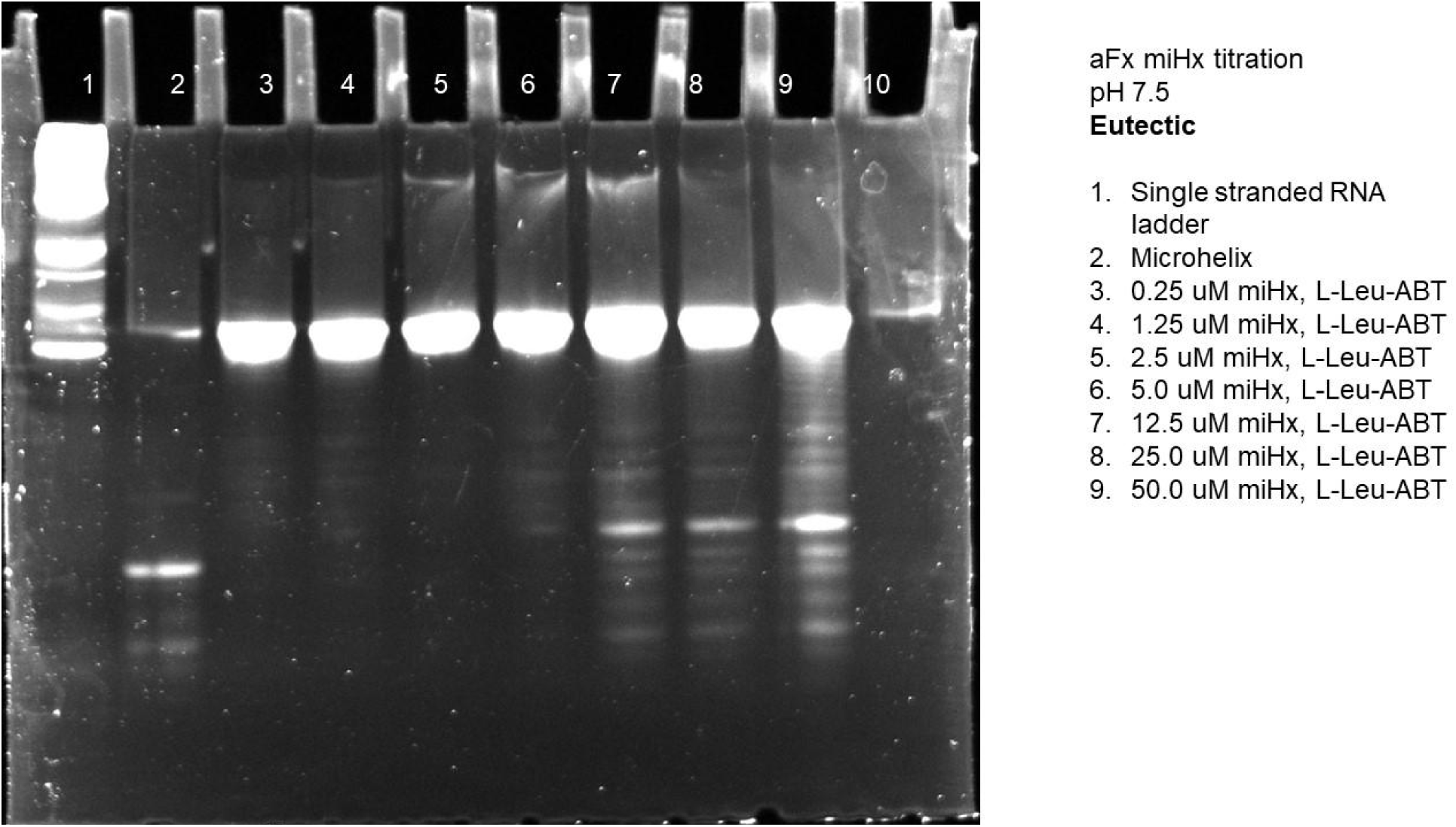
Gel of eutectic microhelix titration with aFx at pH 7.5 using L-Leu-ABT.

**Figure S66.**
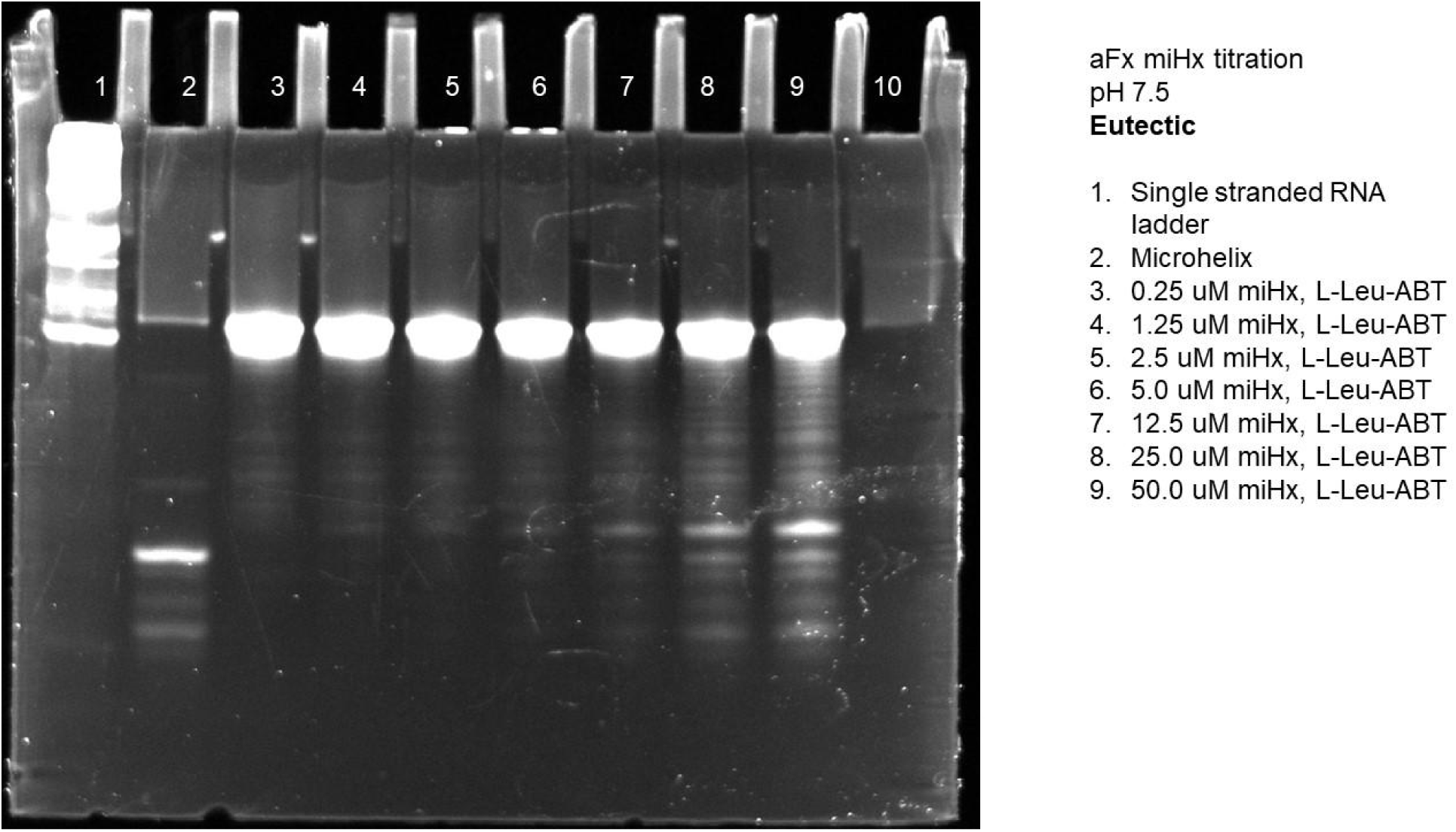
Gel of eutectic microhelix titration with aFx at pH 7.5 using L-Leu-ABT.

**Figure S67.**
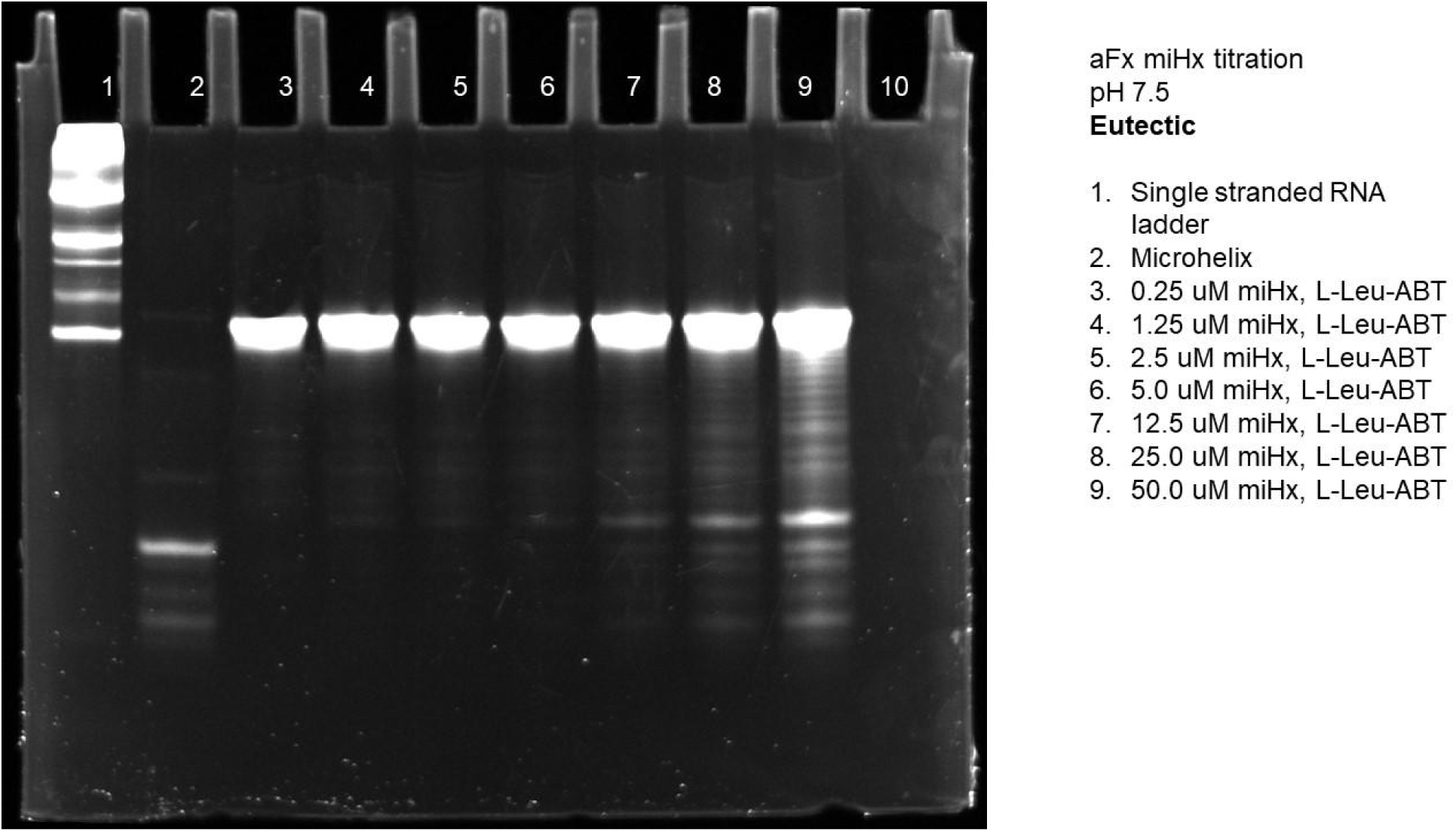
Gel of eutectic microhelix titration with aFx at pH 7.5 using L-Leu-ABT.

**Figure S68.**
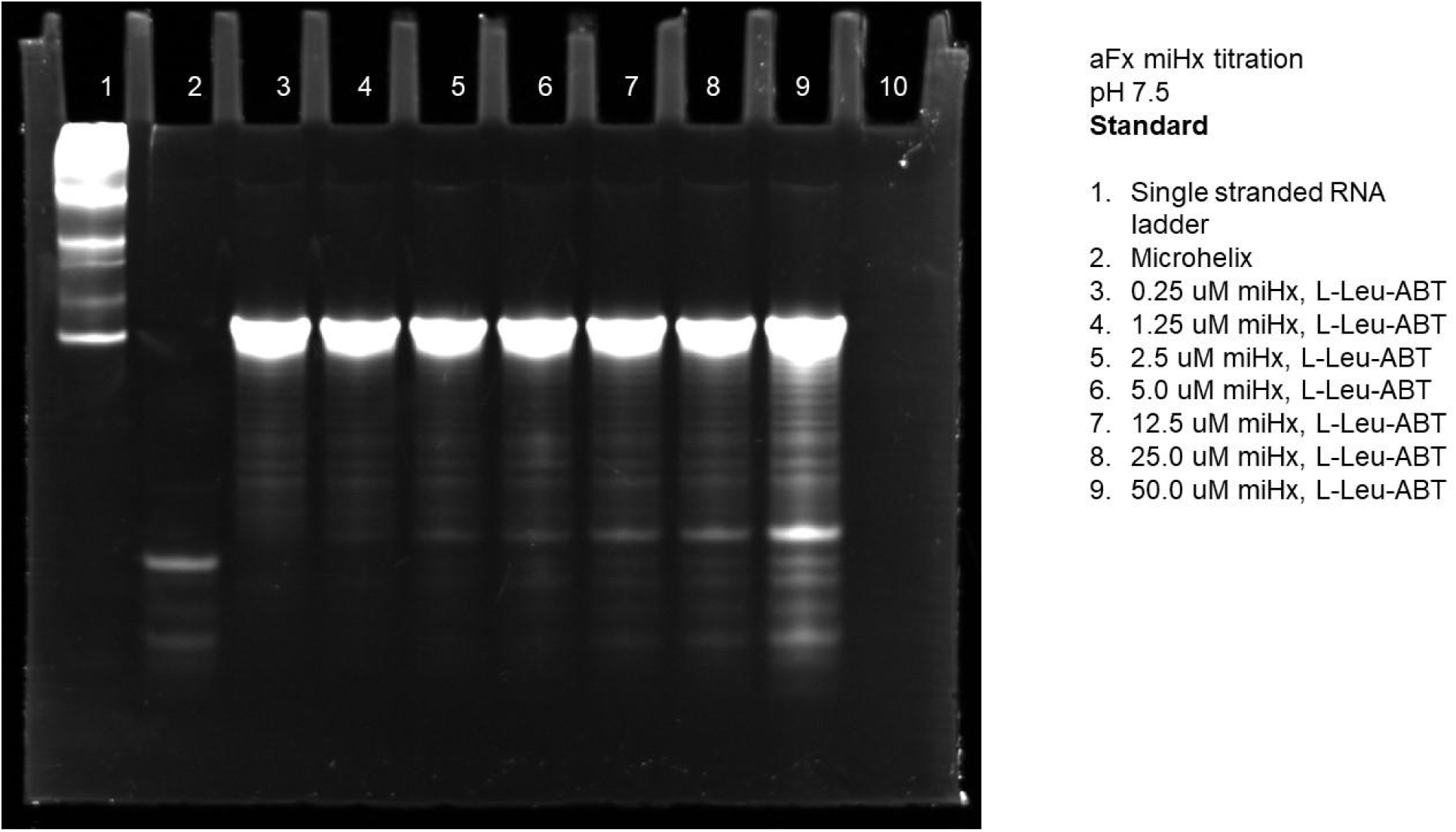
Gel of standard microhelix titration with aFx at pH 7.5 using L-Leu-ABT.

**Figure S69.**
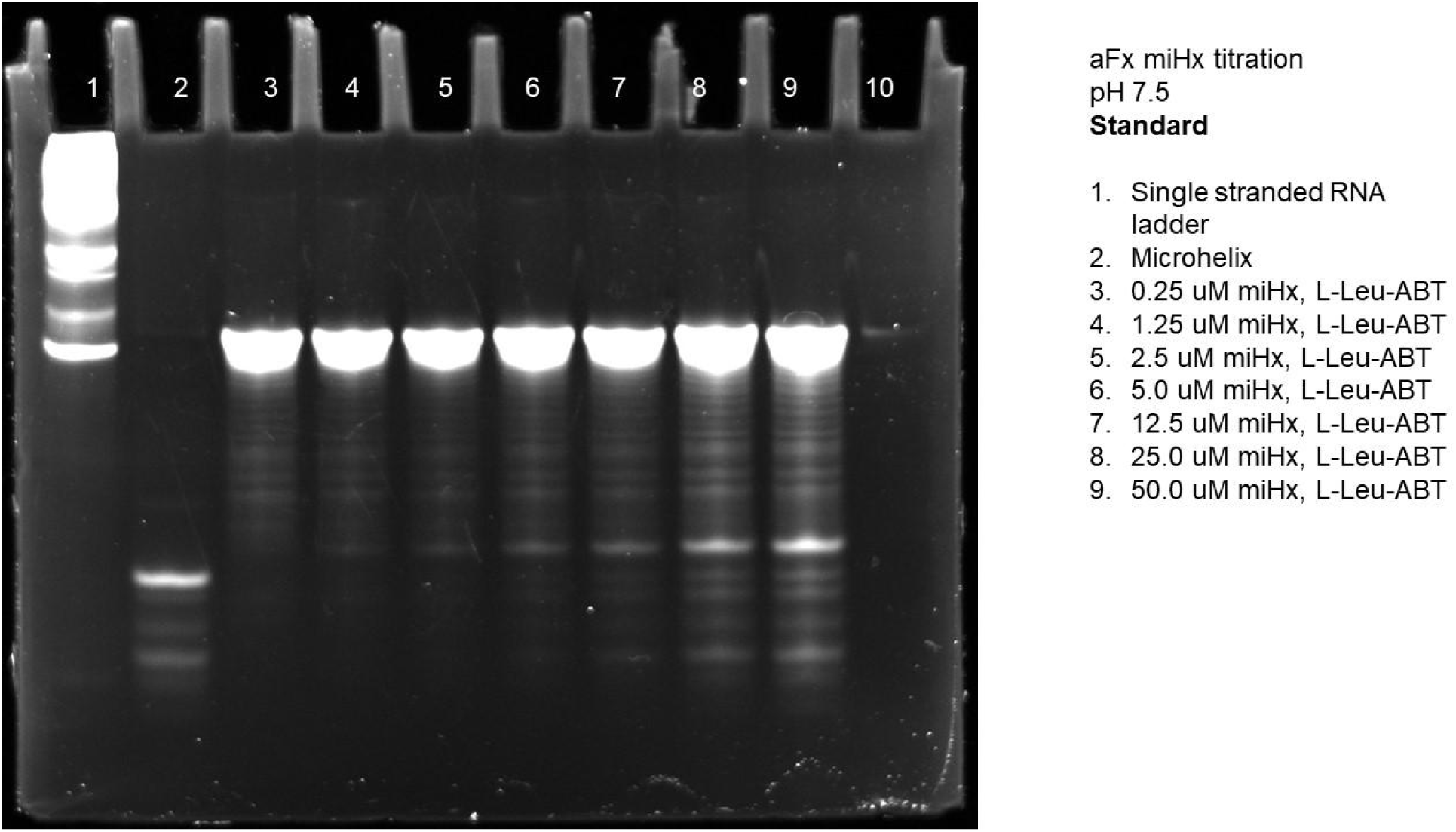
Gel of standard microhelix titration with aFx at pH 7.5 using L-Leu-ABT.

**Figure S70.**
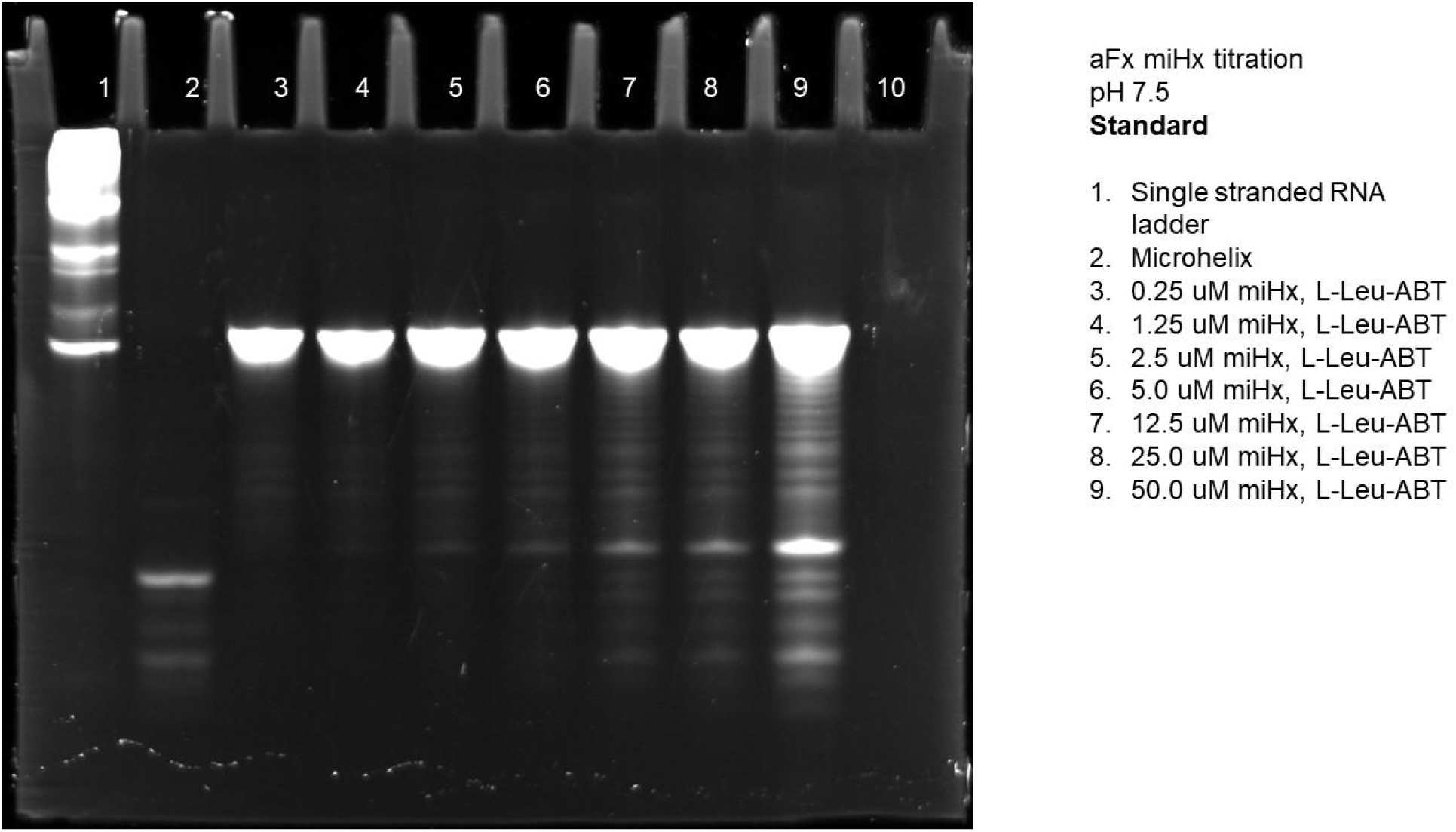
Gel of standard microhelix titration with aFx at pH 7.5 using L-Leu-ABT.

**Figure S72.**
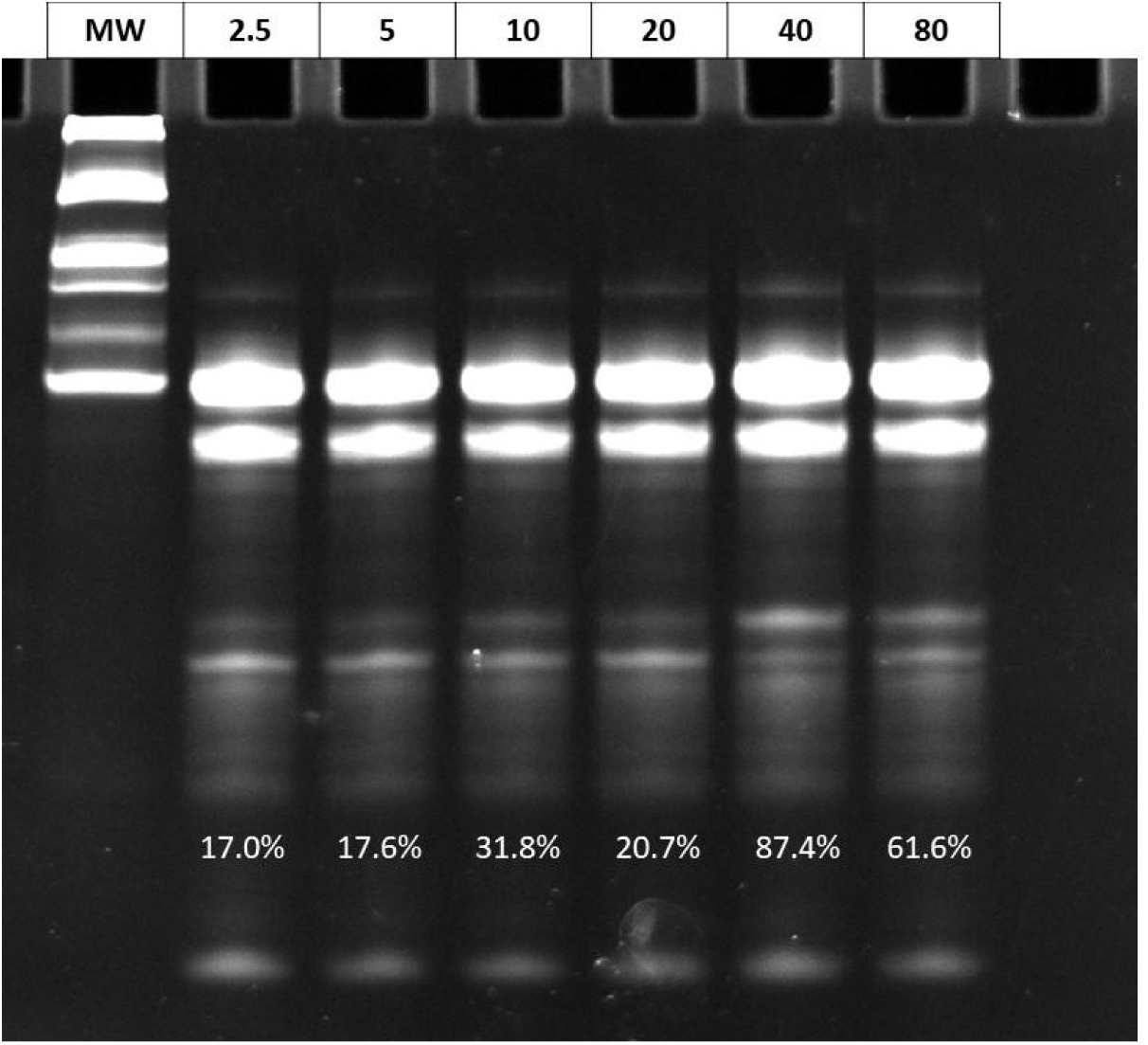
Magnesium titration in eutectic flexizyme reactions. Magnesium was titrated from 2.5 mM to 80 mM and the eutectic reactions were incubated for 6 hours at -20 C.

**Figure S73.**
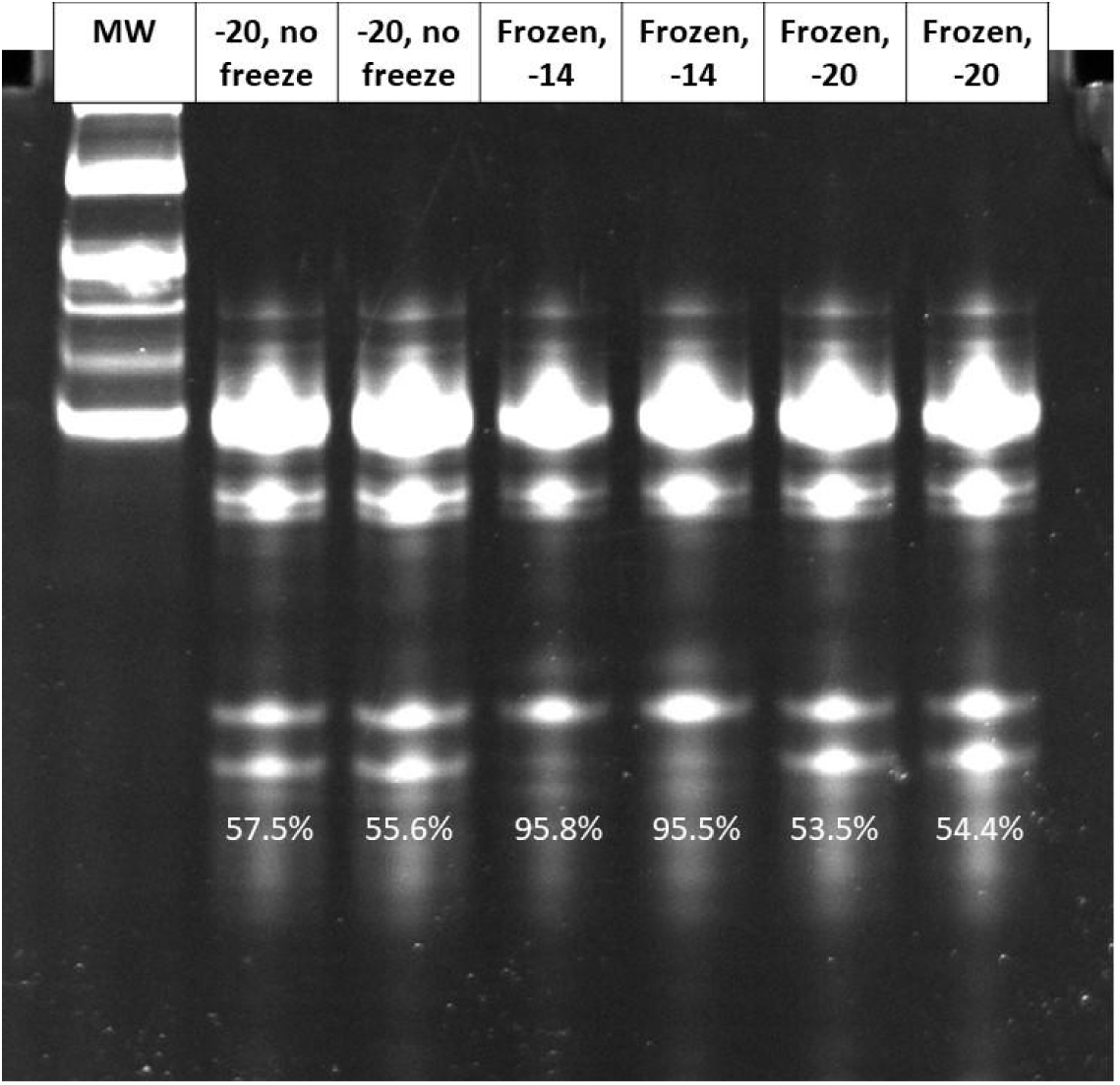
A gel comparing different eutectic reaction conditions using pCNF-CME: with or without snap freezing and -14 vs -20 C

**Figure S74.**
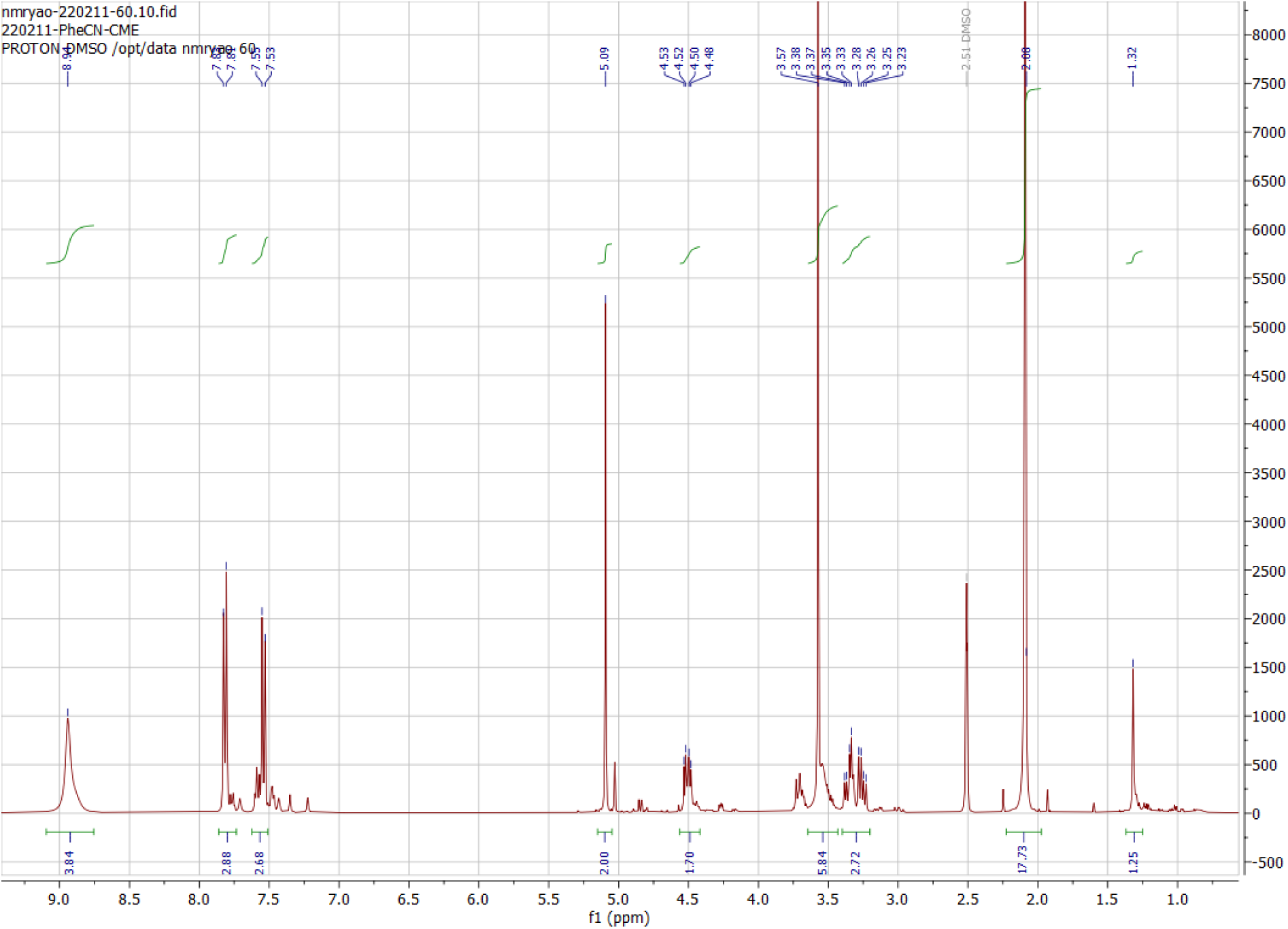
H NMR of 4-cyano-phenylalanine-CME

**Figure S75.**
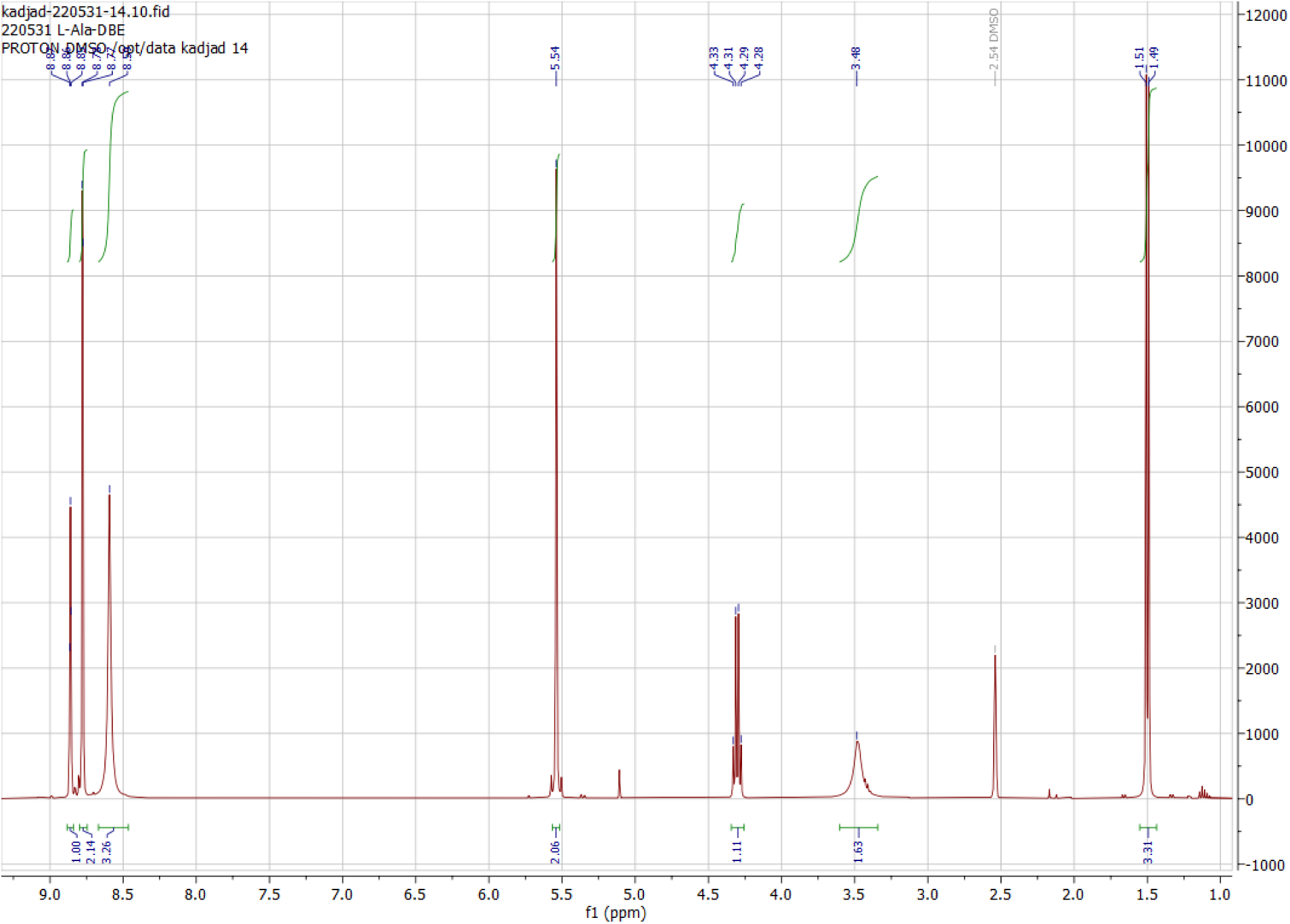
H NMR of L-alanine-DBE

**Figure S76.**
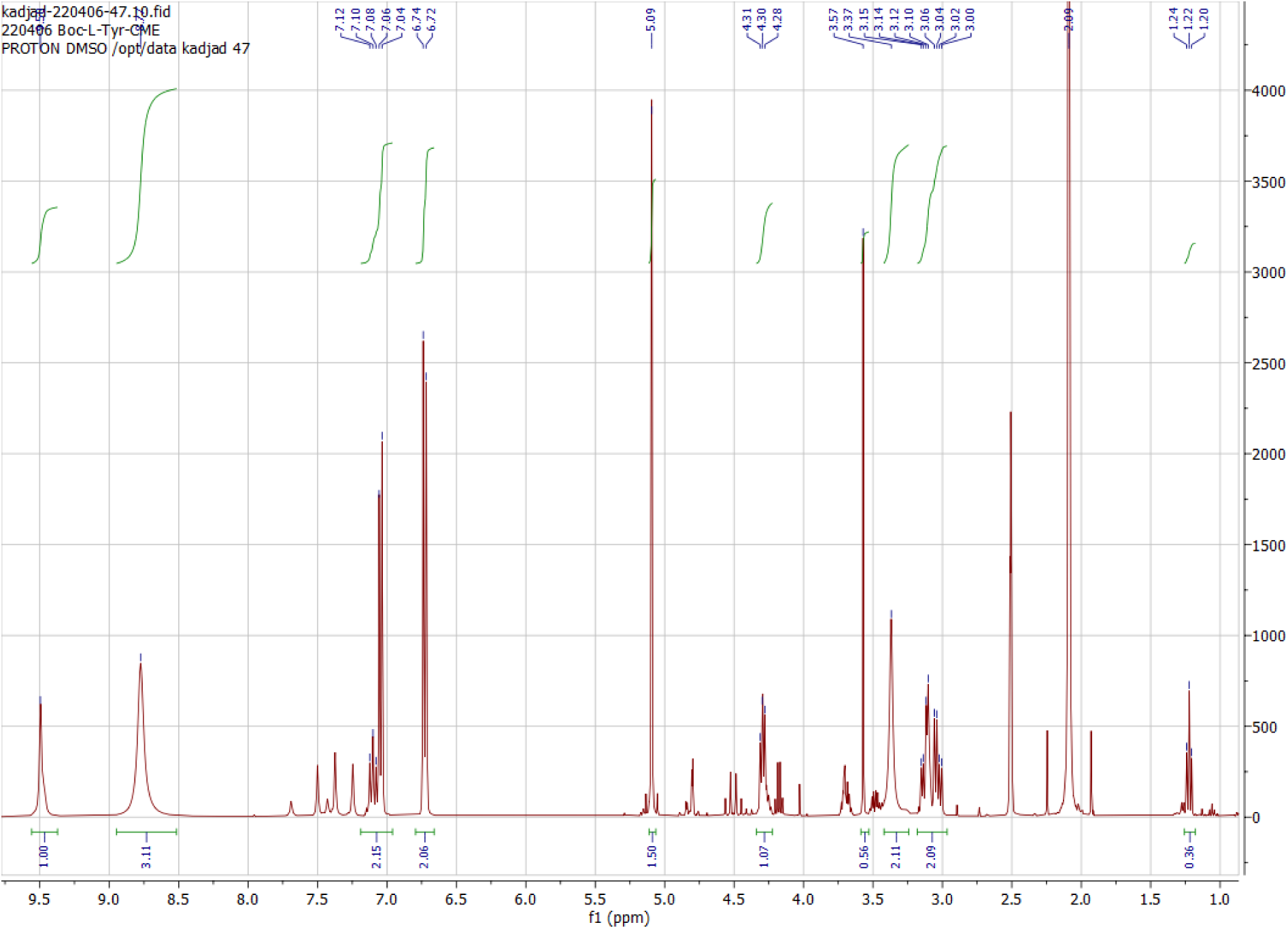
H NMR of L-tyrosine-CME

**Figure S77.**
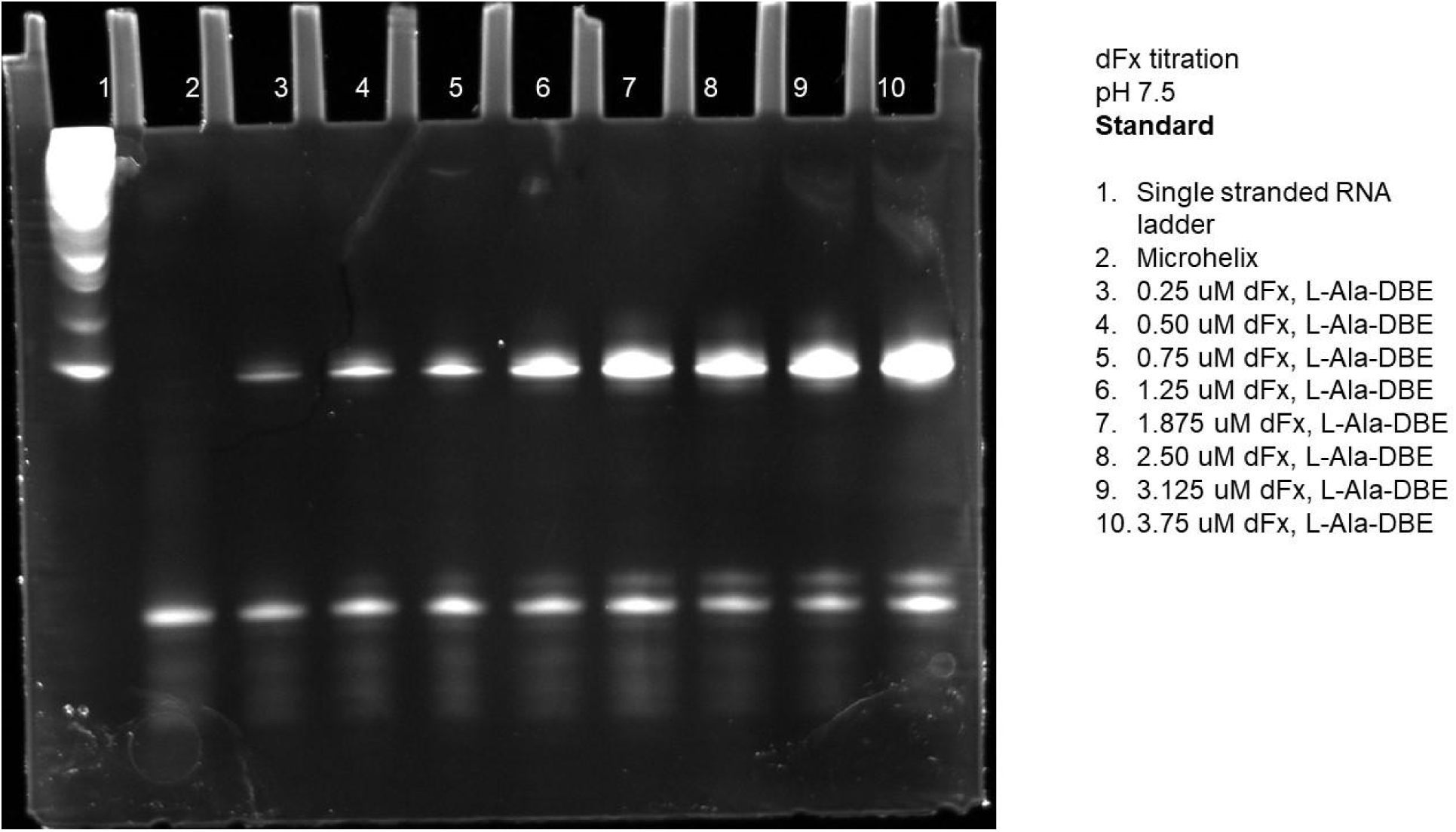
Gel of standard dFx titration at pH 7.5 using L-Ala-DBE.

**Figure S78.**
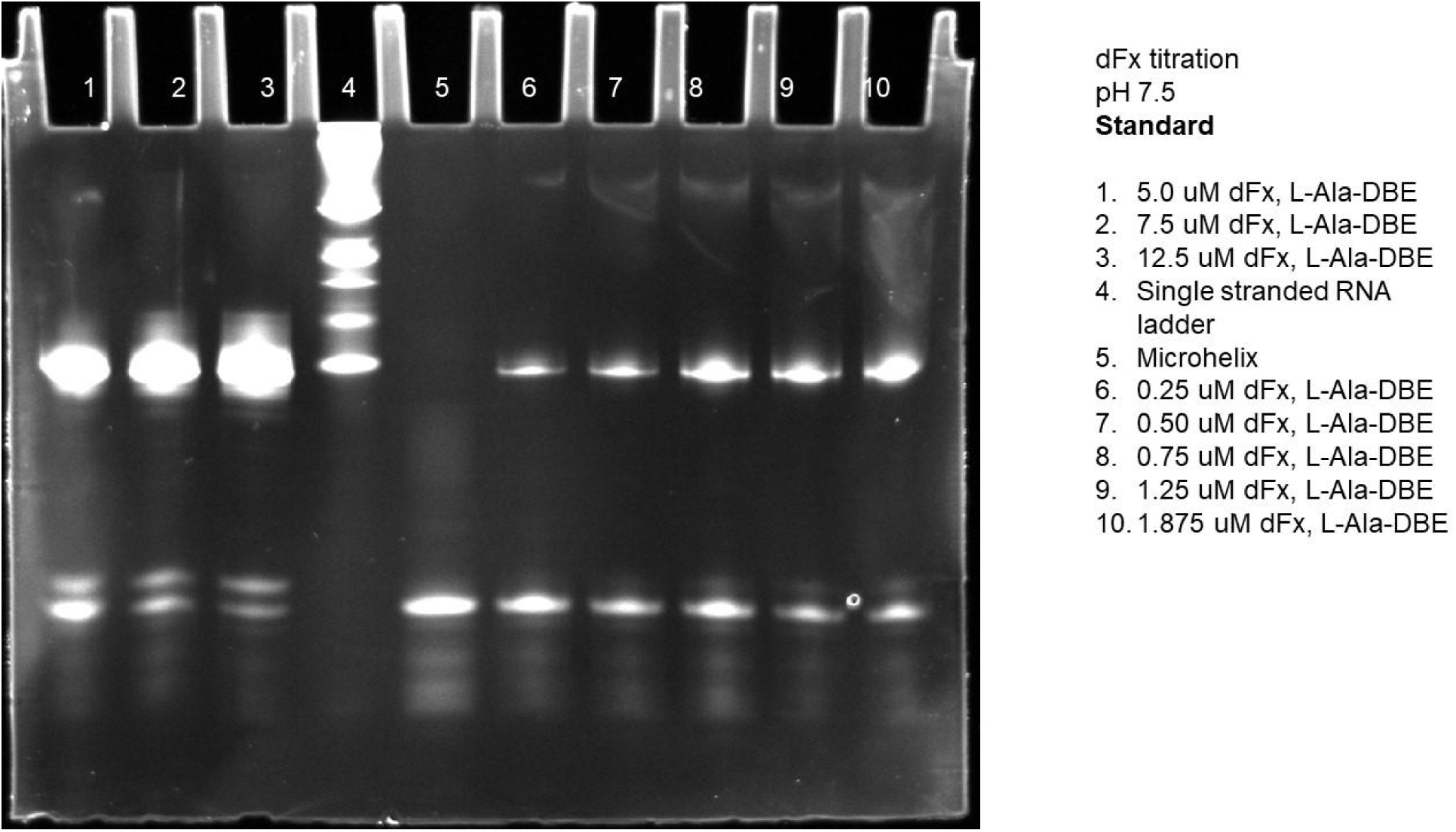
Gel of standard dFx titration at pH 7.5 using L-Ala-DBE.

**Figure S79.**
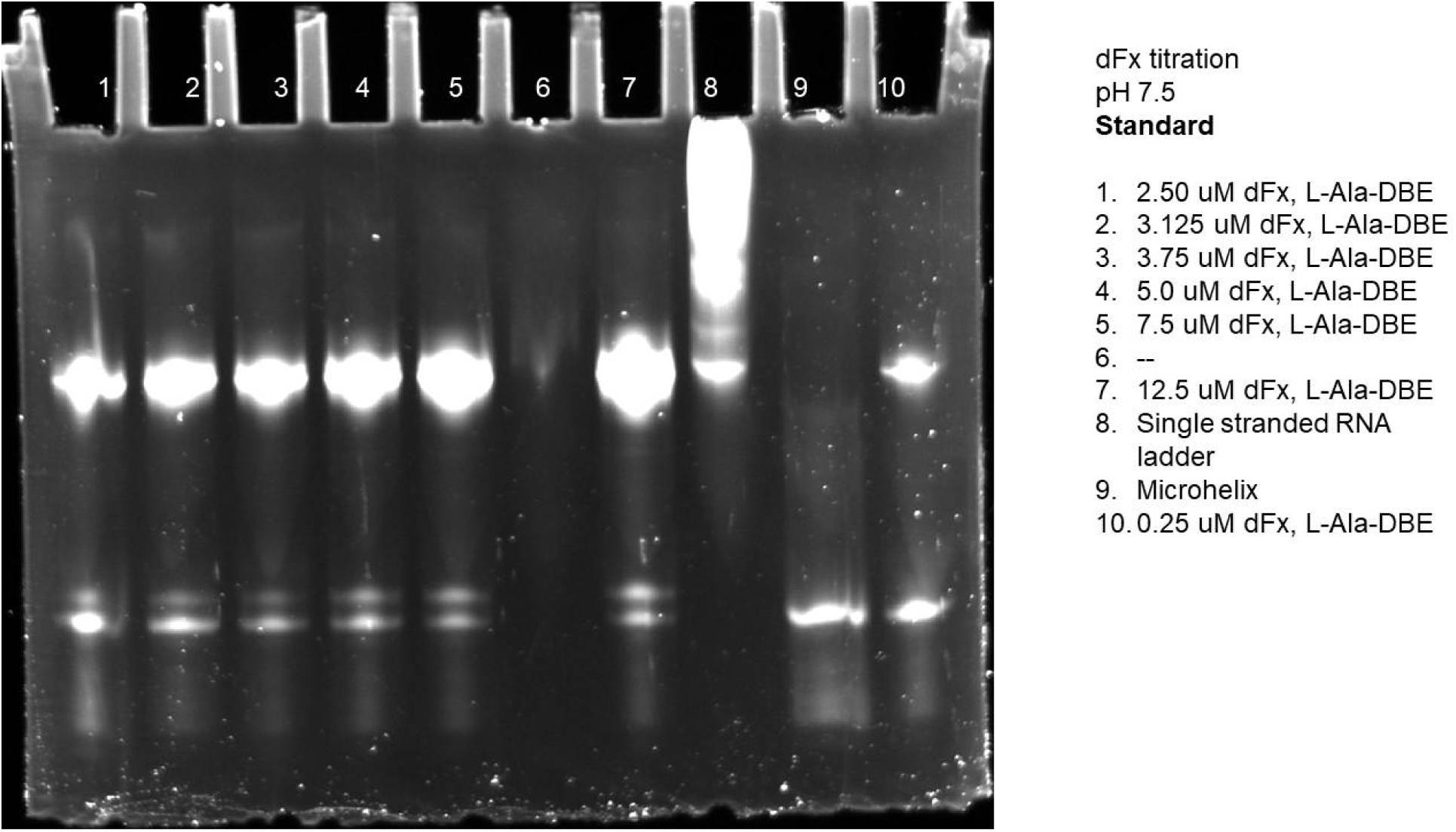
Gel of standard dFx titration at pH 7.5 using L-Ala-DBE.

**Figure S80.**
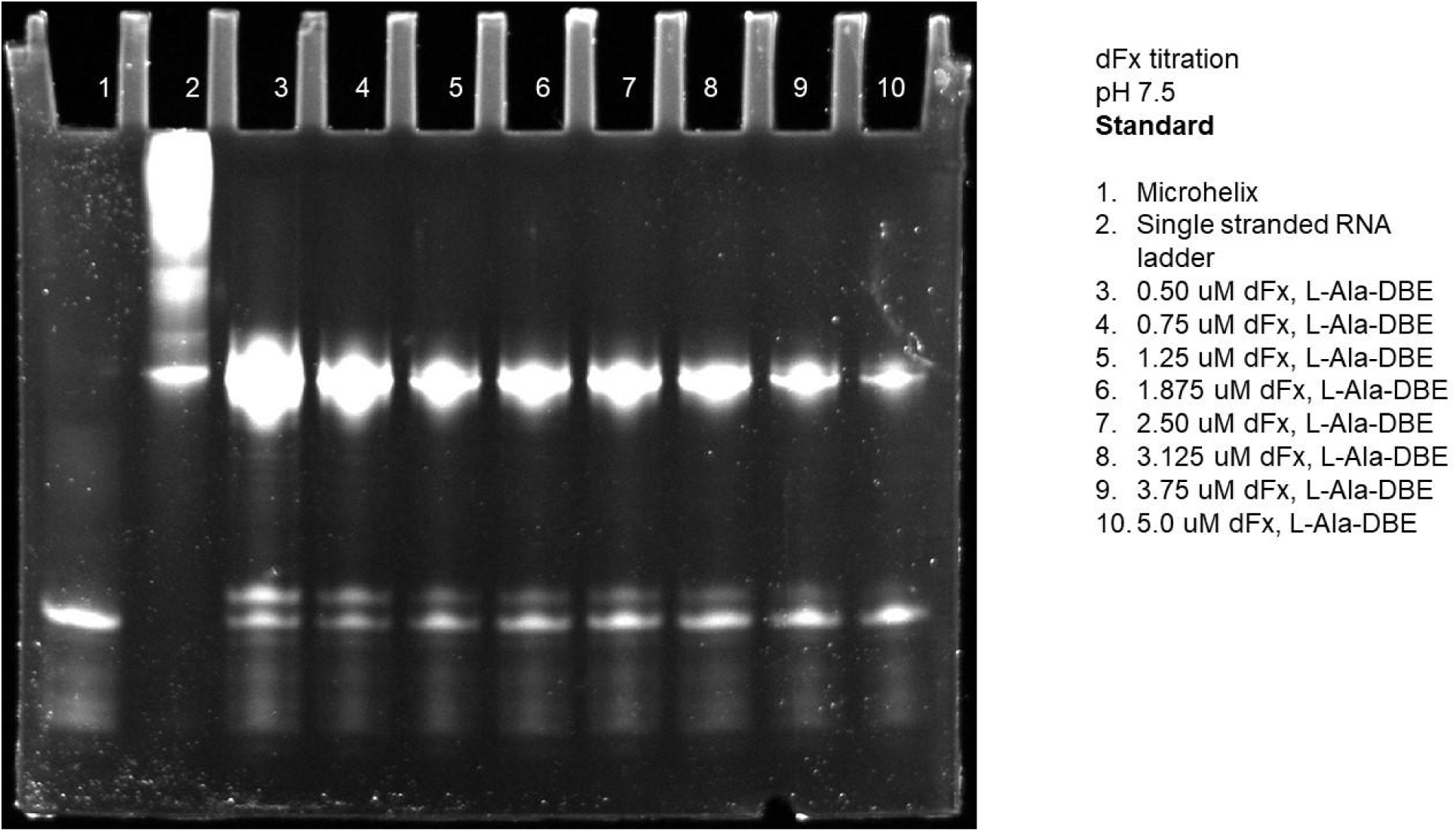
Gel of standard dFx titration at pH 7.5 using L-Ala-DBE.

**Figure S84.**
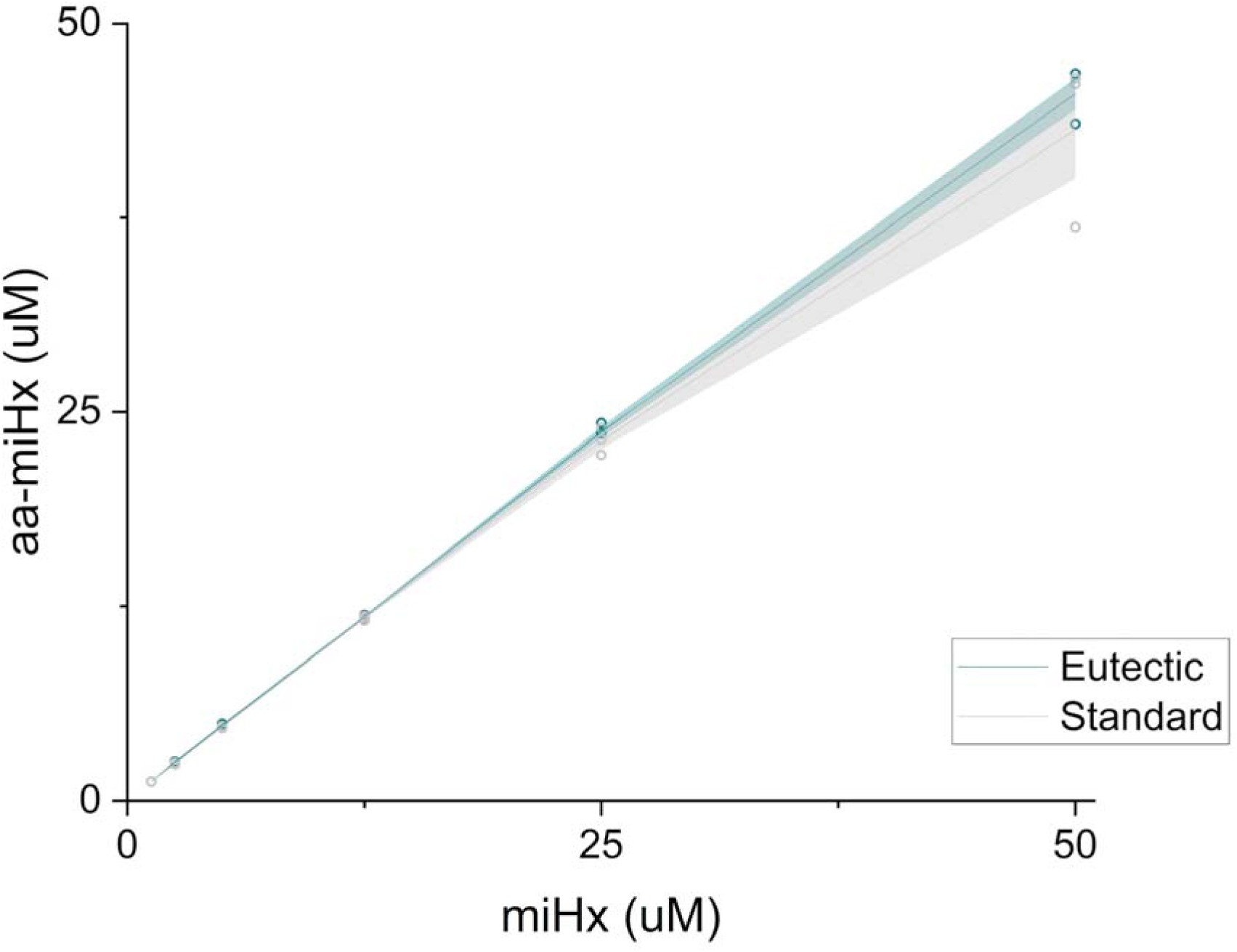
Eutectic and standard miHx titration for eFx with L-Tyr-CME using gels from supplementary figures S53-58.

**Figure S85.**
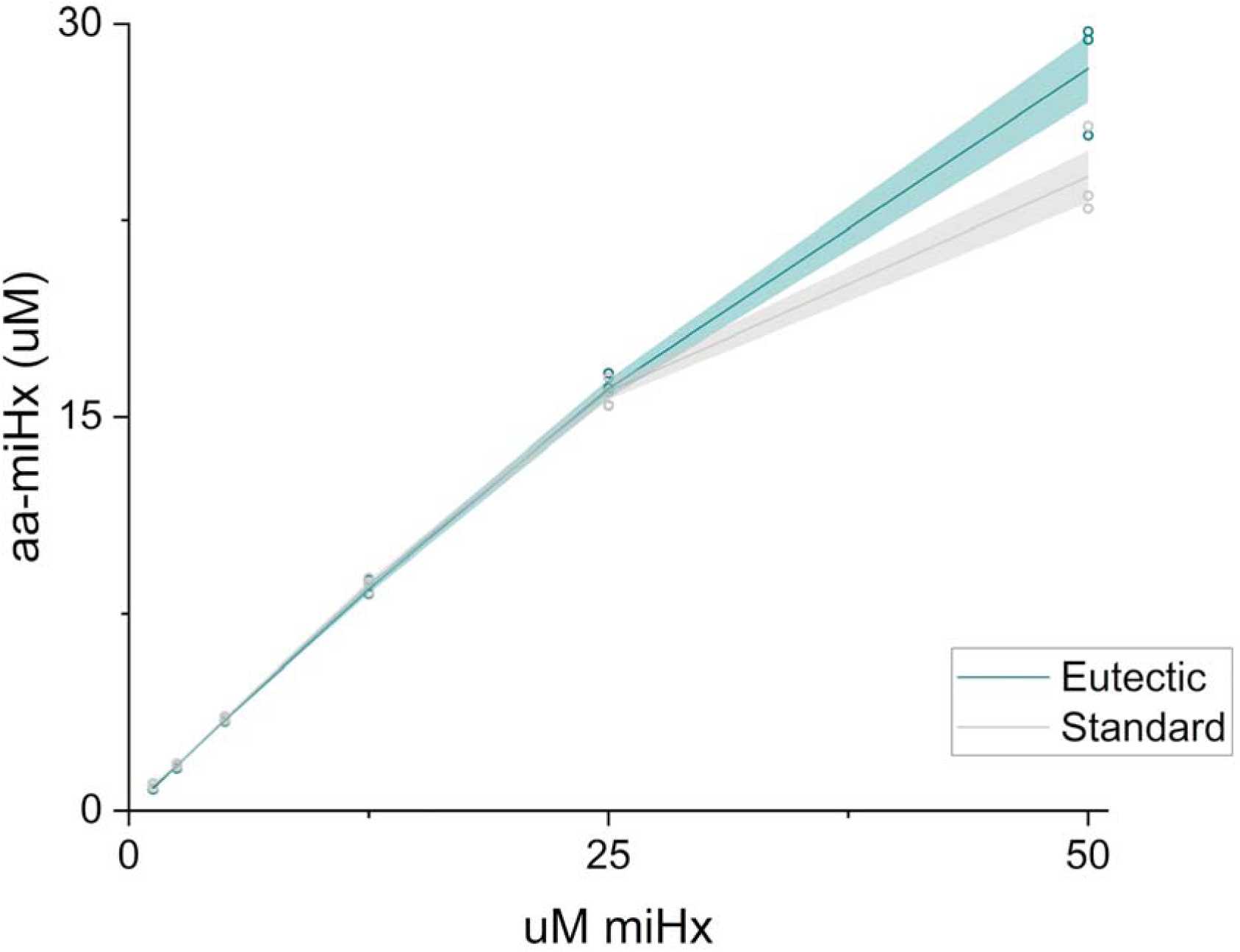
Eutectic and standard miHx titration for dFx with L-Ala-DBE using gels from supplementary figures S59-64.

**Figure S86.**
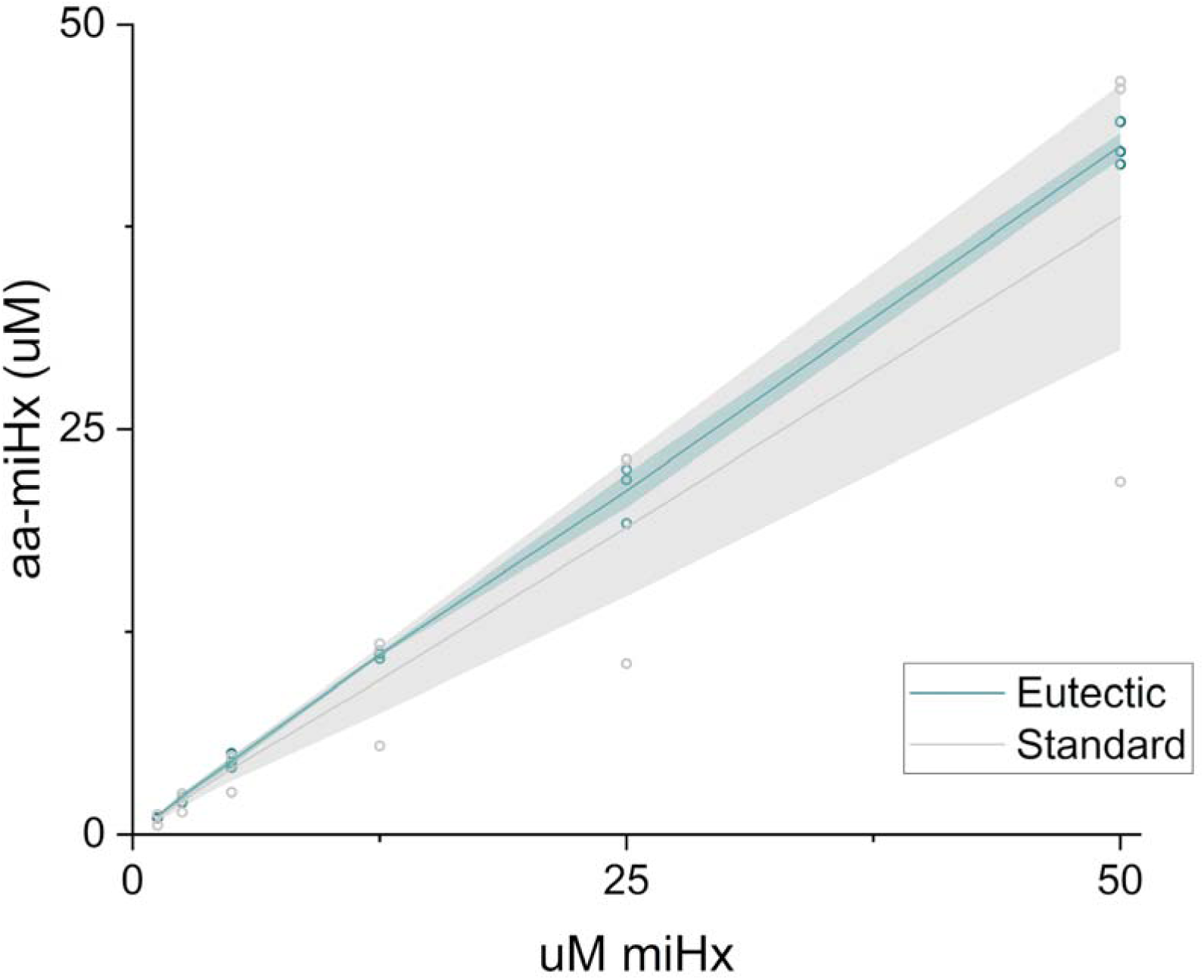
Eutectic and standard miHx titration for aFx with L-Leu-ABT using gels from supplementary figures S65-70.

**Figure S87.**
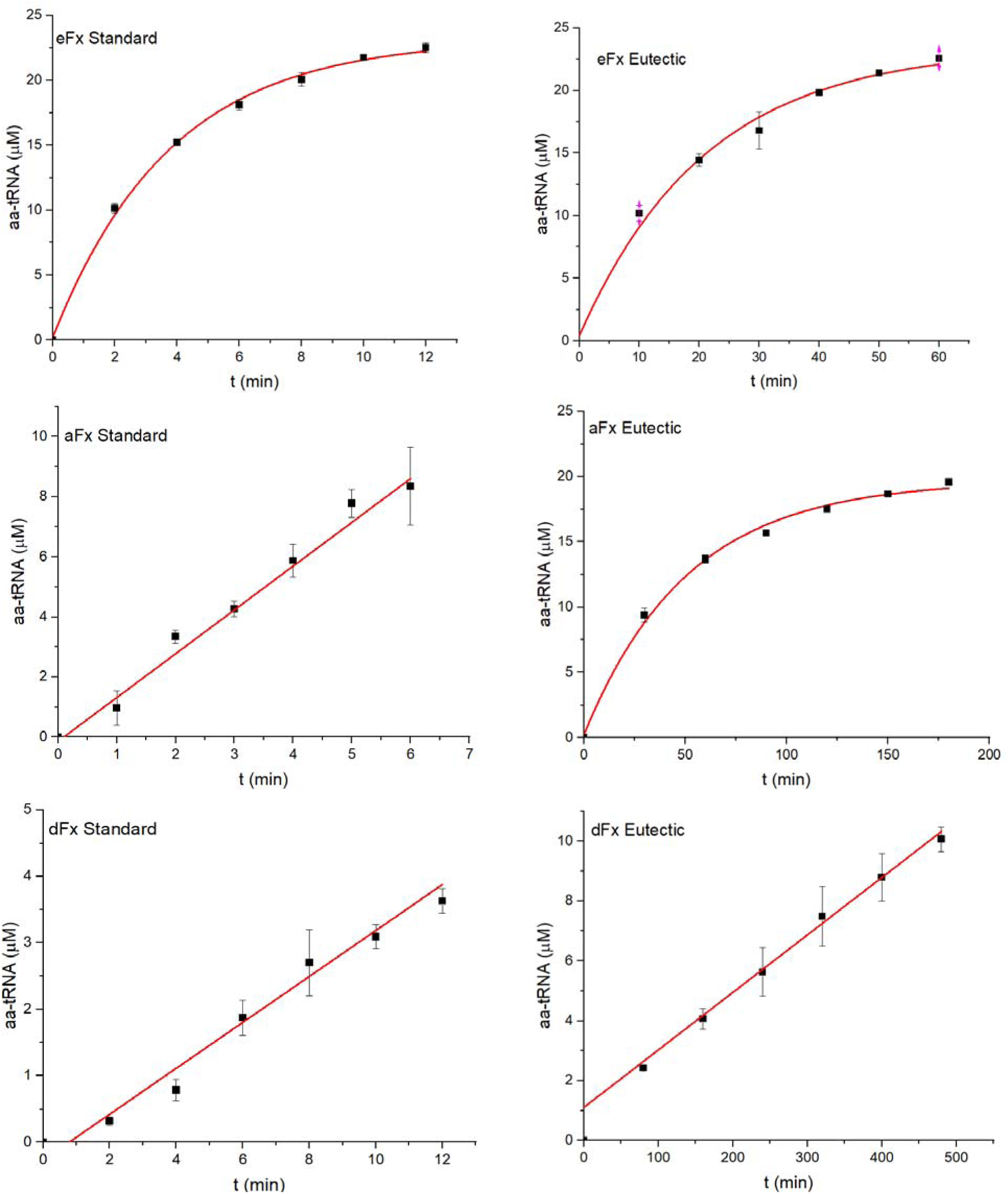
Short aminoacylation time courses used to calculate aminoacylation rate in Figure 2k. Time course representative gels in figures S88.

**Figure S88.**
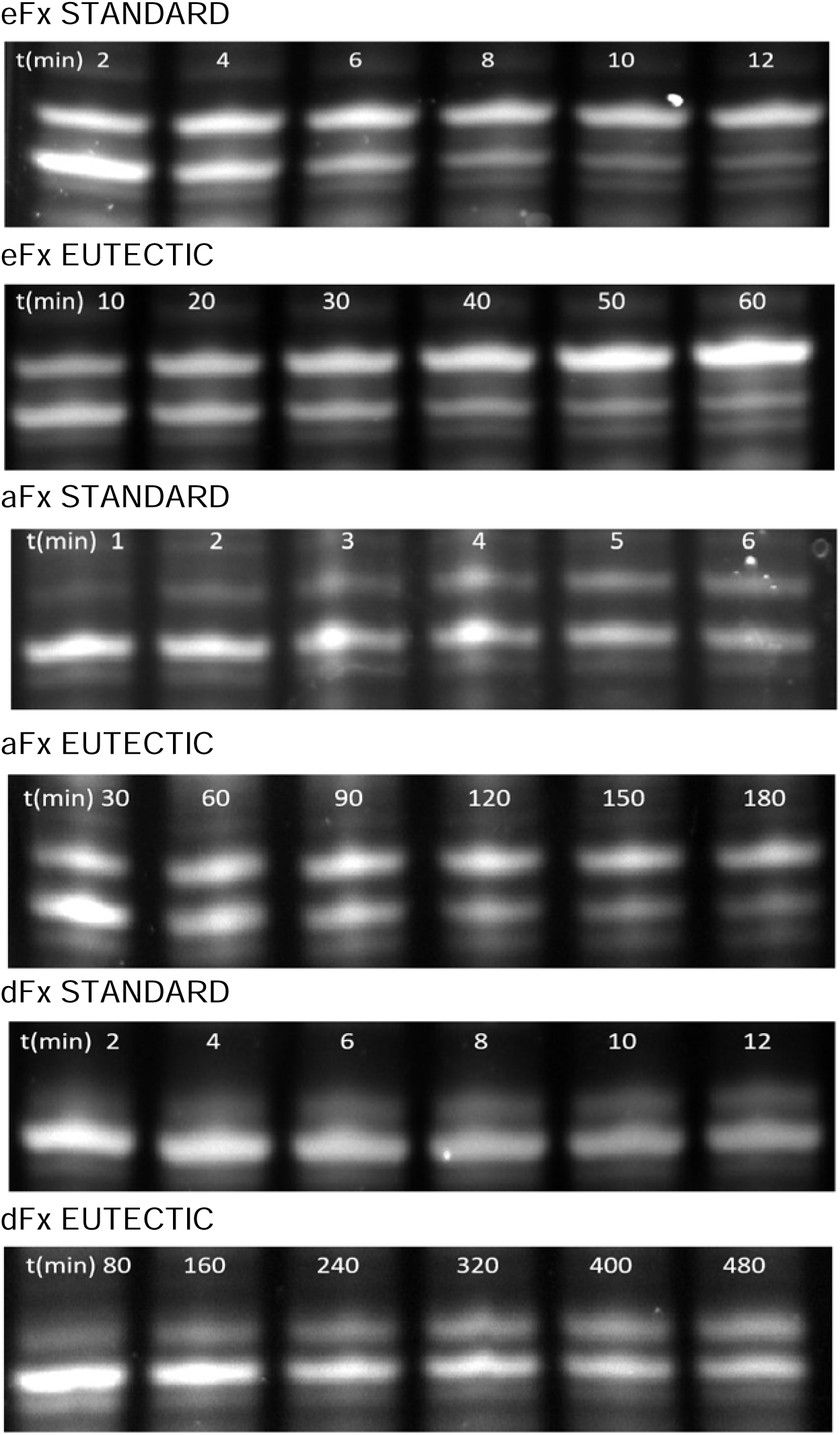
Gels showing short time course aminoacylation using to calculate aminoacylation rate of eutectic vs standard reaction conditions in Figure 2k and S87.

**Table S1.**
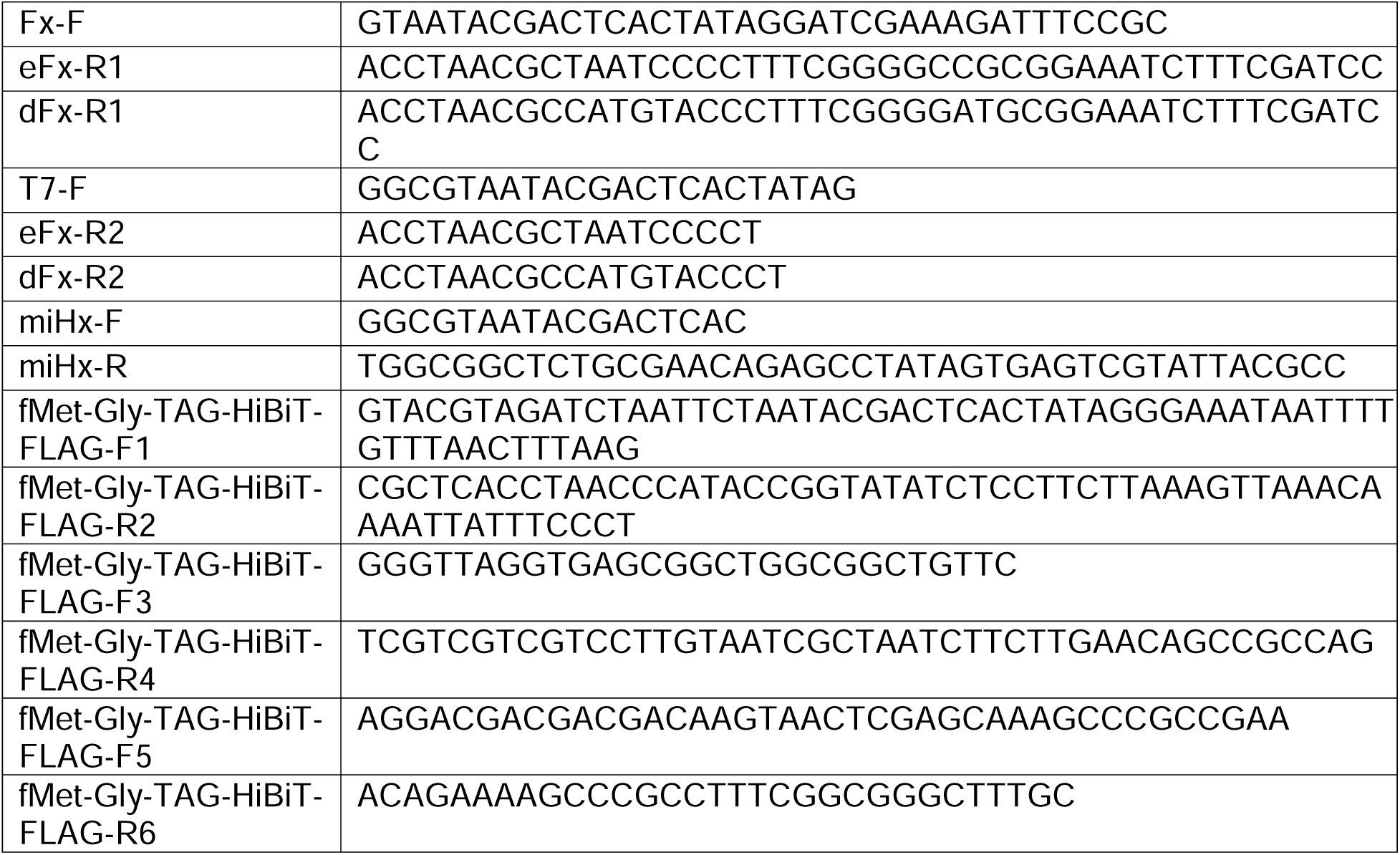
Sequences of oligonucleotides used to make DNA templates for flexizymes and HiBiT templates.

## Notes

### Competing Interest Statement

The authors have declared no competing interest.

## References

Ad, O., Hoffman, K. S., Cairns, A. G., Featherston, A. L., Miller, S. J., Söll, D., & Schepartz, A. (2019). Translation of Diverse Aramid- And 1,3-Dicarbonyl-peptides by Wild Type Ribosomes in Vitro. ACS Central Science, 5(7), 1289–1294. 10.1021/acscentsci.9b00460

Attwater, J., Wochner, A., Pinheiro, V. B., Coulson, A., & Holliger, P. (2010). Ice as a protocellular medium for RNA replication. Nature Communications, 1(6). 10.1038/ncomms1076

Chen, J., Chen, M., & Zhu, T. F. (2021). Translating protein enzymes without aminoacyl-tRNA synthetases. Chem, 7(3), 786–798. 10.1016/j.chempr.2021.01.017

Deich, C., Cash, B., Sato, W., Sharon, J., Aufdembrink, L., Gaut, N. J., Heili, J., Stokes, K., Engelhart, A. E., & Adamala, K. P. (2023). T7Max transcription system. Journal of Biological Engineering, 17(1). 10.1186/s13036-023-00323-1

Dixon, A. S., Schwinn, M. K., Hall, M. P., Zimmerman, K., Otto, P., Lubben, T. H., Butler, B. L., Binkowski, B. F., MacHleidt, T., Kirkland, T. A., Wood, M. G., Eggers, C. T., Encell, L. P., & Wood, K. V. (2016). NanoLuc Complementation Reporter Optimized for Accurate Measurement of Protein Interactions in Cells. ACS Chemical Biology, 11(2), 400–408. 10.1021/acschembio.5b00753

Goto, Y., Katoh, T., & Suga, H. (2011). Flexizymes for genetic code reprogramming. Nature Protocols, 6(6), 779–790. 10.1038/nprot.2011.331

Katoh, T., Iwane, Y., & Suga, H. (2017). Logical engineering of D-arm and T-stem of tRNA that enhances D-amino acid incorporation. Nucleic Acids Research, 45(22), 12601–12610. 10.1093/nar/gkx1129

Katoh, T., & Suga, H. (2020). Ribosomal elongation of aminobenzoic acid derivatives. Journal of the American Chemical Society, 142(39), 16518–16522. 10.1021/jacs.0c05765

Katoh, T., & Suga, H. (2022). Annual Review of Biochemistry In Vitro Genetic Code Reprogramming for the Expansion of Usable Noncanonical Amino Acids. 10.1146/annurev-biochem-040320

Lee, J., Coronado, J. N., Cho, N., Lim, J., Hosford, B. M., Seo, S., Kim, D. S., Kofman, C., Moore, J. S., Ellington, A. D., Anslyn, E. V., & Jewett, M. C. (2022). Ribosome-mediated biosynthesis of pyridazinone oligomers in vitro. Nature Communications, 13(1). 10.1038/s41467-022-33701-2

Monnard, P. A., Kanavarioti, A., & Deamer, D. W. (2003). Eutectic Phase Polymerization of Activated Ribonucleotide Mixtures Yields Quasi-Equimolar Incorporation of Purine and Pyrimidine Nucleobases. Journal of the American Chemical Society, 125(45), 13734–13740. 10.1021/ja036465h

Murakami, H., Ohta, A., Ashigai, H., & Suga, H. (2006). A highly flexible tRNA acylation method for non-natural polypeptide synthesis. Nature Methods, 3(5), 357–359. 10.1038/nmeth877

Niwa, N., Yamagishi, Y., Murakami, H., & Suga, H. (2009). A flexizyme that selectively charges amino acids activated by a water-friendly leaving group. Bioorganic and Medicinal Chemistry Letters, 19(14), 3892–3894. 10.1016/j.bmcl.2009.03.114

Passioura, T., Liu, W., Dunkelmann, D., Higuchi, T., & Suga, H. (2018). Display Selection of Exotic Macrocyclic Peptides Expressed under a Radically Reprogrammed 23 Amino Acid Genetic Code. Journal of the American Chemical Society, 140(37), 11551–11555. 10.1021/jacs.8b03367

Robertson, S. A., Ellman, J. A., & Schultz, P. G. (1991). A General and Efficient Route for Chemical Aminoacylation of Transfer RNAs. In J. Am. Chem. Soc (Vol. 113). https://pubs.acs.org/sharingguidelines

